# Predicted structure of the hepatitis B virus polymerase reveals an ancient conserved protein fold

**DOI:** 10.1101/2022.02.16.480762

**Authors:** Razia Tajwar, Daniel P. Bradley, Nathan L. Ponzar, John E. Tavis

## Abstract

Hepatitis B virus (HBV) replicates by protein-primed reverse transcription. It chronically infects >250 million people, and the dominant anti-HBV drugs are nucleos(t)ide analogs targeting the viral polymerase (P). P has four domains, the terminal protein (TP) that primes DNA synthesis, a spacer, the reverse transcriptase (RT), and the ribonuclease H (RNaseH). Despite being a major drug target and catalyzing a reverse transcription pathway very different from the retroviral pathway, HBV P has resisted structural analysis for decades. Here, we exploited advances in protein structure prediction to model the structure of P. The predicted HBV RT and RNaseH domains aligned to the HIV RT-RNaseH at 3.75 Å RMSD, had a nucleic acid binding groove spanning the two active sites, had DNA polymerase active site motifs in their expected positions, and accommodated two Mg^++^ ions in both active sites. Surprisingly, the TP domain wrapped around the RT domain, with the priming tyrosine poised over the RT active site. This model was validated using published mutational analyses, and by docking RT and RNaseH inhibitors, RNA:DNA heteroduplexes, and the HBV ε RNA stem-loop that templates DNA priming into the model. The HBV P fold, including the orientation of the TP domain, was conserved among hepadnaviruses from rodents to fish and in P from a fish nackednavirus, but not in other non-retroviral RTs. Therefore, this protein fold has persisted since the hepadnaviruses diverged from nackednaviruses >400 million years ago. This model will guide drug development and mechanistic studies into P’s function.

**Significance:** The hepatitis B virus (HBV) polymerase (P) catalyzes an unusual reverse transcription pathway and is a major drug target. However, P’s insolubility and instability have long prevented its structural analyses. This work predicts the structure of P proteins from human to fish viruses, revealing an unanticipated conserved protein fold that is at least 400 million years old. The HBV P model was validated by mechanistically interpreting mutations with strong phenotypes and by computationally docking nucleic acids to P and inhibitors into both of its active sites. The model will advance mechanistic studies into the function of P and enable drug discovery against targets on P other than the reverse transcriptase and ribonuclease H active sites.

## Introduction

Hepatitis B virus (HBV) is the human member of the *Hepadnaviridae* virus family. It is a small, enveloped, partially double-stranded DNA virus that replicates by reverse transcription in hepatocytes (1). HBV is genetically diverse, with nine genotypes differing by >8% at the sequence level (2). Over 250 million people are chronically infected with the virus (3), leading to >850,000 deaths annually through induction of liver failure and hepatocellular carcinoma (4). The dominant treatments for HBV infection employ nucleos(t)ide analog drugs that suppress reverse transcription, but treatment is not curative and is life-long for the majority of patients (5, 6).

Hepadnaviral reverse transcription (7, 8) occurs within cytoplasmic capsid particles and is catalyzed by the viral polymerase protein (termed “P”). Reverse transcription begins with binding of P to the ε RNA stem-loop on the viral pregenomic RNA (pgRNA) in an HSP90-mediated reaction (9, 10) that involves the T3 and RT1 motifs of P (11). The reverse transcriptase (RT) activity of P initiates DNA synthesis using Y63 in its N-terminal domain as a primer and a bulge in ε as the template (12), covalently linking P to the viral minus-polarity DNA strand (i.e., the first strand synthesized). The viral ribonuclease H (RNaseH) activity degrades the pgRNA after it has been copied into minus-polarity DNA. Studies with duck hepatitis B Virus (DHBV), a distant homolog of HBV that shares the same reverse transcription pathway and domain structure of P, reveal that the RNaseH leaves a 15-18 nt capped RNA due to the distance between the RT and RNaseH active sites (13) that primes plus-polarity DNA strand synthesis. Reverse transcription employs three strand transfers that are presumably promoted by P to make the mature partially double-stranded viral DNA genome.

P has four domains defined by homology alignments (Fig. 1A). The terminal protein (TP) domain contains about 180 amino acids (aa), including Y63 that primes DNA synthesis. This is followed by the poorly conserved and putatively unstructured ∼ 175 aa spacer domain with little known function that accommodates the viral pre-S1 surface glycoprotein sequences encoded in an overlapping reading frame (1). The spacer is followed by the catalytic core of the enzyme comprised of the RT and RNaseH domains. The RT domain (∼335 aa) carries the canonical A-E reverse transcription motifs (14), including the YMDD motif that chelates two Mg^++^ ions essential for DNA synthesis. Finally, the RNaseH domain is about 155 aa long and contains the D-E-D-D motif that chelates two Mg^++^ ions essential for RNA hydrolysis (15–17).

**Figure 1.**
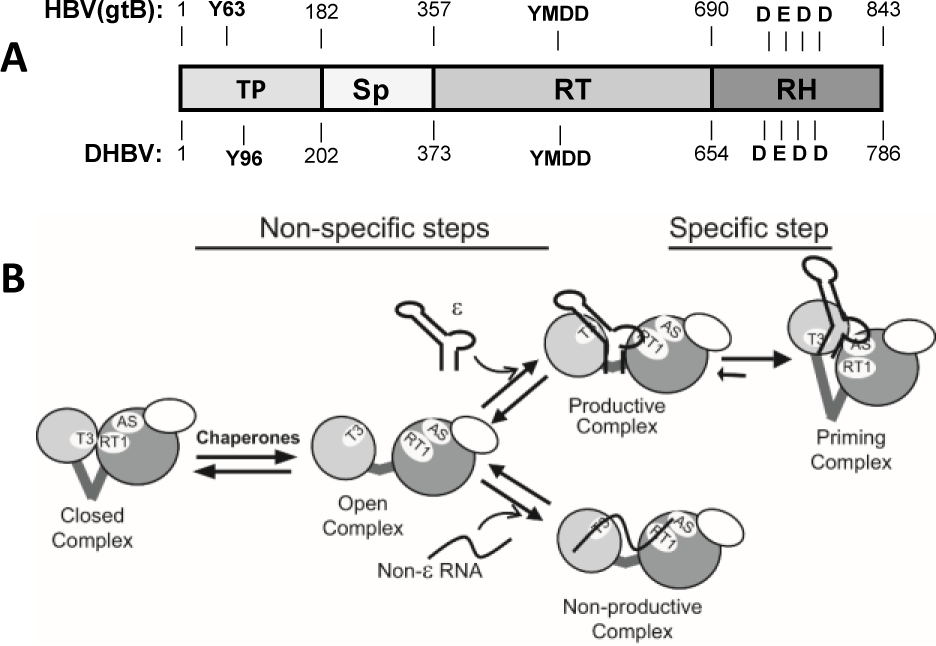
Gene organization for HBV P and conformations adopted by the enzyme. **A.** Gene organization. Genotype B numbers are used for the domain boundaries, and the Y63 priming residue, YMDD RT active site motif, and D-E-D-D RNaseH active site motif are indicated. **B.** Conformations adopted by P identified by prior analyses. The TP domain is in light gray, the RT domain in dark gray, the RNaseH domain as an unshaded oval, and the spacer domain is indicted by thick lines. AS, RT active site; T3, T3 motif in the TP domain; RT1, RT1 motif in the RT domain; ε, the HBV ε stem loop needed for specific RNA encapsidation and priming. Reprinted with permission from (11).

The extreme C-terminal ∼35 residues of P are dispensable for RNaseH activity *in vitro* (18) and are presumed to be unstructured. P is 843 aa long in most HBV’s genotypes, with P from the various genotypes sharing 86-88% identity (HBV genotype B numbering will be used unless indicated otherwise). Genotype A has a two-residue insertion between positions 16 and 17 in the terminal protein, genotype D lacks 11 aa in the spacer domain relative to the other genotypes (residues 184-194), and genotypes E and G lack amino acid 184 in the spacer region.

P is a monomer throughout viral DNA synthesis and remains attached to the viral DNA within mature virions (19), mandating flexibility in the enzyme. As expected from its attachment to DNA, up to five conformations have been identified for P through molecular biology and biochemical analyses, primarily using DHBV (Fig. 1B) (11, 20–23).

No experimentally determined structures exist for any domain of P despite its importance as a drug target and the unusual reverse transcription pathway it catalyzes due to protein production issues and P’s conformational plasticity. An *ab initio* predicted model has been proposed for the TP domain based on secondary structure predictions (24). Multiple homology models generated against retroviral RT domains exist for the RT domain of HBV P (25, 26).

These models display the right-hand shape of a DNA polymerase and are accurate enough in the active site to permit interpretation of the mechanisms of resistance to nucleoside analog drugs. Three homology models exist for the RNaseH domain (27–29). They have most of the established RNaseH fold (30) but their C-termini disagree, with only the Li et al. model (27) correctly predicting the last D-E-D-D residue (15, 31).

AlphaFold is an *ab initio* protein structure prediction program developed by DeepMind (32). It uses correlated sequence variations among homologous sequences to define intra-protein distance constraints. These constraints, plus secondary and tertiary structural information from homologous protein sequences when available, are iteratively combined to predict the structure of the α-carbon backbone of a protein. Side chain orientations are then refined by energy minimization. AlphaFold predicted the backbone structure of all 87 protein domains in the 14th Critical Assessment of Protein Structure Prediction (CASP14) with 0.96 Å root mean square deviation (RMSD) for the CASP domains compared to the experimentally determined structures; accuracy including the side chains was 1.5 Å RMSD. Structures were also predicted by AlphaFold for 10,795 proteins whose structures were previously determined, with > 50% of the predicted structures having α-carbon chain RMSD values ≤ 2 Å compared to the experimental results (32). Accuracy of the AlphaFold predictions is lower for protein regions with about 30 or fewer homologous sequences from which to derive distance constraints, with accuracy plateauing around 100 homologous sequences (32). Accuracy of the predictions is based in part on the lowest distance difference test (lDDT) that computes the accuracy of the local structure around each amino acid, allowing accuracy assessment for different sections of the protein model with little contribution from domain packing (33). The predicted lDDT scores (pLDDT) generated by AlphaFold correlate well with the actual lDDT scores (32). These studies validate AlphaFold’s use in structural prediction, with the understanding that the predicted structures need to be experimentally evaluated.

Here, we predicted the structure of HBV P for each of its nine genotypes and validated the fold using data from 30 years of mutational analyses of P. Structures of P proteins from animal hepadnaviruses and nackednaviruses with hosts ranging from woodchucks to fish were predicted and compared to the HBV P models. These studies reveal the overall fold of the enzyme for the first time and permitted approximation of its antiquity.

## Results and Discussion

### Modeling

The initial set of HBV P models were generated using sequences from genotype B (Genbank AB554017). Models were generated for each domain independently, the catalytic core of the enzyme containing the RT and RNaseH domains, and the full-length enzyme using AlphaFold. Models with the highest pLDDT scores were energy-minimized using the Amber-Relax module and used for all analyses. The domain boundaries employed correspond to those used by Donlin et al. (34), with the N-terminus of the RT domain moved to residue 357 (genotype B numbering) to include a β-sheet that is conserved at the boundary of the spacer and RT domains in all models. This is residue rt-12 in the RT domain nomenclature used for assessing nucleos(t)ide analog resistance. The individual domain models are in Figs. S1-S4.

### Catalytic core model

A two-domain model of the HBV catalytic core containing the RT and RNaseH domains folded with high confidence (pLDDT > 80) for the bulk of the domain, with lower confidence at the N-terminus of the RT domain that is adjacent to the spacer domain, ∼80 aa in the center of the domain that includes the YMDD motif that chelates the Mg^++^ ions, and the extreme C-terminus (Fig. S5).

The HBV RT domain adopted the right-hand shape of a DNA polymerase with fingers, palm, and thumb subdomains (Fig. 2A). The conserved A-E DNA polymerase motifs (14) were in the palm subdomain, similar to their locations in other DNA polymerases. The model accommodated two Mg^++^ ions separated by 3.23 Å in the RT active site that were coordinated by the YMDD motif in positions analogous to those in the HIV RT enzyme. The RT domain in the two-domain model shared a 2.88 Å RMSD similarity with the isolated RT domain model (S24A). The RT domain within the predicted HBV catalytic core model could be superimposed with the RT domain model predicted by Das et al. (25) with an RMSD = 3.38 Å, with the major difference being that the AlphaFold structure predicted more α-helixes and β-sheets in the fingers subdomain (Fig. 3A). The HBV RT domain within the catalytic core model aligned to the HIV RT domain (PDB: 5XN1) with an RMSD = 3.34 Å (Fig.3B). The α-helixes and β-sheets in the fingers subdomain from the AlphaFold HBV RT model were more similar to the HIV structure than the Das et al. model.

**Figure 2.**
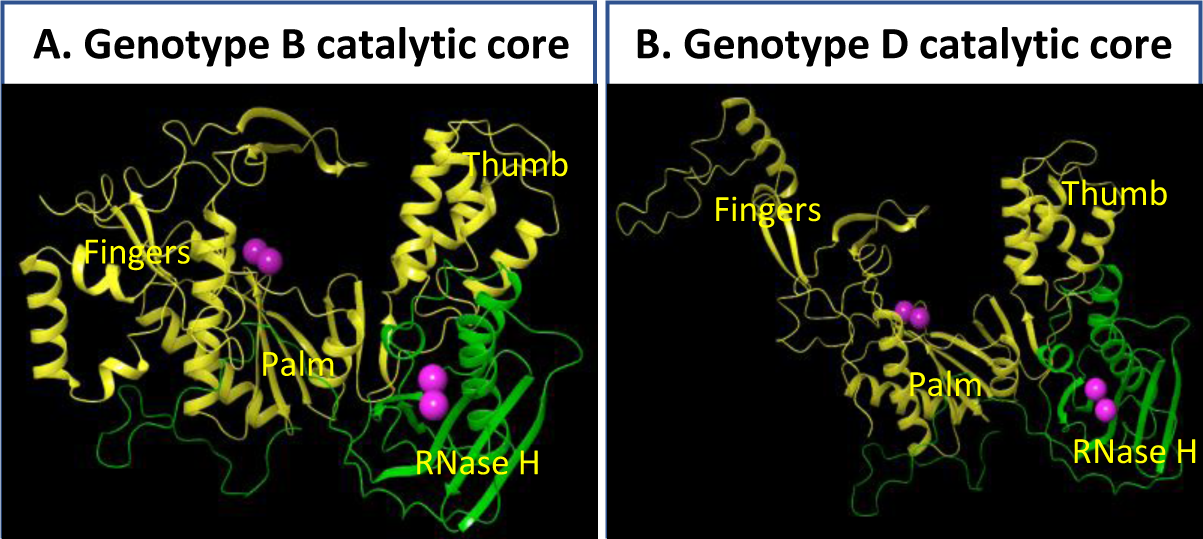
Models for the catalytic core of HBV P. **A.** Genotype B RT and RNaseH domains. **B.** Genotype D RT and RNaseH domains. Yellow, RT domain; Green, RNaseH domain; Magenta spheres, Mg^++^ ions.

**Figure 3.**
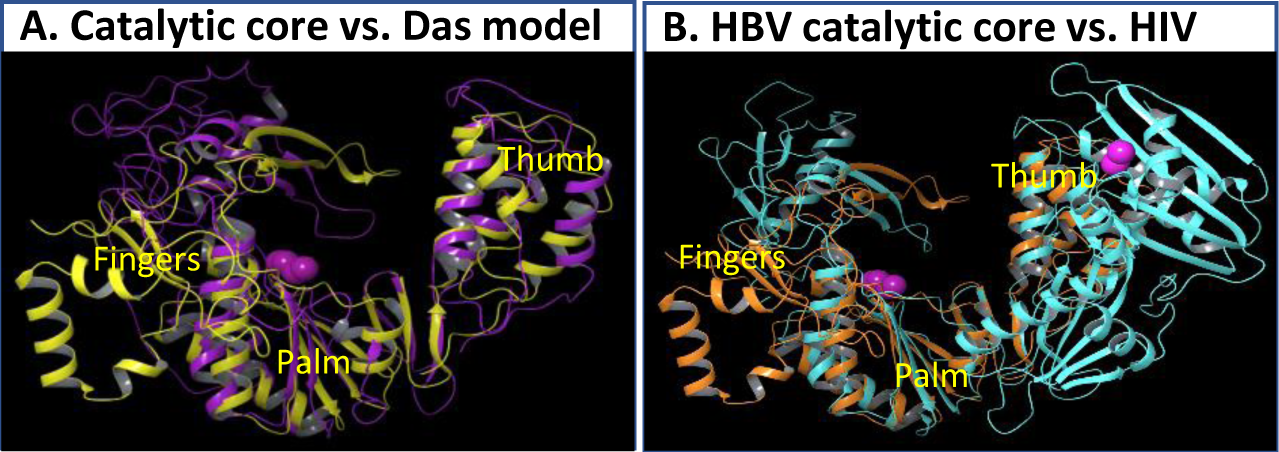
Superpositions of HBV P models. **A.** HBV genotype B RT domain from the catalytic core model (yellow) vs. the HBV RT domain model (purple) from Das et al. (25). **B.** HBV genotype B catalytic core model (orange) vs. HIV RT-RNaseH structure (cyan). Magenta spheres, Mg^++^ ions.

The RNaseH domain adopted the canonical RNaseH fold (16, 17) with a β-sheet platform overlaid by α-helices arranged in an “H” (Figs. S5 and S24B-D). The model accommodated two Mg^++^ ions 3.96 Å apart that were coordinated by the D-E-D-D motif that binds divalent cations in other RNases H, as expected because the HBV RNaseH requires Mg^++^ or Mn^++^ for activity and mutating the D-E-D-D residues yields an RNaseH-deficient phenotype (15, 31). The RNaseH domain from the catalytic core model aligned well with the isolated RNaseH domain model (RMSD = 1.38 Å; Fig. S24D), the HIV RNaseH (PDB: 3K2P, RMSD = 2.84 Å; Fig. S24B), and the human RNase H1 structure (PDB: 2QKK, RMSD = 2.66 Å; Fig. S24C). The AlphaFold model appears to be superior to our prior RNaseH model (27) because it predicted an α helix and a β sheet strand of the standard RNaseH fold (16, 17) that were previously poorly formed.

The relative orientation of the two domains in the HBV catalytic core model was similar to the HIV RT-RNaseH (Figs. 3B and S24F), and there was a plausible nucleic acid binding channel connecting the two catalytic domains. However, the catalytic core model did not align as well against the HIV RT-RNaseH (Fig. S24F) as its individual domains did when aligned separately in Figs. 3B and S24B because the HBV model lacked the linker region between the two domains found in the HIV RT-RNaseH enzyme.

We also predicted a two-domain model for HBV genotype D as it is a commonly used genotype for inhibitor screening (Fig. S6). The genotype B and D RT-RNaseH models aligned with an RMSD = 2.15 Å (Fig. S24E). The primary difference was a large variance in the orientation of part of the fingers subdomain that is implausible in the genotype B model because it did not resemble the orientation commonly seen in DNA polymerases. There were also small differences in the orientation of E729 within the D-E-D-D motif that slightly affected positioning of the RNaseH Mg^++^ ions.

### Full-length HBV polymerase model

Full-length P folded with high confidence (pLDDT > 80) with the exception of the extreme N-terminus, spacer domain, and the C-terminal ∼50 residues, and short regions of lesser confidence within each domain (Figs. 4A and S8). A full-length P model for genotype D was also predicted that aligned well with the genotype B model (RMSD = 1.96 Å) (Figs. S10 and S25C).

**Figure 4.**
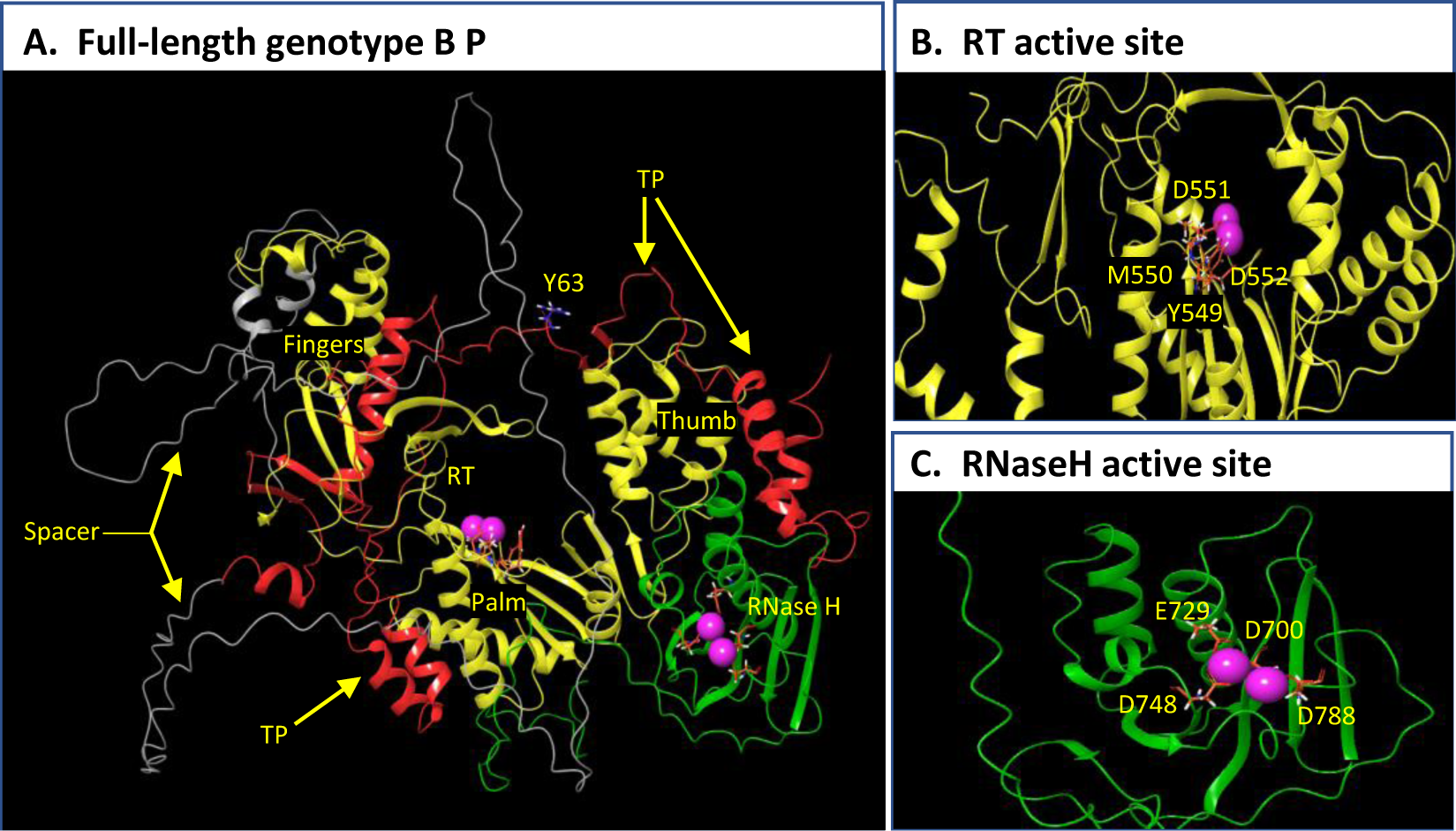
Predicted model of genotype B P model. **A.** Full-length P. **B.** RT active site. **C.** RNaseH active site. Red, TP domain; Gray, Spacer domain; Yellow, RT domain; Green, RNaseH domain. Magenta spheres, Mg^++^ ions. Y63 and the residues in the YMDD and D-E-D-D motifs are labeled and shown as sticks.

Intriguingly, the TP domain was projected to form a C-shaped structure that cups around the outside of the catalytic core of P (Fig. 4A). The TP domain was similar to the model predicted for the isolated TP domain (RMSD = 3.07 Å) (Figs. S1 and S26A). The Y63 residue in the TP domain that primes DNA synthesis and covalently links P to the viral minus-polarity DNA strand (12) was predicted to be on a loop located above the RT domain YMDD active site motif. Y63 is near the N-terminal edge of a region of high pLDDT confidence that was flanked by regions with pLDDT values of 50-60 that can imply poorly structured and/or flexible sequences. The putative flexibility of the loop is consistent with the need for conformational plasticity surrounding Y63 to permit protein priming in the RT active site followed by displacement of the priming loop from the active site by the growing DNA. It is also consistent with the conformational changes needed in the priming loop during the three DNA strand transfers that occur while P remains covalently attached to the 5’ end of the minus-polarity DNA strand via Y63.

The spacer domain was predicted to form an extended unstructured loop (Figs. 4A and S2) that was different in all five models generated for the domain by AlphaFold, consistent with its low sequence conservation. It is also consistent with the ability of the TP and RT domains to complement each other in *trans* as independent proteins (35–37) and with the ability to delete or add sequences to the TP domain without interfering with reverse transcription (19, 38–40).

The RT and RNaseH domain predictions in the catalytic core of P model were very similar to their structures in the individual domain models (Fig. S26B-C), and the catalytic core in the two- and four-domain genotype B models aligned with an RMSD of 2.37 Å (Fig. S26E). The fingers subdomain of the AlphaFold HBV P model was more similar to the structure found in the HIV RT than the fingers predicted by the Das et al. (25) RT domain model, with α-helixes and β-sheets of similar lengths and orientations as in the HIV RT (Figs. S26D and S27A). The C-terminal portion of the fingers subdomain, consisting of two β sheets, 2 α helices and a loop was oriented more vertically in the full-length model compared to their orientation in the isolated RT model and in the two domain RT-RH model. The orientation in the full-length model is more similar to what is seen with the HIV RT, implying that folding of the fingers into the HIV RT-like conformation may depend on the adjacent HBV spacer domain residues. A plausible nucleic acid binding channel between the two active sites featuring positively charged and neutral residues was present, although the channel was not as positively charged as in the HIV enzyme. Both catalytic domains in the full-length model accommodated Mg^++^ ions coordinated by the YMDD or D-E-D-D motifs in positions very similar to those in the HIV RT and RNaseH active sites (Fig. 4B and C). There were minor differences in the location of Mg^++^ ions in the RNaseH active sites between the genotype B and D models and between the catalytic core and full-length models that were due to differences in the side chain positions for the D-E-D-D residues, primarily E729 (Figs. S24E and S25C), but the ions were in plausible positions in all models.

### Nucleic acid docking

RNA:DNA heteroduplex structures (21 and 24 bp) were extracted from two HIV RT-RNaseH co-crystal structures (PDB: 4B3Q and 6BSH) and docked into the HBV RT-RNaseH two-domain catalytic core and full-length P models for genotypes B and D employing the Schrödinger BioLuminate module. Both full-length models and the genotype B catalytic core model docked the heteroduplex into the RT active site in a range of plausible poses, but none engaged both the RT and RNaseH active sites simultaneously. Rather, the heteroduplex was located just outside the RNaseH active site (Figs.5A, 5B, S28B, and S28D). Similar results were obtained with the genotype D two-domain RT-RNaseH model, but one pose aligned the DNA strand near the Mg^++^ ions in the RT active site and the RNA strand opposite the Mg^++^ ions in the RNaseH active site (Figs. S28E-G). The heteroduplex followed the nucleic acid binding channel predicted by the apo models in this genotype D model. The 19 bp spanning the RT and RNaseH active sites is similar to the 15-18 bp estimated for DHBV by measuring the length of the 5’ RNA fragment remaining when the RT active site reaches the end of the pgRNA (13). The infrequency of binding poses that engage both active sites is similar to what has been seen with HIV RT-RNaseH co-crystals where the heteroduplexes engaged the RT active site but were located outside the RNaseH active site (41–43). Many P residues contacted the heteroduplex among the various binding poses, with residues F59, R219, Q226, K378, H381, R387, S396, R397, R401, W404, K406, F407, V409, F434, K585, Y598, K614, W630, K631, R635, F642, Y658, Q662, K756, and Y779 contacting the heteroduplex in at least two binding poses. This variability between predicted binding poses is similar to the variation in residues contacting duplex nucleic acids between the three HIV RT-RNaseH heteroduplex co-crystals that has been attributed to the structures trapping the enzyme at different stages of its catalytic cycle (41, 42, 44).

**Figure 5.**
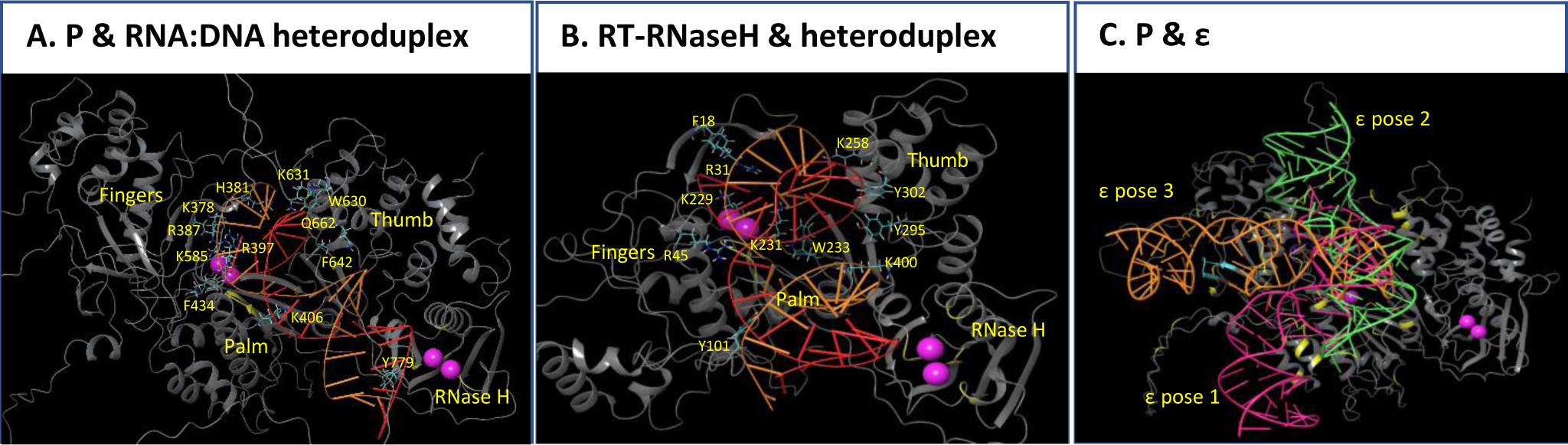
Computational nucleic acid docking into the HBV genotype B catalytic core and full-length P models. Docking was conducted using PIPER program in the BioLuminate module within the Schrödinger molecular analysis suite. **A. – C.** Representative binding poses between the indicated model and nucleic acids. *A.* and *B.* The fingers, palm, and thumb subdomains of the RT domain are labeled. Residues in the RT and RH domains contacting the RNA:DNA heteroduplex are shown as sticks and labeled. *C.* Representative poses are shown for the three classes of binding poses identified for the ε RNA. Locations of residues that interact with ε among the various binding poses are indicated: Violet, RT1 motif; Cyan, T3; Yellow, all other interacting residues.

The HBV ε RNA stem-loop that templates DNA priming [PDB: 6VAR (45)] was also docked into the full-length model of genotype B P (Figs. 5C and S29A-C). Diverse poses in which ε conformed closely to the surface of P were found that fell into three classes. Two of the classes wrapped end-to-end around the side of P, and the third class docked perpendicularly to the other poses. All poses contacted the palm subdomain, the TP domain, and the spacer domain, but none placed ε in the RT active site. Sixty-five percent of residues in direct contact with ε in the various poses were within residues 42-196 or 291-520 that are necessary for P to bind ε (46). The observation of diverse binding poses outside the RT active site suggests that one or more of them may represent a non-specific early step in binding between ε and P (Fig. 1B) (11).

Overall, the P models docked heteroduplexes in poses similar to what is seen with the HIV RT-RNaseH enzyme, and docked ε in manners consistent with what is known about how HBV P initially contacts ε. However, interpretation of the RNA docking studies is constrained by the diversity of binding poses for ε. This may stem from imprecisions in the accuracy of the AlphaFold models, limitations to the docking algorithm in recapitulating complex multi-domain nucleic acid interactions, lack of the chaperones that promote nucleic acid binding to P during the docking studies, and/or the ability of each model to reflect only one of P’s multiple conformations.

### Evaluation of the predicted relative orientation of the HBV RT and RNaseH domains

The relative orientation of the RT and RNaseH domains that form the catalytic core of P is highly plausible for four reasons. First, the HBV genotype B catalytic core aligns with the HIV enzyme with an RMSD of 3.75 Å (Fig. S24F) and with the Ty3 RT-RNaseH enzyme with an RMSD of 3.60 Å (Fig. S24H), although in both cases the RMSD value is exaggerated by positioning differences of the two catalytic domains that stem from the absence of a linker region in HBV P. Second, the two HBV active sites were connected with a continuous nucleic acid binding channel as is seen in other RT-RNaseH enzymes. Finally, the HBV two-domain catalytic core and the full length P models can dock RNA:DNA heteroduplexes with the DNA and RNA strands in their expected conformations.

### Assessment of the overall fold for the predicted HBV P model

Plausibility of the full-length genotype B P model was evaluated by (i) assessing the location of the T3 and RT1 motifs that form a bi-partite RNA binding motif (11), (ii) Assessing locations in the DHBV P model for residues whose surface exposures have been experimentally determined, and (iii) assessing how well the model provided a mechanistic explanation for mutations to residues that induce strong phenotypes in RNA binding, DNA priming, or RNaseH activity when mutated.

The T3 motif in the TP domain and the RT1 motif in the RT domain were previously found to form two parts of a discontinuous RNA binding motif in HBV and DHBV that is obscured when P is initially translated but that promotes binding of ε to P upon being exposed by cellular chaperones (Fig. 1B) (11, 20, 21, 47, 48). Despite being separated by 207 residues, the T3 and RT1 motifs formed a nearly contiguous motif wrapping over the fingers subdomain and into the interior face of the RT domain (Figs. 6A). All the binding poses predicted by docking ε to P had at least one contact between ε and RT1, but few contacts were detected with T3 (Fig. 5C). A strand of the TP domain overlaid the RT1 motif near where it contacts T3, and one of the few structured portions of the spacer domain obscures most of T3. Comparing the full P model to conformations defined during identification of the T3 and RT1 motifs implies the AlphaFold model of P may approximate the *closed complex* (11) initially formed upon translation of P that cannot bind specifically to ε (Fig. 1B).

**Figure 6.**
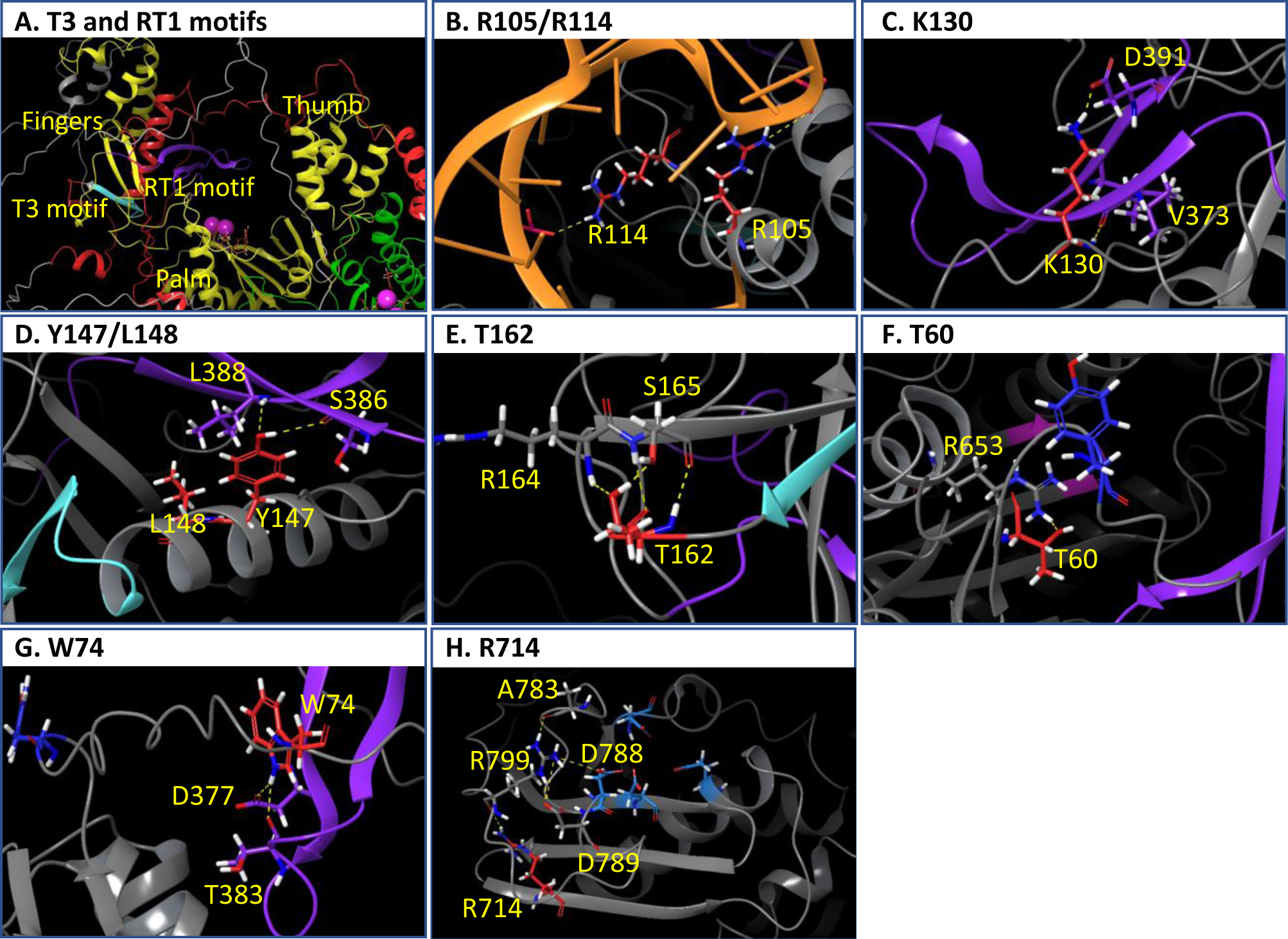
Motifs and residues used for assessing validity of the predicted for genotype B model for full-length P. Interactions among HBV motifs and key residues are shown in the genotype B model. **A.** Positions of the T3 and RT1 motifs. Magenta sphere, Mg^++^ ions. **C. - H.** Networks of polar interactions involving the wild-type residues at the indicated positions are indicated as yellow dashed lines. Mutating the indicated residues yielded strong phenotypes as described in the text. Comparisons between the wild-type and mutant interaction networks are in Fig. S30. Red, sites that were mutated and used for validation analyses; Orange, ε RNA with phosphates interacting with P shown; Violet, RT1 motif; Cyan, T3 motif; Blue, Y63 priming residue; Magenta, YMDD RT active site motif; Light blue, D-E-D-D motif.

Further support for the hepadnaviral P fold comes from the DHBV P model (Fig. S16; see below). First, partial proteolysis revealed that E17 or E18, E369, E370, E371, D468, E555, E565, E571, and E572 are protease accessible and D468 is inaccessible on the protein (21, 23). The location of these 11 residues on the DHBV P model is consistent with these observations. Second, E164, E176, and E199 are resistant to proteolysis in the absence of a chaperone-mediated conformational change but become sensitive upon chaperone action (21). E164 is partially obscured in a groove and could be exposed by minor structural changes, E176 is exposed in the model but could be covered by the spacer depending on its native conformation, and E199 is buried but could be exposed by shifting the helix in which it is located. E106 and E124 were protease resistant (21) but were exposed in the model, indicating either a weakness in the model or that the sites are obscured by the spacer or chaperones in the native holoenzyme complex. Finally, the epitope for monoclonal antibody 11 (residues 53-59) is constitutively exposed, and the epitopes for monoclonal antibodies 5 (residues 141-147) and 6 (residues 191-197) are obscured before the chaperone-mediated conformational change leading to formation of the *open complex* (Fig. 1B) (11, 21). The antibody 11 epitope is fully exposed in the DHBV P model, the antibody 6 epitope is in a deep groove that could be exposed by shifting the C-terminus of the spacer, and the spacer domain may overlay the antibody 5 epitope and be exposed by chaperone activation.

We next evaluated the ability of the P model to provide mechanistic explanations for effects of mutations that induce strong effects on HBV RNA binding, DNA priming, or RNaseH activity. We introduced these mutations into the full-length genotype B model and polar interactions involving the residues were identified in both the wild-type and mutant sequences. The wild-type and mutated models were then energy minimized using the Schrödinger OPSL4 force field and the effects of the mutations were assessed.

Four sets of mutations in the TP domain that impair RNA binding and/or RNA packaging into viral capsids can be explained by the proposed model for HBV P. R105A is deficient in RNA packaging, R114E reduces RNA binding and packaging, and both mutations ablate protein priming and DNA synthesis [R105 (48, 49); R114 (50)]. The model places these residues adjacent to each other on the outside of the fingers subdomain (Fig. 6B), and docking studies predict that they may bind to the phosphate backbone of ε (Fig. S30D-E). Mutating these residues removes these interactions, which would impair ε binding and the subsequent DNA priming. Second, K130 in the TP domain is predicted to contact D391 and the backbone of V373 in the RT1 motif in the RT domain (Fig. 6C), and it binds to ε in docking pose 2 (Fig. 5C). K130L ablates these interactions (Fig. S30F) and is defective in packaging RNA into viral capsids (51). This would be expected if the orientation, flexibility, and/or exposure of RT1 were changed by disrupting K130:D391 binding, or alternatively, if K130 works together with RT1 to promote ε binding as predicted by the docking results. Similarly, Y147 and L148 in the TP domain are adjacent to the T3 motif and L148 binds to the backbone of S386 and L388 in the RT1 motif (Fig. 6D). Y147A/L148A removes these contacts (Fig. S30G) and is deficient in RNA packaging into capsids for both HBV and DHBV (11, 47), as would be expected if disrupting these interactions between the TP and RT1 impaired T3 and/or RT1 function. Alternatively, residues 147-148 may comprise part of the T3 motif that was not identified when T3 was defined. Finally, T162 is two residues C-terminal to the T3 motif in the TP domain and interacts with R164 and S165 in the model. It is predicted to be adjacent to a turn between two strands of a β-sheet and to form a hydrogen bond with S165 and the backbone of R164, the last residue in the turn (Fig. 6E). T162P is defective in RNA packaging into capsids (52), which is dependent upon binding to ε that requires T3 function. Inserting a proline disrupts two polar interactions to R164 and S165 (Fig. S30H), which may disrupt the local structure or impede conformational changes required for the specific binding of ε to P.

Two interactions predicted by the HBV P model can explain mutations defective in DNA priming. T60 is predicted to be in the priming loop in the TP domain and to interact with S64 in the TP domain and R653 in the RT domain (Fig. 6F). T60E is defective in protein priming, but T60A has no phenotype (50). A nonpolar substitution at T60 would lessen its affinity with R653 in the RT domain and retain flexibility of the priming loop (Fig. S30A). In contrast, T60E inserts a negative charge which would increase affinity for R653 (Fig. S30B) and would likely impair folding of the priming loop downward, reducing Y63’s ability to access the RT active site during priming. Second, W74 at the C-terminus of the priming loop hydrogen bonds with T383 and D377 within the RT1 motif to help place Y63 above the YMDD RT active site motif (Fig. 6G). W74A removes these interactions (Fig. S30C) and is deficient in protein priming but not RNA packaging, as would be expected if the mutation disrupted a network of interactions anchoring the priming loop containing Y63 in a position where it can shift downwards to access the RT active site and ε during priming (48, 53).

Finally, the effects of the RNaseH-deficient R714A mutation (31) were assessed. R714 anchors R799 in position by hydrogen bonding to the backbone of R799, which is in a loop near the C-terminus of the RNaseH domain. R799 forms polar interactions with A783, D788, and D789 (Fig. 6H). R714A removes an interaction at the base of this interaction network (Fig. S30I) which would increase flexibility of the loop. Shifting the loop would inhibit RNaseH activity by moving R799 from its location adjacent to the D-E-D-D motif and altering the location of D788 in the D-E-D-D motif.

### Inhibitor docking

#### Reverse transcriptase active site

Plausibility of the RT active site fold was evaluated by docking the active triphosphate form of three nucleos(t)ide analog HBV reverse transcriptase inhibitors, Entecavir, Tenofovir, and Lamivudine (5) into the HBV RT active site in the catalytic core and full-length P models for genotypes B and D. A double-stranded DNA duplex from the HIV:substrate co-crystal (PDB: 1RTD) was superimposed on the HBV RT active site in the four models, and then the inhibitors were computationally docked using Glide XP program within the Schrödinger Maestro suite. The three inhibitors bound in the expected poses in all four models, with the α and β phosphates being coordinated by the active site Mg^++^ ions, and the inhibitor pairing with the template strand of the primer-template (Fig. 7A). Docking energies ranged from -8.86 to -13.3 kCal/mol (Table 1).

**Figure 7.**
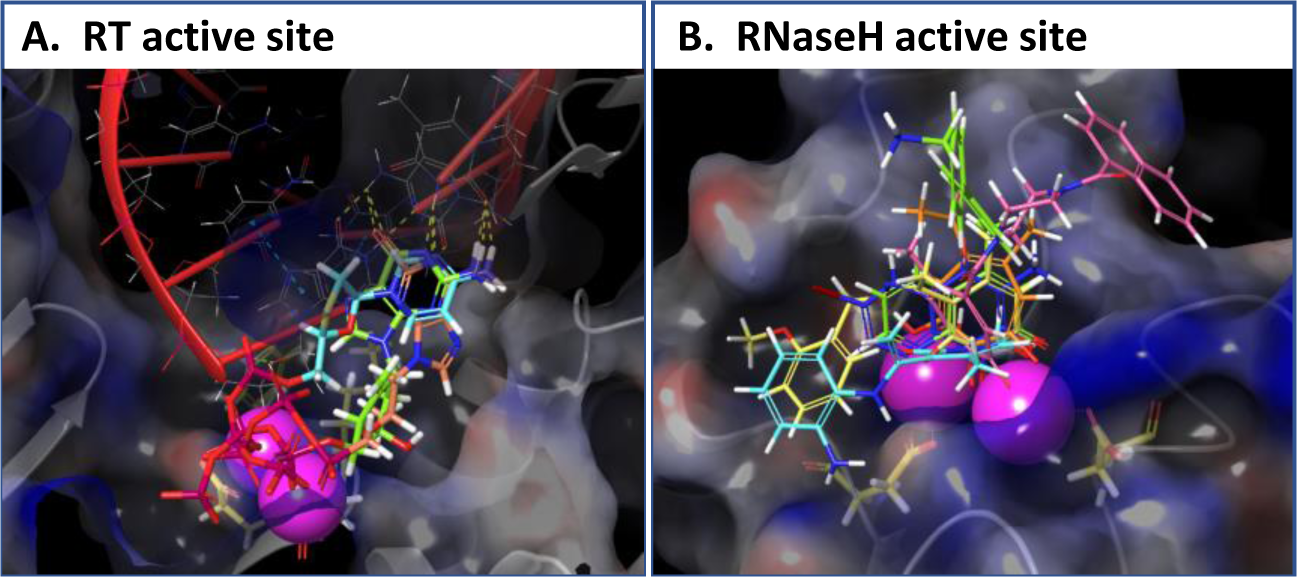
Computational docking of compounds into the RT and RNaseH active sites. Compounds were docked using the Glide XP program within the Schrödinger molecular analysis suite. Magenta spheres, Mg^++^ ions. **A.** Triphosphate form of nucleos(t)ide analog drugs docked into the RT active site. Red; DNA duplex; Orange, Tenofovir triphosphate; Green, Entecavir triphosphate; Cyan, Lamivudine triphosphate. **B.** RNaseH inhibitors docked into the RNaseH active site. Yellow, Compound **A25**; Cyan, **208**; Pink, **404**; Orange, **110**; Blue, **1073**; Green, **12**.

**Table 1.** Docking scores for the most stable poses of compounds into the RT and RNaseH active sites

| Compound<sup>1</sup> | Class<sup>2</sup> | gtB<br>RT-RNaseH | gtB P | gtD<br>RT-RNaseH | gtD P |
| --- | --- | --- | --- | --- | --- |
| **RT active site<sup>3</sup>** |  |  |  |  |  |
| Entecavir-TP | NA | -13.31 | -13.01 | -12.92 | -10.42 |
| Tenofovir-TP | NA | -11.05 | -8.86 | -9.27 | -12.96 |
| Lamivudine-TP | NA | -11.76 | -8.94 | -9.61 | -11.31 |
| **RNaseH active site<sup>3</sup>** |  |  |  |  |  |
| 208 | HPD | -10.82 | -9.45 | -9.58 | -9.38 |
| A25 |  | -10.69 | -9.71 | -9.92 | -10.74 |
| 110 | αHT | -7.88 | -7.78 | -7.64 | -7.80 |
| 404 |  | -8.64 | -8.72 | -8.31 | -8.47 |
| 12 | HNO | -7.44 | -8.12 | -5.10 | -7.46 |
| 1073 |  | -7.72 | -7.73 | -7.39 | -7.64 |

<sup>1</sup> Compounds described in (15, 27, 54-57)
<sup>2</sup> NA, nucleos(t)ide analog; HPD, *N*-hydroxypyridinedione; $\alpha$ HT, $\alpha$ -hydroxytropolone; HNO, *N*-hydroxynaphthyridinone
<sup>3</sup> Values in kCal/mol

#### Ribonuclease H active site

HBV RNaseH active site inhibitors were docked into the RNaseH active site in the genotype B and D catalytic core and full-length models using Glide XP. Inhibitors were α-hydroxytropolones (compounds **110** and **404**), *N*-hydroxypyridinediones (**208** and **A25**), and *N*-hydroxynapthyridinones (**12** and **1073**) (15, 27, 54–57). All four models docked HBV RNaseH inhibitors in poses where the compounds coordinated the Mg^++^ ions via their trident of metal chelating atoms (Fig. 7B), as predicted from the experimentally determined binding poses of similar compounds against the HIV RNaseH (58–62) and the need for an intact metal chelating trident of atoms for the inhibitors to work against HBV. The compounds adopted a variety of binding poses in the active site as anticipated given their wide structural diversity, usually along the predicted nucleic acid binding groove, with docking energies of -5.1 to -10.8 kCal/mol (Table 1).

### Effects of HBV’s genetic diversity on the models

HBV has nine genotypes (2) with P proteins that range from 832-845 aa long and differ by 11-16% at the amino acid level (Table S1). To evaluate how these variations may affect P’s structure, models for P proteins from all genotypes were generated using AlphaFold (Figs. S7-S15) and compared to the genotype B model. All models had similar pLDDT score profiles, with the C-terminal half of the TP domain and the catalytic core having high pLDDT values (∼70-90), while there were regions of much lower pLDDT values for the spacer domain and the N- and C-termini. Superpositioning all full-length models against the genotype B model revealed the same overall predicted fold, with an average pairwise RMSD of 1.98 Å (1.67 to 2.16 Å) outside the highly variable spacer domain (Figs. S25A-H). The key features of the genotype B model were conserved in all models, including the cupping of the TP domain around the catalytic core, the position of the priming tyrosine residue above the YMDD active site motif, the relative orientation of the T3 and RT1 motifs, and the relative orientation of the RT and RNaseH domains (Figs. S7-S15). Differences were present among the models, such as in the position of E729 from the D-E-D-D motif in the genotype F model compared to the others, that likely stem from limitations to the AlphaFold algorithm and/or and true structure variations.

### Models of animal P proteins and non-retroviral reverse transcriptases

To evaluate the phylogenetic conservation of the HBV P fold, AlphaFold models were constructed for P proteins from animal hepadnaviruses (2, 63), including woodchuck hepatitis virus (WHV, rodent), DHBV (bird), skink HBV (SkHBV, reptile), Tibetan frog HBV (TFHBV, amphibian), and tetra metahepadnavirus (TMDV, fish) (Fig. 8, Table S1, and Figs. S16-S20). These viruses share the same basic four-domain structure of P and replicate by protein-primed reverse transcription (63). We also generated a model for P from the rockfish nackednavirus (RNDV; Fig. S21).

**Figure 8.**
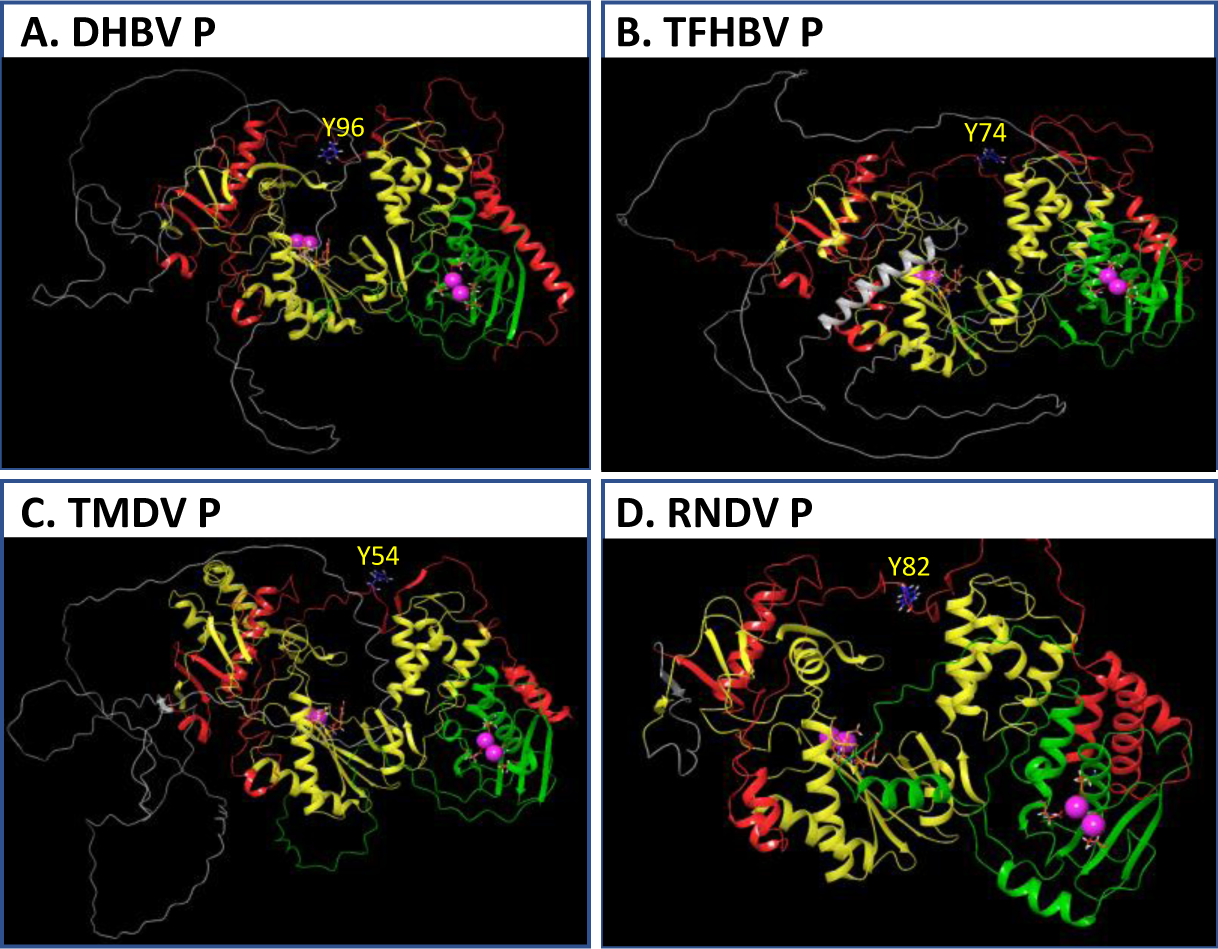
Predicted folds of animal hepadnavirus and nackednavirus P proteins. **A.** DHBV (avian hepadnavirus). **B.** TFHBV (frog hepadnavirus). **C.** TMDV (fish hepadnavirus). **D.** RNDV (fish nackednavirus). Red, TP domain; Gray, spacer domain; Yellow, RT domain; Green, RNaseH domain. The priming tyrosine residues, YMDD RT active site motif residues, and D-E-D-D RNaseH active site residues are shown as sticks. Putative priming residues are labeled. Magenta spheres, Mg^++^ ions.

Nackednaviruses and hepadnaviruses diverged about 400 million years ago, before the lineage that became the hepadnaviruses acquired its surface glycoproteins (63). Consequently, nackednavirus P proteins lack the spacer domain that encodes part of the surface protein gene in an overlapping reading frame. The priming mechanism is conserved well enough between HBV, DHBV, and RNDV that their P proteins can prime DNA synthesis using ε RNAs from the other viruses with only minor nucleotide substitutions in ε (64).

All animal virus P proteins shared the same basic predicted fold, with the catalytic core of the RT and RNaseH domain forming a globular unit featuring a contiguous nucleic acid binding groove, the TP domain cupping around the catalytic core, and the priming Y residue poised over the YMDD active site motif (Figs. 8, S16-S21). All models had the key A-E DNA polymerase active site motifs and the D-E-D-D RNaseH motifs in positions and orientations analogous to their locations in the HBV P model. The T3 and RT1 motifs in all six animal P protein models were in the same relative position as in HBV P. As expected, the spacer domain in the animal hepadnaviruses was unstructured, and RNDV had only a very short sequence linking the TP to the RT domain (Fig. 8D). The genotype B HBV P model superimposed well with the RNDV RT model (RMSD = 2.76 Å; Fig. 27D), with the largest differences outside of the spacer domain being in the flexible priming loop, the tip of the fingers subdomain, and position of the RNaseH domain (Fig. S27D).

These predicted structures imply that the hepadnaviral P fold existed before the split between the nackednaviruses and hepadnaviruses 400 million years ago (63). The models also imply that the evolutionary adoption of an envelope in the hepadnaviruses included insertion of sequences for part of the surface glycoproteins into sequences encoding amino acids corresponding to 174 – 192 of the modern nackednavirus RNDV.

Finally, we compared the predicted hepadnaviral P fold to models for reverse transcriptases from cauliflower mosaic virus (CaMV) (65, 66) and the mitochondrial retrotransposon pFOXC3 from the fungus *Fusarium oxysporum* f. sp. Matthiolae (67, 68) (Figs. S22 and S23). CaMV primes reverse transcription using a host tRNA similar to the retroviruses, whereas the pFOXC3 RT primes reverse transcription using an unidentified tyrosine residue in a mechanism similar to the hepadnaviruses. The predicted folds for the CaMV and pFOXC3 RTs revealed a nucleic acid binding groove and also YVDD (CaMV) and YADD (pFOXC3) motifs analogous to the hepadnaviral YMDD RT active site motif in their expected positions. The pFOXC3 model aligned to the HBV P RT-RNaseH domains in the full-length model with an RMSD = 4.52 Å (Fig. S27E) and positioned Y35 in a plausible position for priming reverse transcription. The HBV RT and RNaseH domains aligned with the CaMV RT-RNaseH domains with an RMSD = 3.96 Å (Fig. S27G). Neither the CaMV nor the pFOXC3 models had domains analogous to the hepadnaviral TP or spacer domains. The pFOXC3 model did not have a readily identifiable RNaseH domain or an identifiable D-E-D-D RNaseH active site motif. The CaMV RNaseH active site was predicted to contain a D-E-E-D motif rather than D-E-D-D, and the CaMV N-terminal aspartic proteinase was predicted to form a globular domain absent in the hepadnaviral proteins. Overall, these models imply that the hepadnaviral fold is a feature of the nackednavirus/hepadnavirus lineages rather than being widespread among non-retroviral eukaryotic RTs.

### Limitations to the models

There are two primary limitations to this analysis. First, all models reported here are predictions. The models are well supported by the available biological and biochemical data, but they are not based on experimental structural data. Second, HBV P is a structurally dynamic protein, so there is no single structure that can represent all of its biologically relevant conformations. The full-length P models presented here likely correspond best to the *closed conformation* previously identified for the enzyme (11), but they could represent an average of multiple conformations adopted by P. The catalytic core of the genotype D two-domain model may approximate the catalytically active form of the enzyme in the *active conformation* (11) of P because it can dock an RNA:DNA heteroduplex into the RT and RNaseH active sites simultaneously.

### Utility of the models

HBV P has resisted empirical structural analyses for decades, so the models reported here provide the first global structural approximation of the enzyme. They reveal a novel orientation of the TP domain relative to the rest of the enzyme that makes clear predictions for ε binding, DNA priming, and the conformational dynamics of P during viral genomic replication. The catalytic core models enable formulation of detailed hypotheses regarding the mechanisms of DNA elongation during reverse transcription and how the RT and RNaseH domains coordinate during reverse transcription. Finally, the models provide a least moderate assistance for structure-guided drug design against the RT and RNaseH active sites, and they open the door for rational drug discovery against targets other than the two enzymatic active sites.

## Methods

### Sequences and protein structures employed

All P protein sequences employed are in Supplemental Dataset 1. Table S1 contains the Genbank numbers or literature references for the sequences plus their pairwise identities with the genotype B reference sequence (AB554017). Supplemental Text 1 lists the protein domain and motif boundaries employed.

The Tibetan frog hepatitis B virus (TFHBV) sequence lacks an in-frame ATG for P, so the start of P was arbitrarily set to match the N-terminal length of the HBV genotype B sequence.

### Molecular modeling with AlphaFold

Sequences were folded using AlphaFold2 Advanced (https://colab.research.google.com/github/sokrypton/ColabFold/blob/main/beta/AlphaFold2_advanced.ipynb) using default parameters. Five models were generated for each sequence, and the model having the highest pLDDT value was refined by the Amber-Relax module to optimize side chain orientation. The relaxed structure was used for all analyses. The predicted protein database (PDB) files are in Supplemental Dataset 2.

### Superposition studies

Superposition studies were done using Protein Structure Alignment in Maestro module of Schrödinger suite (Schrödinger LLC, New York, NY, USA). Superpositions were done with no constraints, with the exception that the HBV P and catalytic core superpositions with HIV RT-RNase H, Ty3 RT and CaMV RT were done by selecting residues in fingers, palm, and thumb subdomains of HBV RT domain due to differences in the sizes of the linker domain between HBV and the other enzymes.

### Nucleic acid docking

A 21 bp heteroduplex (PDB: 4B3Q) and a 24 bp heteroduplex (PDB: 6BSH) were extracted from HIV RT-RNase H co-crystals and docked into HBV P models using the PIPER program in the BioLuminate module of the Schrödinger suite (Schrödinger LLC, New York, NY, USA). Similarly, an NMR structure for ε (PDB: 6VAR) was docked into HBV P. Protein and nucleic acid structures were prepared with Protein Preparation wizard in BioLuminate module. Proteins were protonated using Epik at pH 7.5 +/- 2, hydrogen bonds were assigned employing PROPKA at pH 7.5 and finally energy minimization was done with the OPSL4 force field. Each protein model was docked with both substrates separately without any constraints, and 30 docking poses were generated in each case. Final poses were selected based on substrate placement into the binding channel between active sites of RT and RNase H domains while considering the engagement of RNA or DNA strands with active site residues.

### Compound docking

Compound docking into the HBV P models employed the Glide XP program in the Schrödinger Maestro module. Triphosphates for the nucleos(t)ide analogs Entecavir, Lamivudine and Tenofovir were extracted from HIV-1 RT-RNase H:inhibitor co-crystals (PDB: 5XN1, 6KDJ & 3JSM) for docking. RNase H inhibitors were selected for structural diversity among the α-hydroxytropolones, *N*-hydroxypyridinediones and *N*-hydroxynapthyridinones. All ligands were prepared with LigPrep (Schrödinger LLC), where compound energy minimizations were done with OPSL4 force field, different deprotonation states of the ligands were generated using Epik at pH 7.5 +/-2 to set the ionization state of the metal binding motifs, and compounds were desalted and tautomerized while retaining their chirality. The HBV RT-RNase H and full-length P protein structures containing Mg^++^ ions in the appropriate ionization states were prepared with the Schrödinger Protein Preparation wizard as described above. A 20 bp dsDNA substrate was placed into the RT active site of the HBV RT-RNase H and P models for nucleos(t)ide analog docking by superposition of HIV RT-RNase H:dsDNA co-crystal (PDB: 1RTD). The RT active site docking grid was defined by placing Entecavir-triphosphate from the HIV RT-RNaseH structure co-crystal (PDB: 5XN1) into the active site of RT domain of HBV models by superposing the YMDD motifs. The RNase H docking grid was defined by placing β-thujaplicinol into the active site by superposing the D-E-D-D motif from the HIV RNaseH:β-thujaplicinol co-crystal (PDB: 3K2P) onto the HBV model. The centroids of the ligands in the active sites were used to create 10 Å receptor grids for docking. Docking employed Schrödinger Glide XP with default settings.

### Identification of mutants for model validation

Mutations to HBV P with clear phenotypic effects were identified from the literature as indicated, with the TP domain analyses especially benefitting from Clark et al. (50). Relevant sites were mapped onto the full-length HBV genotype B P model (genotype B numbers are used, which may differ from the source numbers if a different genotype was used). Potential hydrogen bonds were identified using PyMOL’s *Find polar contacts* function (h-bond cutoff center = 4.0 Å); all predicted polar interactions assessed were ≤3.0 Å, implying moderate to strong interactions. Interactions identified by these analyses were interpreted in context of their published phenotypes to propose mechanisms by which the mutations may have altered P function.

## Supplemental Data

**Supplemental Text 1.** Domain and motif boundaries used in these analyses.

**Supplemental Figures 1-23.** Images of all predicted models generated here and key quality metrics for the models.

**Supplemental Figures 24-27.** Images of the superpositions used in these analyses.

**Supplemental Figures 28-29.** Detailed images for the docking experiments.

**Supplemental Figures 30.** Comparisons of interaction networks for the wild-type and mutant residues used to validate the HBV P model.

**Supplemental Table 1.** Pairwise identities of P proteins relative to P from HBV genotype B.

**Supplemental Dataset 1.** Sequences employed in modeling.

**Supplemental Dataset 2.** Protein database (PDB) files for all predicted models.

## Supporting information

Tajwar et al supplementary data

Supplemental dataset S1- sequence employed

emental dataset S2- PDB files

## Acknowledgements

The authors have no conflicts to declare. This work was supported by NIH grants R01 AI150610 and R01 AI148362 and Department of Defense grant W81XWH-18-1-0307 to JET. We thank Dr. John Kennell for input on the pFOX-C3 reverse transcriptase, Dr. Juan Villa for early modeling work on the HBV RNaseH, and the AlphaFold team for enabling this study.

## References

1. Seeger C, Zoulim F, & Mason WS (2021) Hepadnaviridae. Fields Virology, Fields Virology, (Wolters Kluwer, Philadelphia), Vol 2: DNA Viruses, pp 640–682.

2. Glebe D, Goldmann N, Lauber C, & Seitz S (2021) HBV evolution and genetic variability: Impact on prevention, treatment and development of antivirals. Antiviral Res 186:104973.

3. Polaris Observatory Collaborators (2018) Global prevalence, treatment, and prevention of hepatitis B virus infection in 2016: a modelling study. Lancet Gastroenterol Hepatol 3(6):383–403.

4. Trepo C, Chan HL, & Lok A (2014) Hepatitis B virus infection. Lancet 384(9959):2053–2063.

5. Pierra Rouviere C, Dousson CB, & Tavis JE (2020) HBV replication inhibitors. Antiviral Res 179:104815.

6. Ghany MG (2017) Current treatment guidelines of chronic hepatitis B: The role of nucleos(t)ide analogues and peginterferon. Best practice & research. Clinical gastroenterology 31(3):299–309.

7. Tavis JE & Badtke MP (2009) Hepadnaviral Genomic Replication. *Viral Genome Replication*, eds Cameron CE, Goette M, & Raney KD (Springer Science+Business Media, LLC, New York), pp 129–143.

8. Beck J & Nassal M (2007) Hepatitis B virus replication. World J Gastroenterol 13(1):48–64.

9. Hu J, Toft DO, & Seeger C (1997) Hepadnavirus assembly and reverse transcription require a multi-component chaperone complex which is incorporated into nucleocapsids. EMBO J. 16:59–68.

10. Hu J, Flores D, Toft D, Wang X, & Nguyen D (2004) Requirement of heat shock protein 90 for human hepatitis B virus reverse transcriptase function. J.Virol. 78(23):13122–13131.

11. Badtke MP, Khan I, Cao F, Hu J, & Tavis JE (2009) An interdomain RNA binding site on the hepadnaviral polymerase that is essential for reverse transcription. Virology 390(1):130–138.

12. Lanford RE, Notvall L, Lee H, & Beams B (1997) Transcomplementation of Nucleotide Priming and Reverse Transcription between Independently Expressed TP and RT Domains of the Hepatitis B Virus Reverse Transcriptase. J.Virol. 71:2996–3004.

13. Loeb DD, Hirsch RC, & Ganem D (1991) Sequence-independent RNA cleavages generate the primers for plus strand DNA synthesis in hepatitis B viruses: implications for other reverse transcribing elements. EMBO J. 10(11):3533–3540.

14. Poch O, Sauvaget I, Delarue M, & Tordo N (1989) Identification of four conserved motifs among the RNA-dependent polymerase encoding elements. EMBO J. 8:3867–3874.

15. Tavis JE, et al. (2013) The hepatitis B virus ribonuclease H is sensitive to inhibitors of the human immunodeficiency virus ribonuclease H and integrase enzymes. PLoS pathogens 9(1):e1003125.

16. Yang W & Steitz TA (1995) Recombining the structures of HIV integrase, RuvC and RNase H. Structure. 3(2):131–134.

17. Nowotny M (2009) Retroviral integrase superfamily: the structural perspective. EMBO Rep. 10(2):144–151.

18. Villa JA, et al. (2016) Purification and enzymatic characterization of the hepatitis B virus ribonuclease H, a new target for antiviral inhibitors. Antiviral Res 132:186–195.

19. Zhang Z & Tavis JE (2006) The Duck Hepatitis B Virus Reverse Transcriptase Functions as a Full-length Monomer. J.Biol.Chem. 281(47):35794–35801.

20. Cao F, et al. (2005) Identification of an essential molecular contact point on the duck hepatitis B virus reverse transcriptase. J.Virol. 79(16):10164–10170.

21. Stahl M, Beck J, & Nassal M (2007) Chaperones activate hepadnavirus reverse transcriptase by transiently exposing a C-proximal region in the terminal protein domain that contributes to epsilon RNA binding. J.Virol. 81(24):13354–13364.

22. Wang X & Hu J (2002) Distinct requirement for two stages of protein-primed initiation of reverse transcription in hepadnaviruses. J Virol 76(12):5857–5865.

23. Lin L, Wan F, & Hu J (2008) Functional and structural dynamics of hepadnavirus reverse transcriptase during protein-primed initiation of reverse transcription: effects of metal ions. J.Virol. 82(12):5703–5714.

24. Buhlig TS, et al. (2020) Molecular, Evolutionary, and Structural Analysis of the Terminal Protein Domain of Hepatitis B Virus Polymerase, a Potential Drug Target. Viruses 12(5).

25. Das K, et al. (2001) Molecular modeling and biochemical characterization reveal the mechanism of hepatitis B virus polymerase resistance to lamivudine (3TC) and emtricitabine (FTC). J. Virol. 75(10):4771–4779.

26. Xu X, et al. (2016) Modeling the functional state of the reverse transcriptase of hepatitis B virus and its application to probing drug-protein interaction. BMC Bioinformatics 17 Suppl 8:280.

27. Li Q, et al. (2020) Amide-containing alpha-hydroxytropolones as inhibitors of hepatitis B virus replication. Antiviral Res 177:104777.

28. Potenza N, et al. (2007) Optimized expression from a synthetic gene of an untagged RNase H domain of human hepatitis B virus polymerase which is enzymatically active. Protein Expr.Purif. 55(1):93–99.

29. Hayer J, et al. (2014) Ultradeep pyrosequencing and molecular modeling identify key structural features of hepatitis B virus RNase H, a putative target for antiviral intervention. J Virol 88(1):574–582.

30. Hyjek M, Figiel M, & Nowotny M (2019) RNases H: Structure and mechanism. DNA Repair (Amst) 84:102672.

31. Ko C, et al. (2014) Residues Arg703, Asp777, and Arg781 of the RNase H domain of hepatitis B virus polymerase are critical for viral DNA synthesis. J Virol 88(1):154–163.

32. Jumper J, et al. (2021) Highly accurate protein structure prediction with AlphaFold. Nature 596(7873):583–589.

33. Mariani V, Biasini M, Barbato A, & Schwede T (2013) lDDT: a local superposition-free score for comparing protein structures and models using distance difference tests. Bioinformatics 29(21):2722–2728.

34. Donlin MJ, Szeto B, Gohara DW, Aurora R, & Tavis JE (2012) Genome-wide networks of amino acid covariances are common among viruses. J Virol 86(6):3050–3063.

35. Lanford RE, Kim YH, Lee H, Notvall L, & Beames B (1999) Mapping of the hepatitis B virus reverse transciptase TP and RT domains by transcomplementation for nucleotide priming and by protein-protein interaction. J.Virol. 73:1885–1893.

36. Beck J & Nassal M (2001) Reconstitution of a functional duck hepatitis B virus replication initiation complex from separate reverse transcriptase domains expressed in Escherichia coli. J.Virol. 75(16):7410–7419.

37. Boregowda RK, Adams C, & Hu J (2012) TP-RT domain interactions of duck hepatitis B virus reverse transcriptase in cis and in trans during protein-primed initiation of DNA synthesis in vitro. J Virol 86(12):6522–6536.

38. Tavis JE & Ganem D (1993) Expression of functional hepatitis B virus polymerase in yeast reveals it to be the sole viral protein required for correct initiation of reverse transcription. Proc.Natl.Acad.Sci.U.S.A. 90:4107–4111.

39. Beck J & Nassal M (2003) Efficient Hsp90-independent in vitro activation by Hsc70 and Hsp40 of duck hepatitis B virus reverse transcriptase, an assumed Hsp90 client protein. J.Biol.Chem. 278(38):36128–36138.

40. Hu J, Toft D, Anselmo D, & Wang X (2002) In vitro reconstitution of functional hepadnavirus reverse transcriptase with cellular chaperone proteins. J.Virol. 76(1):269–279.

41. Huang H, Chopra R, Verdine GL, & Harrison SC (1998) Structure of a covalently trapped catalytic complex of HIV-1 reverse transcriptase: implications for drug resistance. Science 282(5394):1669–1675.

42. Ha B, et al. (2021) High-resolution view of HIV-1 reverse transcriptase initiation complexes and inhibition by NNRTI drugs. Nat Commun 12(1):2500.

43. Sarafianos SG, et al. (2001) Crystal structure of HIV-1 reverse transcriptase in complex with a polypurine tract RNA:DNA. EMBO J 20(6):1449–1461.

44. Das K, Martinez SE, DeStefano JJ, & Arnold E (2019) Structure of HIV-1 RT/dsRNA initiation complex prior to nucleotide incorporation. Proc Natl Acad Sci U S A 116(15):7308–7313.

45. LeBlanc RM, et al. (2021) Structural insights of the conserved “priming loop” of hepatitis B virus pre-genomic RNA. J Biomol Struct Dyn:1–13.

46. Hu J & Boyer M (2006) Hepatitis B virus reverse transcriptase and epsilon RNA sequences required for specific interaction in vitro. J.Virol. 80(5):2141–2150.

47. Cao F, et al. (2014) Sequences in the terminal protein and reverse transcriptase domains of the hepatitis B virus polymerase contribute to RNA binding and encapsidation. J Viral Hepat 21(12):882–893.

48. Jones SA, Clark DN, Cao F, Tavis JE, & Hu J (2014) Comparative analysis of hepatitis B virus polymerase sequences required for viral RNA binding, RNA packaging, and protein priming. J Virol 88(3):1564–1572.

49. Shin YC, Park S, & Ryu WS (2011) A conserved arginine residue in the terminal protein domain of hepatitis B virus polymerase is critical for RNA pre-genome encapsidation. The Journal of general virology 92(Pt 8):1809–1816.

50. Clark DN, Flanagan JM, & Hu J (2017) Mapping of Functional Subdomains in the Terminal Protein Domain of Hepatitis B Virus Polymerase. J Virol 91(3).

51. Roychoudhury S, Faruqui AF, & Shih C (1991) Pregenomic RNA encapsidation analysis of eleven missense and nonsense polymerase mutants of human hepatitis B virus. Journal of Virology 65:3617–3624.

52. Blum HE, Galun E, Liang TJ, von Weizsacker F, & Wands JR (1991) Naturally occurring missense mutation in the polymerase gene terminating hepatitis B virus replication. J Virol 65(4):1836–1842.

53. Shin YC, Ko C, & Ryu WS (2011) Hydrophobic residues of terminal protein domain of hepatitis B virus polymerase contribute to distinct steps in viral genome replication. FEBS Lett 585(24):3964–3968.

54. Edwards TC, et al. (2019) Inhibition of HBV replication by N-hydroxyisoquinolinedione and N-hydroxypyridinedione ribonuclease H inhibitors. Antiviral Res 164:70–80.

55. Edwards TC, et al. (2017) Inhibition of hepatitis B virus replication by N-hydroxyisoquinolinediones and related polyoxygenated heterocycles. Antiviral Res 143:205–217.

56. Lu G, et al. (2015) Hydroxylated Tropolones Inhibit Hepatitis B Virus Replication by Blocking the Viral Ribonuclease H Activity. Antimicrob Agents Chemother 59(2):1070–1079.

57. Chauhan R, et al. (2021) Efficient Inhibition of Hepatitis B Virus (HBV) Replication and cccDNA Formation by HBV Ribonuclease H Inhibitors during Infection. Antimicrob Agents Chemother 65(12):e0146021.

58. Lansdon EB, et al. (2011) Structural and binding analysis of pyrimidinol carboxylic acid and N-hydroxy quinazolinedione HIV-1 RNase H inhibitors. Antimicrob Agents Chemother 55(6):2905–2915.

59. Kirschberg TA, et al. (2009) RNase H active site inhibitors of human immunodeficiency virus type 1 reverse transcriptase: design, biochemical activity, and structural information. J.Med.Chem. 52(19):5781–5784.

60. Himmel DM, et al. (2009) Structure of HIV-1 reverse transcriptase with the inhibitor beta-Thujaplicinol bound at the RNase H active site. Structure. 17(12):1625–1635.

61. Chung S, et al. (2011) Synthesis, activity, and structural analysis of novel alpha-hydroxytropolone inhibitors of human immunodeficiency virus reverse transcriptase-associated ribonuclease H. J.Med.Chem. 54(13):4462–4473.

62. Billamboz M, et al. (2011) 2-hydroxyisoquinoline-1,3(2H,4H)-diones as inhibitors of HIV-1 integrase and reverse transcriptase RNase H domain: influence of the alkylation of position 4. European journal of medicinal chemistry 46(2):535–546.

63. Lauber C, et al. (2017) Deciphering the Origin and Evolution of Hepatitis B Viruses by Means of a Family of Non-enveloped Fish Viruses. Cell Host Microbe 22(3):387–399 e386.

64. Beck J, Seitz S, Lauber C, & Nassal M (2021) Conservation of the HBV RNA element epsilon in nackednaviruses reveals ancient origin of protein-primed reverse transcription. Proc Natl Acad Sci U S A 118(13).

65. Haas M, Bureau M, Geldreich A, Yot P, & Keller M (2002) Cauliflower mosaic virus: still in the news. Mol Plant Pathol 3(6):419–429.

66. Volovitch M, Modjtahedi N, Yot P, & Brun G (1984) RNA-dependent DNA polymerase activity in cauliflower mosaic virus-infected plant leaves. EMBO J 3(2):309–314.

67. Walther TC & Kennell JC (1999) Linear mitochondrial plasmids of F. oxysporum are novel, telomere-like retroelements. Mol Cell 4(2):229–238.

68. Galligan JT, Marchetti SE, & Kennell JC (2011) Reverse transcription of the pFOXC mitochondrial retroplasmids of Fusarium oxysporum is protein primed. Mob DNA 2(1):1.

