## Supplementary material for "Predicted structure of the hepatitis B virus polymerase reveals an ancient conserved protein fold": Tajwar et al supplementary data: Tajwar et al Supplementary data.pdf

**Supplemental Text 1** Domain and motif boundaries employed.

Domain and motif residue boundaries are given as inclusive numbers.

| **HBV gtA** | |
|---|---|
| TP | 1 – 183 |
| SP | 184 – 358 |
| RT | 359 – 691 |
| RH | 692 – 845 |
| Priming residue Y65 | |
| YMDD motif | 549 – 552 |
| DEDD moitf | 702, 731, 750, 790 |
| T3 motif | 155 – 162 |
| RT1 motif | 370 – 400 |

| **HBV gtB** | |
|---|---|
| TP | 1 – 181 |
| SP | 182 – 356 |
| RT | 357 – 689 |
| RH | 690 – 843 |
| Priming residue Y63 | |
| YMDD motif | 549 – 552 |
| DEDD motif | 700, 729, 748, 788 |
| T3 motif | 153 – 160 |
| RT1 motif | 368 – 398 |

| **HBV gtB Poch box motifs** | |
|---|---|
| A | 421 – 437 |
| B | 509 – 535 |
| C | 546 – 552 |
| D | 576 – 587 |
| E | 593 – 603 |

| **HBV gtC** | |
|---|---|
| TP | 1 – 181 |
| SP | 182 – 356 |
| RT | 357 – 689 |
| RH | 690 – 843 |
| Priming residue Y63 | |
| YMDD motif | 549-552 |
| DEDD motif | 700, 729, 748, 788 |
| T3 motif | 153 – 160 |
| RT1 motif | 368 – 398 |

| **HBV gtD** | |
|---|---|
| TP | 1 – 180 |
| SP | 181 – 345 |
| RT | 346 – 678 |
| RH | 679 – 832 |
| Priming residue Y63 | |
| YMDD motif | 538 – 541 |
| DEDD motif | 689, 718, 737, 777 |
| T3 motif | 153 – 160 |
| RT1 motif | 357 – 387 |

**HBV gtE**

| | |
|---|---|
| TP | 1 – 181 |
| SP | 182 – 355 |
| RT | 356 – 688 |
| RH | 689 – 842 |
| Priming residue Y63 | |
| YMDD motif | 549-552 |
| DEDD motif | 700, 729, 748, 788 |
| T3 motif | 154 – 160 |
| RT1 motif | 368 – 398 |

**HBV gtF**

| | |
|---|---|
| TP | 1 – 181 |
| SP | 182 – 356 |
| RT | 357 – 689 |
| RH | 690 – 843 |
| Priming residue | Y63 |
| YMDD motif | 549-552 |
| DEDD motif | 700, 729, 748, 788 |
| T3 motif | 153 – 160 |
| RT1 motif | 367 – 398 |

**HBV gtG**

| | |
|---|---|
| TP | 1 – 181 |
| SP | 182 – 355 |
| RT | 356 – 688 |
| RH | 689 – 842 |
| Priming residue Y63 | |
| YMDD motif | 548-551 |
| DEDD motif | 699, 728, 747, 787 |
| T3 motif | 153 – 160 |
| RT1 motif | 367 – 397 |

**HBV gtH**

| | |
|---|---|
| TP | 1 – 181 |
| SP | 182 – 356 |
| RT | 357 – 689 |
| RH | 690 – 843 |
| Priming residue | Y63 |
| YMDD motif | 549 – 552 |
| DEDD motif | 700, 729, 748, 788 |
| T3 motif | 153 – 160 |
| RT1 motif | 368 – 398 |

**HBV gtI**

| | |
|---|---|
| TP | 1 – 181 |
| SP | 182 – 356 |
| RT | 357 – 689 |
| RH | 690 – 843 |
| Priming residue | Y63 |
| YMDD motif | 549 – 552 |
| DEDD motif | 700, 729, 748, 788 |
| T3 motif | 154 – 160 |
| RT1 motif | 368 – 398 |

**WHV**

| | |
|---|---|
| TP | 1 – 188 |
| SP | 189 – 397 |
| RT | 398 – 728 |
| RH | 729 – 884 |
| Priming residue | Y68 |
| YMDD motif | 588 – 591 |
| DEDD motif | 739, 768, 787, 827 |
| T3 motif | 152 – 165 |
| RT1 motif | 409 – 439 |

**DHBV**

| | |
|---|---|
| TP | 1 – 201 |
| SP | 202 – 372 |
| RT | 373 – 653 |
| RH | 654 – 786 |
| Priming residue Y96 | |
| YMDD motif | 511 – 514 |
| DEDD motif | 666, 696, 715, 755 |
| T3 motif | 176 – 183 |
| RT1 motif | 383 – 415 |

**SkHBV**

| | |
|---|---|
| TP | 1 – 207 |
| SP | 208 – 460 |
| RT | 461 – 745 |
| RH | 746 – 889 |
| Priming residue Y98 | |
| YMDD motif | 601 – 604 |
| DEDD motif | 756, 787, 806, 846 |
| T3 motif | 178 – 185 |
| RT1 motif | 475 – 505 |

**TFHBV**

| | |
|---|---|
| TP | 1 – 192 |
| SP | 193 – 393 |
| RT | 394 – 678 |
| RH | 679 – 814 |
| Priming residue Y74 | |
| YMDD motif | 534 – 537 |
| DEDD motif | 689, 719, 738, 778 |
| T3 motif | 154 – 161 |
| RT1 motif | 408 – 438 |

**TMDV**

| | |
|---|---|
| TP | 1 – 162 |
| SP | 163 – 348 |
| RT | 349 – 672 |
| RH | 673 – 823 |
| Priming residue Y54 | |
| YMDD motif | 532 – 535 |
| DEDD motif | 683, 713, 732, 772 |
| T3 motif | 134 – 141 |
| RT1 motif | 360 – 390 |

**RNDV**

| | |
|---|---|
| TP | 1 – 173 |
| SP | 174 – 192 |
| RT | 193 – 465 |
| RH | 466 - 634 |
| Priming residue Y82 | |
| YMDD motif | 333 – 336 |
| DEDD motif | 486, 519, 536, 577 |
| T3 motif | 165 – 172 |
| RT1 motif | 205 – 237 |

**Table S1. Pairwise identities of P proteins relative to P from HBV genotype B isolate AB554017.**

| Virus or retrotransposon | Genotype or strain | Genbank number or publication | Length (residues) | Identity with HBV genotype B (%) | Number of gap residues in alignment (% of alignment) |
|---|---|---|---|---|---|
| HBV | A | AP007263 | 845 | 88 | 2 (0%) |
| | B | AB554017 | 843 | 100 | 0 (0%) |
| | C | AB560661 | 843 | 89 | 0 (0%) |
| | D | AB554016 | 832 | 86 | 11 (1%) |
| | E | AB091255 | 842 | 85 | 1 (0%) |
| | F | AB064316 | 843 | 84 | 0 (0%) |
| | G | AB064310 | 842 | 85 | 1 (0%) |
| | H | AB298362 | 843 | 84 | 0 (0%) |
| | I | MH368022 | 843 | 88 | 0 (0%) |
| WHV | | NC_004107 | 884 | 50 | 61 (6%) |
| DHBV | 3 | DQ195079 | 786 | 29 | 191 (20%) |
| SkHBV | | Lauber (8) | 889 | 27 | 230 (23%) |
| TFHBV | | Lauber (8) | 814 | 27 | 0 (20%) |
| TMDV | | Lauber (8) | 823 | 34 | 98 (11%) |
| RNDV | | Lauber (8) | 634 | 22 | 337 (37%) |
| CaMV | | M90542.1 | 679 | 16 | 244 (27%) |
| pFOX C3 | | AAD38504.1 | 527 | 14 | 352 (40%) |

**Dataset S1.** Sequences for all proteins modeled. Sequences are provided individually in FASTA format with all files compiled into within a .zip file.

**Dataset S2.** PDB files for all models. Coordinates for all models are provided in PDB format with all files compiled into a .zip file.

### Fig. S1. HBV TP domain, Genotype B, Isolate AB554017

#### Predicted fold

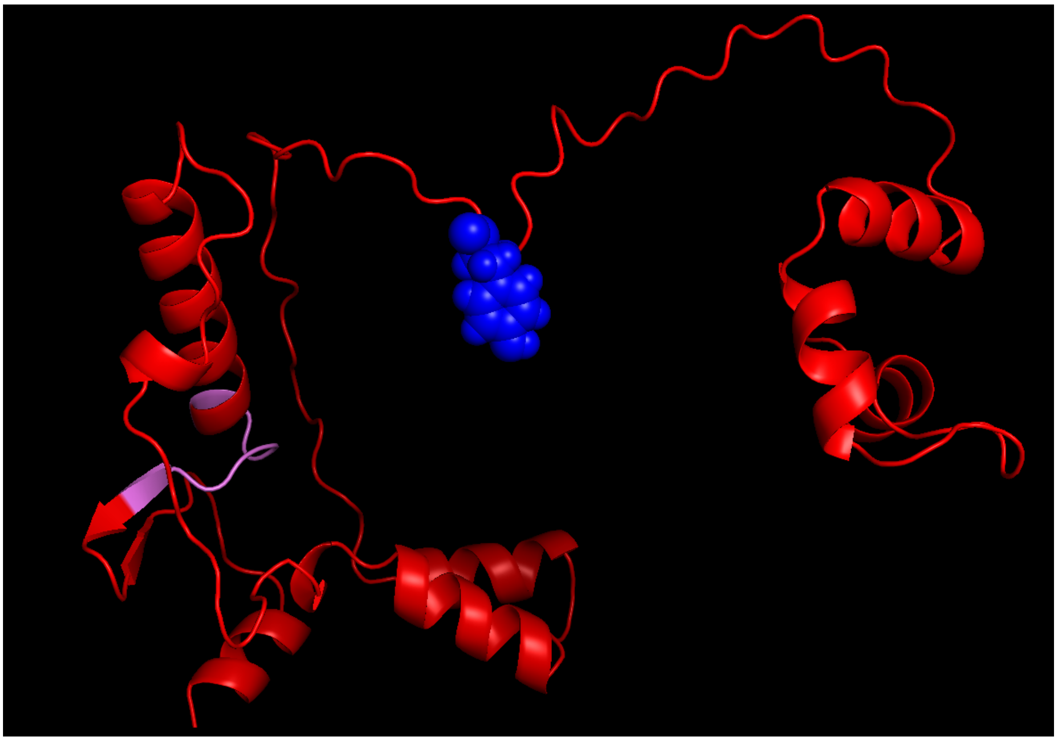

N-terminus, cyan; TP domain, red; Y63, blue; T3 motif, violet

#### MSA coverage

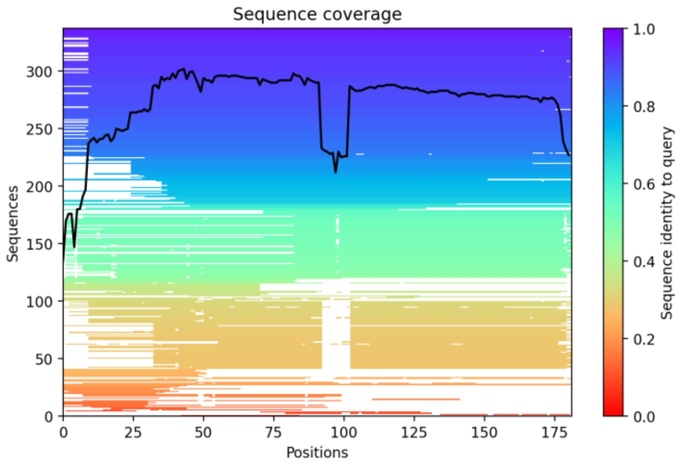


#### Predicted LDDT

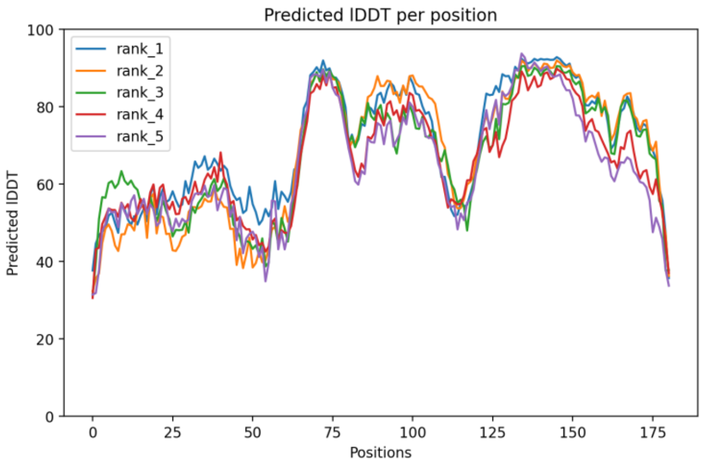


#### Predicted Distogram

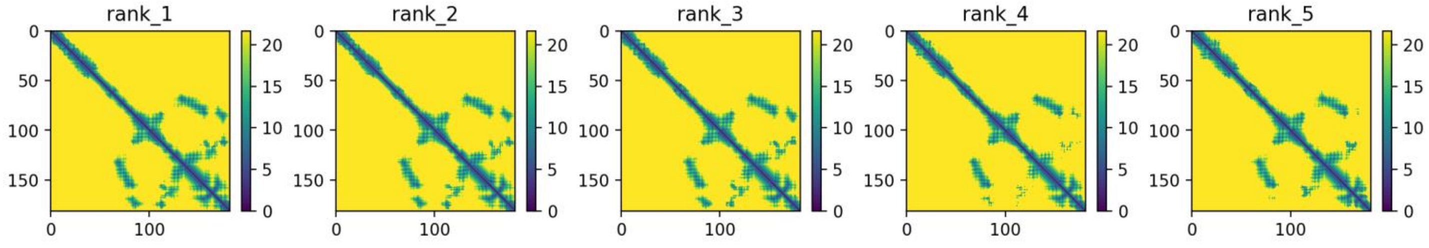


**Legend to Figs. S1-S23.** Images of all models and key quality metrics for the models. The predicted protein folds for all models are displayed in common orientations and color schemes. The color pallet used is: N-terminus, cyan; TP domain, red; T3 motif, violet; Spacer domain, grey; RT1 motif, light blue; RT domain, yellow; RNaseH domain, green; Priming residue, YMDD/YADD/YVDD motifs, D-E-D-D and D-E-E-D motifs, blue spheres; and Aspartate proteinase, magenta. Not all domains/motifs are found in all models. The predicted protein fold is shown the *Predicted Fold* panel. Depth of the multiples sequence alignment used to identify the coordinated variations from which the distance constraints are derived are in the *MSA Coverage* panel. The predicted LDDT value as a function of residue position is in the *Predicted LDDT* panel. Distograms based on the coordinated variations for each of the five models generated for each sequence are found in the *Predicted Distogram* panel.

### Fig. S2. HBV Spacer domain, Genotype B, Isolate AB554017

Predicted fold

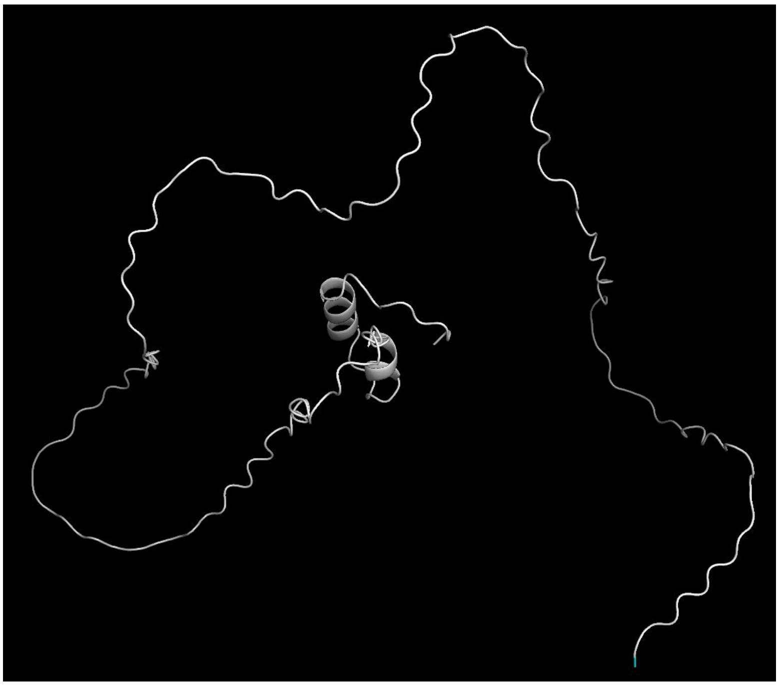

N-terminus, cyan; Spacer, grey

MSA coverage

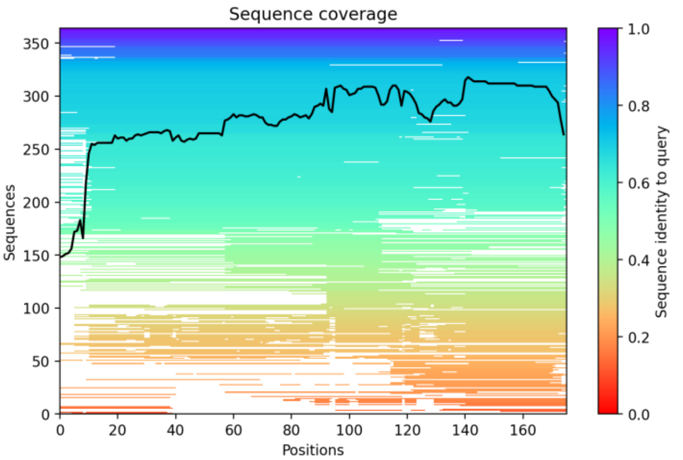


Predicted LDDT

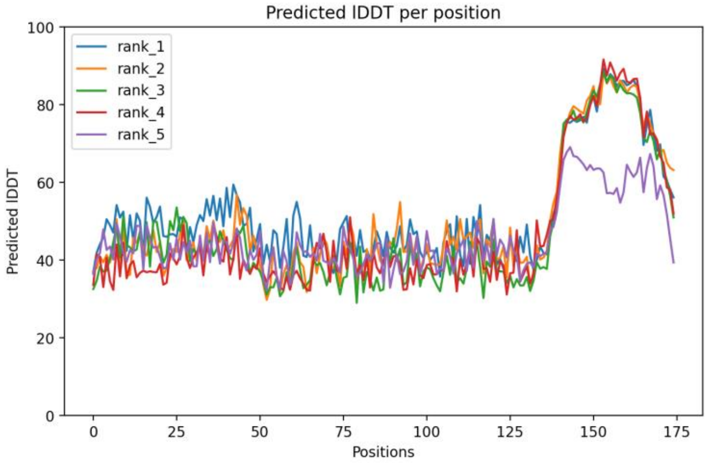


Predicted Distogram

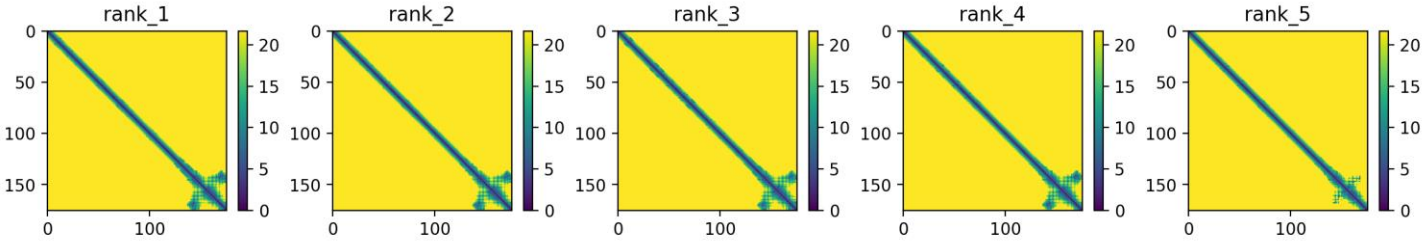


**Legend to Figs. S1-S23.** Images of all models and key quality metrics for the models. The predicted protein folds for all models are displayed in common orientations and color schemes. The color pallet used is: N-terminus, cyan; TP domain, red; T3 motif, violet; Spacer domain, grey; RT1 motif, light blue; RT domain, yellow; RNaseH domain, green; Priming residue, YMDD/YADD/YVDD motifs, D-E-D-D and D-E-E-D motifs, blue spheres; and Aspartate proteinase, magenta. Not all domains/motifs are found in all models. The predicted protein fold is shown the *Predicted Fold* panel. Depth of the multiples sequence alignment used to identify the coordinated variations from which the distance constraints are derived are in the *MSA Coverage* panel. The predicted LDDT value as a function of residue position is in the *Predicted LDDT* panel. Distograms based on the coordinated variations for each of the five models generated for each sequence are found in the *Predicted Distogram* panel.

### Fig. S3. HBV RT domain, Genotype B, Isolate AB554017

#### Predicted fold

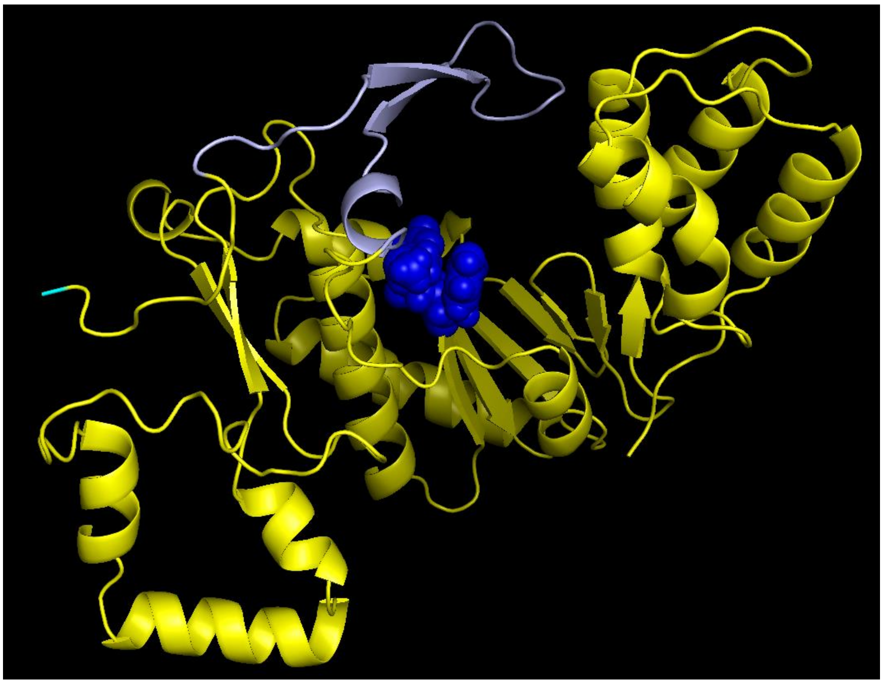

N-terminus, cyan; RT domain, yellow; YMDD, blue; RT1 motif, light blue

#### MSA coverage

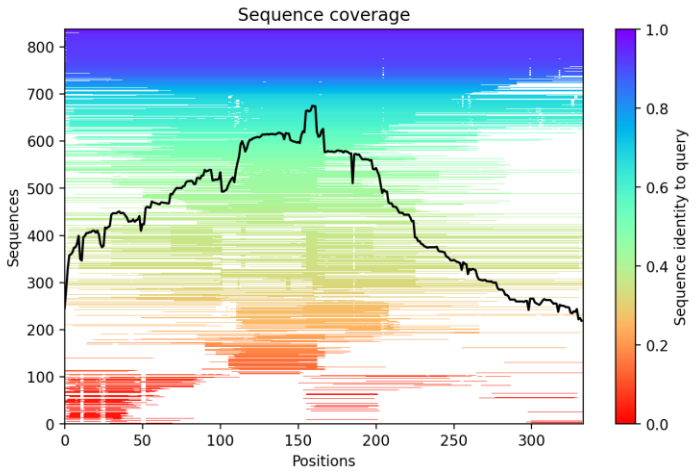


#### Predicted LDDT

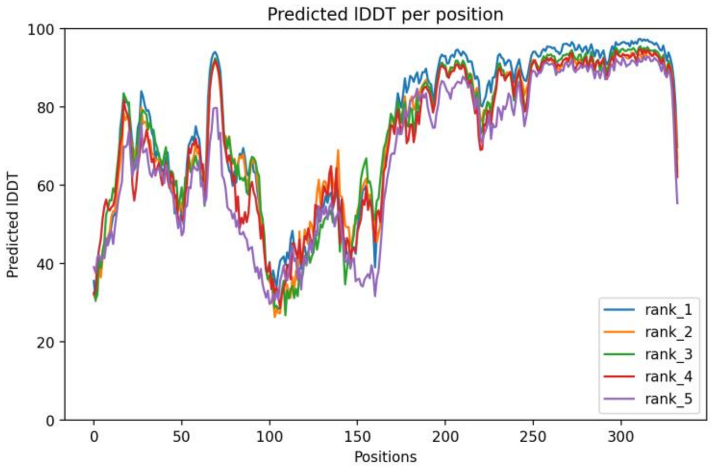


#### Predicted Distogram

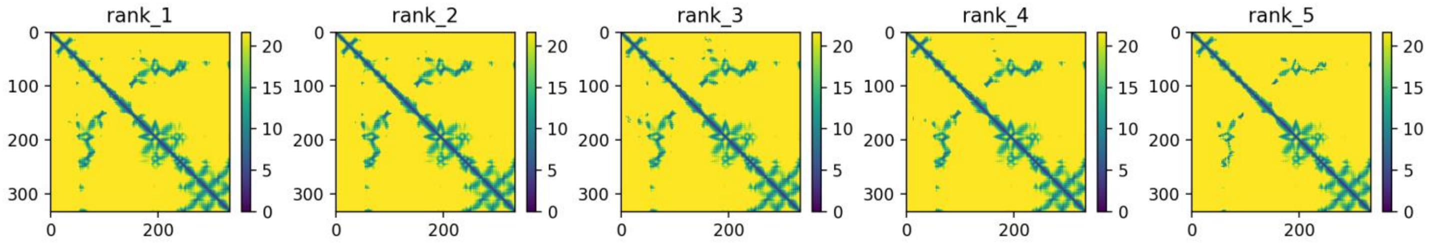


**Legend to Figs. S1-S23.** Images of all models and key quality metrics for the models. The predicted protein folds for all models are displayed in common orientations and color schemes. The color pallet used is: N-terminus, cyan; TP domain, red; T3 motif, violet; Spacer domain, grey; RT1 motif, light blue; RT domain, yellow; RNaseH domain, green; Priming residue, YMDD/YADD/YVDD motifs, D-E-D-D and D-E-E-D motifs, blue spheres; and Aspartate proteinase, magenta. Not all domains/motifs are found in all models. The predicted protein fold is shown the *Predicted Fold* panel. Depth of the multiples sequence alignment used to identify the coordinated variations from which the distance constraints are derived are in the *MSA Coverage* panel. The predicted LDDT value as a function of residue position is in the *Predicted LDDT* panel. Distograms based on the coordinated variations for each of the five models generated for each sequence are found in the *Predicted Distogram* panel.

### Fig. S4. HBV RNaseH domain, Genotype B, Isolate AB554017

#### Predicted fold

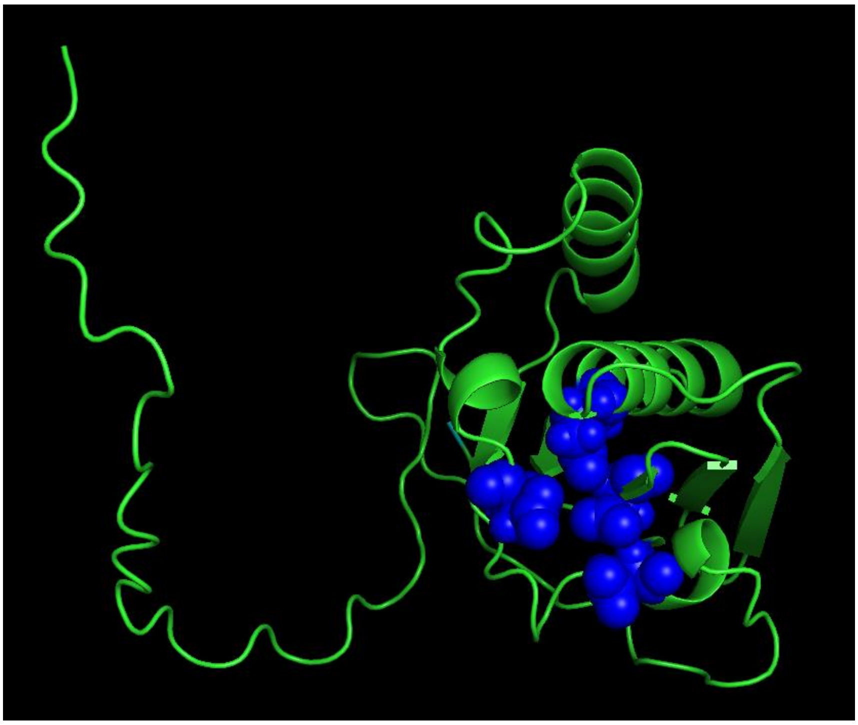

Cyan, N-terminus; RNaseH domain, green; D-E-D-D, blue

#### MSA coverage

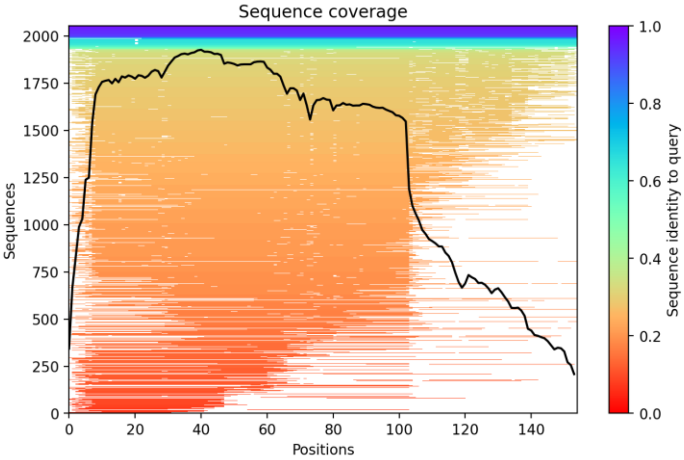


#### Predicted LDDT

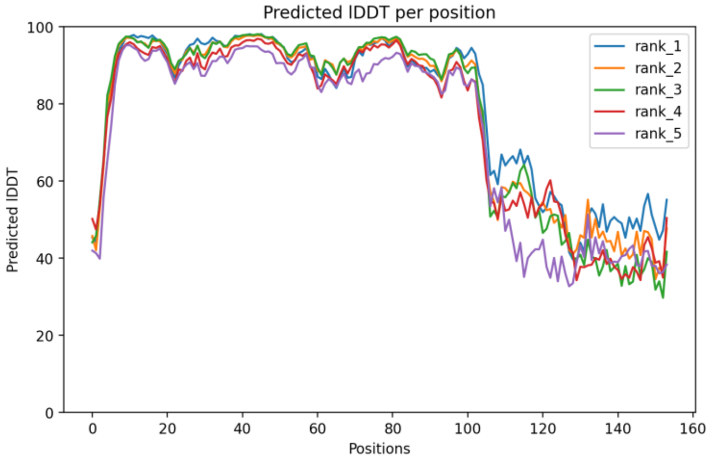


#### Predicted Distogram

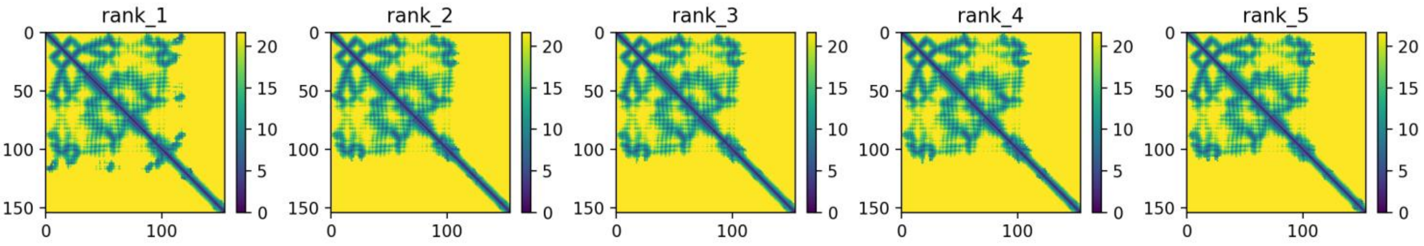


**Legend to Figs. S1-S23.** Images of all models and key quality metrics for the models. The predicted protein folds for all models are displayed in common orientations and color schemes. The color pallet used is: N-terminus, cyan; TP domain, red; T3 motif, violet; Spacer domain, grey; RT1 motif, light blue; RT domain, yellow; RNaseH domain, green; Priming residue, YMDD/YADD/YVDD motifs, D-E-D-D and D-E-E-D motifs, blue spheres; and Aspartate proteinase, magenta. Not all domains/motifs are found in all models. The predicted protein fold is shown the *Predicted Fold* panel. Depth of the multiples sequence alignment used to identify the coordinated variations from which the distance constraints are derived are in the *MSA Coverage* panel. The predicted LDDT value as a function of residue position is in the *Predicted LDDT* panel. Distograms based on the coordinated variations for each of the five models generated for each sequence are found in the *Predicted Distogram* panel.

### Fig. S5. HBV RT-RNaseH domains, Genotype B, Isolate AB554017

#### Predicted fold

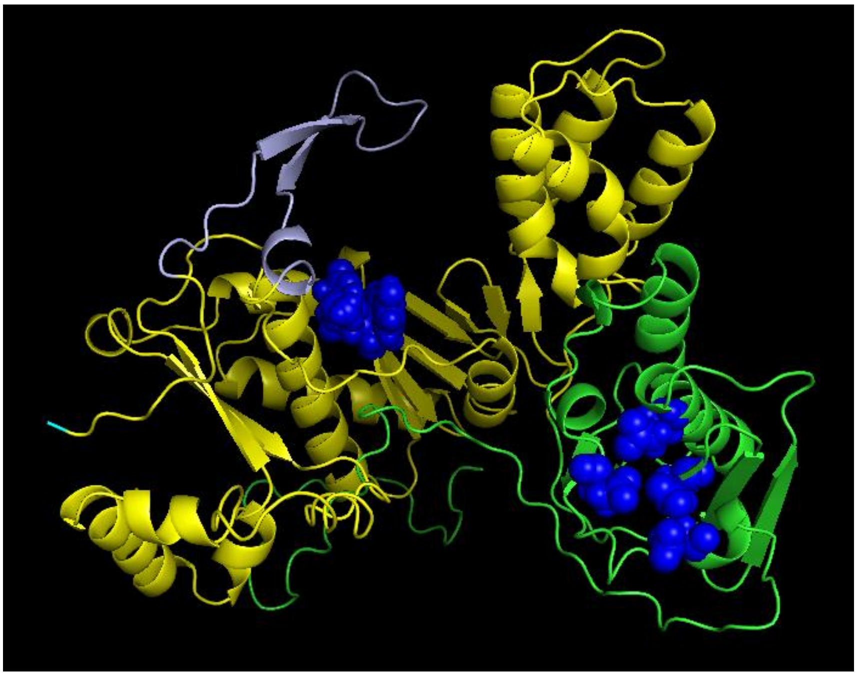

N-terminus, cyan; RT, yellow; RNaseH, green; YMDD and D-E-D-D, blue

#### MSA coverage

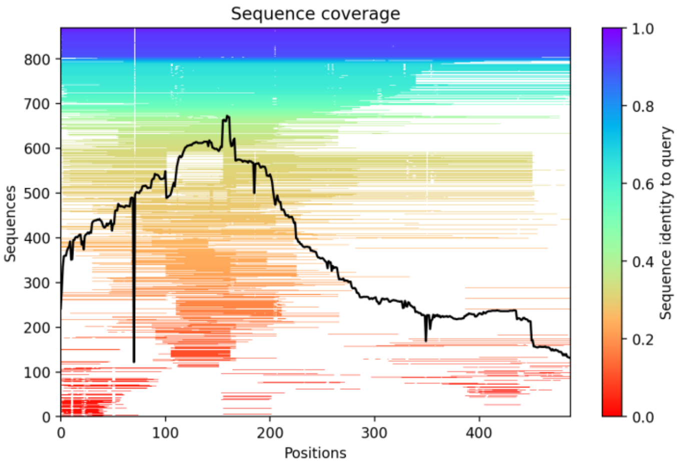


#### Predicted LDDT

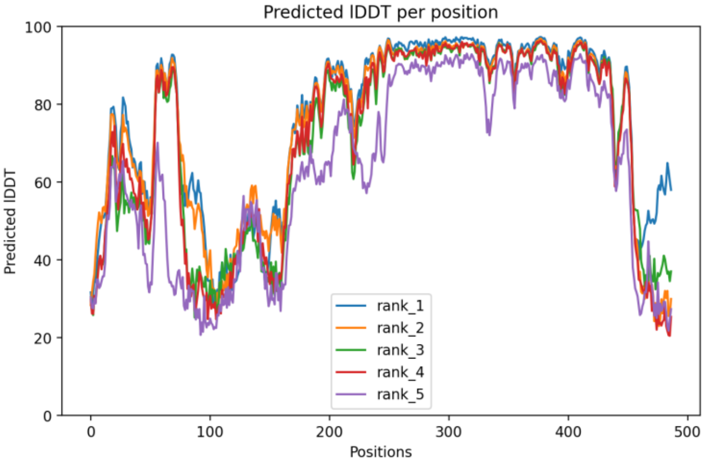


#### Predicted Distogram

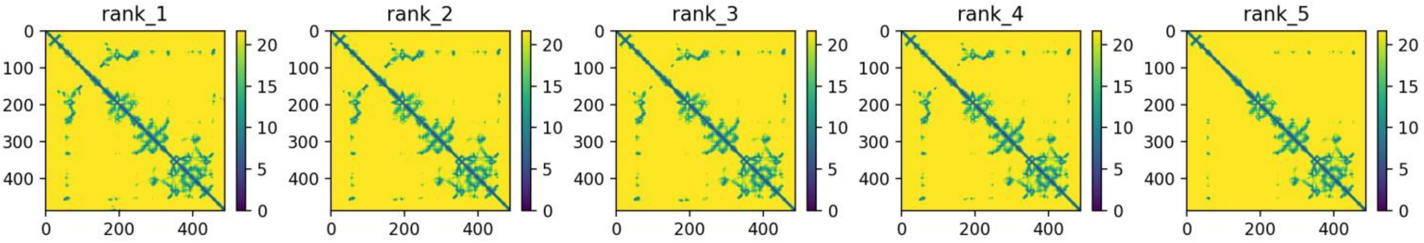


**Legend to Figs. S1-S23.** Images of all models and key quality metrics for the models. The predicted protein folds for all models are displayed in common orientations and color schemes. The color pallet used is: N-terminus, cyan; TP domain, red; T3 motif, violet; Spacer domain, grey; RT1 motif, light blue; RT domain, yellow; RNaseH domain, green; Priming residue, YMDD/YADD/YVDD motifs, D-E-D-D and D-E-E-D motifs, blue spheres; and Aspartate proteinase, magenta. Not all domains/motifs are found in all models. The predicted protein fold is shown the *Predicted Fold* panel. Depth of the multiples sequence alignment used to identify the coordinated variations from which the distance constraints are derived are in the *MSA Coverage* panel. The predicted LDDT value as a function of residue position is in the *Predicted LDDT* panel. Distograms based on the coordinated variations for each of the five models generated for each sequence are found in the *Predicted Distogram* panel.

### Fig. S6. HBV RT-RNaseH domains, Genotype D, Isolate AB554016

#### Predicted fold

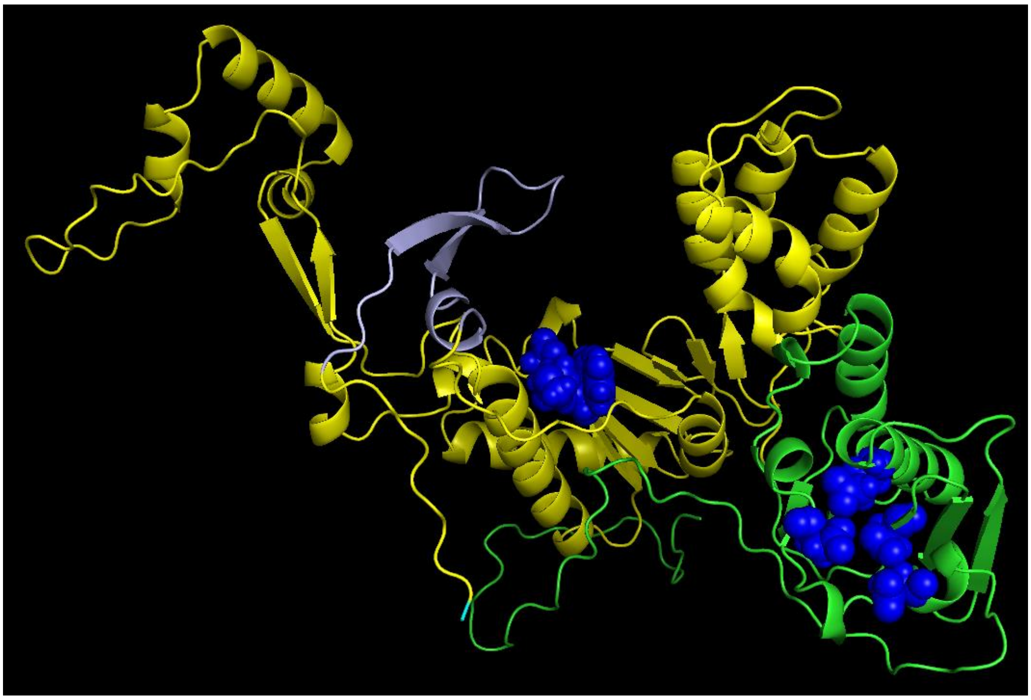

N-terminus, cyan; RT, yellow; RNaseH, green; YMDD and D-E-D-D, blue

#### MSA coverage

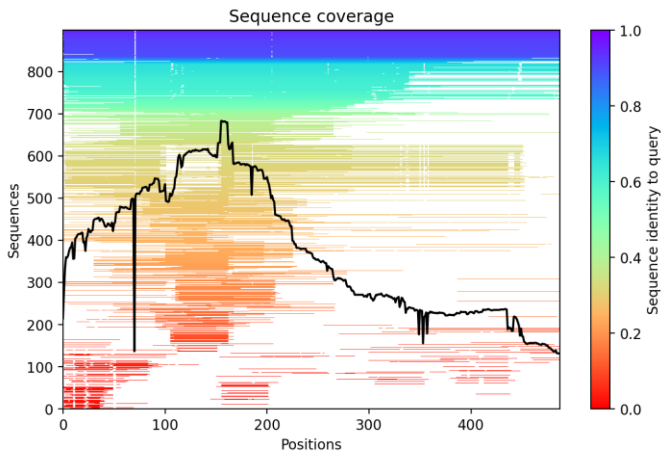


#### Predicted LDDT

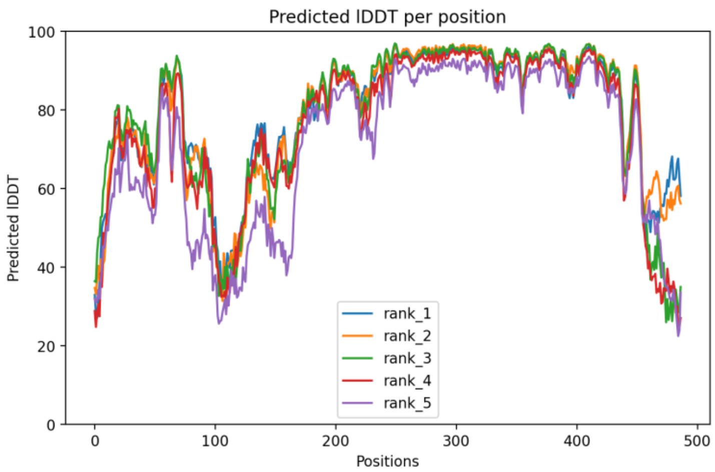


#### Predicted Distogram

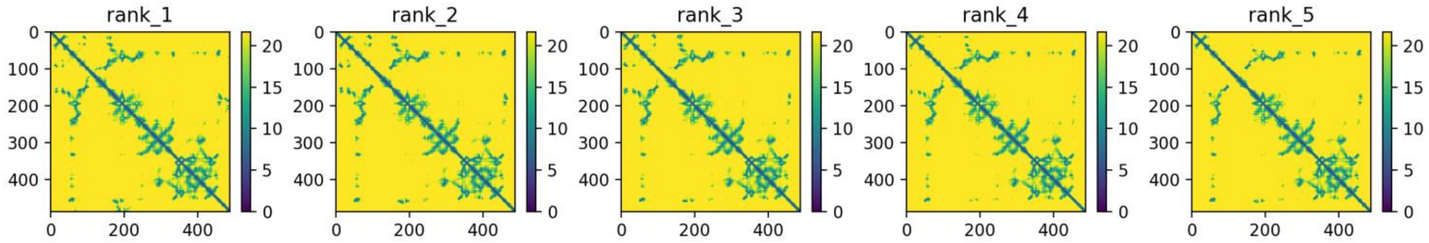


**Legend to Figs. S1-S23.** Images of all models and key quality metrics for the models. The predicted protein folds for all models are displayed in common orientations and color schemes. The color pallet used is: N-terminus, cyan; TP domain, red; T3 motif, violet; Spacer domain, grey; RT1 motif, light blue; RT domain, yellow; RNaseH domain, green; Priming residue, YMDD/YADD/YVDD motifs, D-E-D-D and D-E-E-D motifs, blue spheres; and Aspartate proteinase, magenta. Not all domains/motifs are found in all models. The predicted protein fold is shown the *Predicted Fold* panel. Depth of the multiples sequence alignment used to identify the coordinated variations from which the distance constraints are derived are in the *MSA Coverage* panel. The predicted LDDT value as a function of residue position is in the *Predicted LDDT* panel. Distograms based on the coordinated variations for each of the five models generated for each sequence are found in the *Predicted Distogram* panel.

### Fig. S7. HBV P, Genotype A, Isolate AP007263

#### Predicted fold

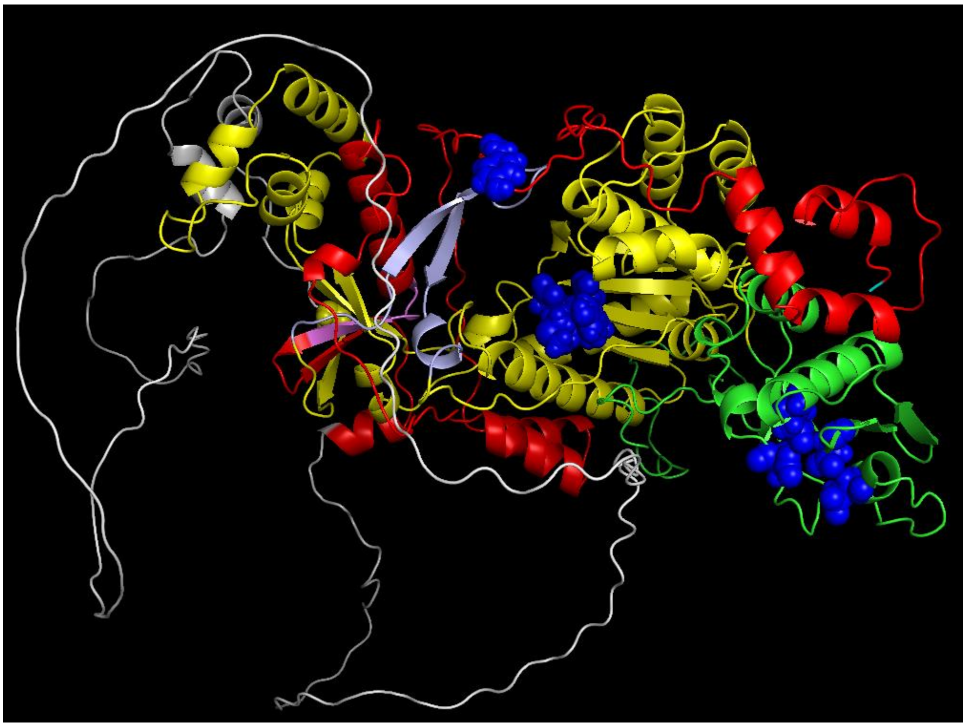

N-terminus, cyan; TP, red; T3 motif, violet; Spacer, grey; RT1 motif, light blue; RT, yellow; RNaseH, green; Y65, YMDD and D-E-D-D, blue

#### MSA coverage

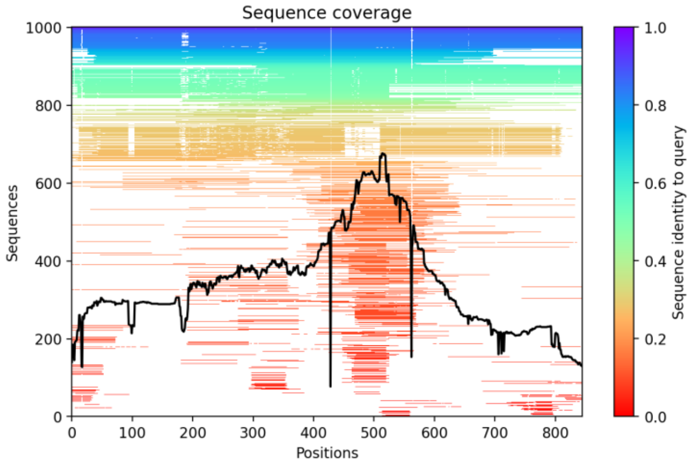


#### Predicted LDDT

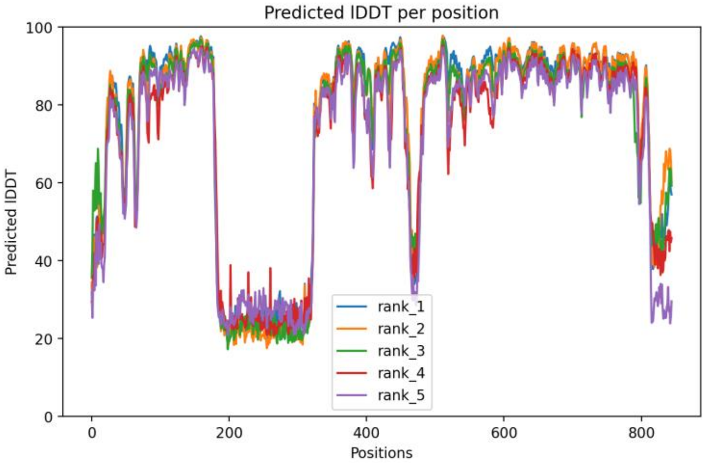


#### Predicted Distogram

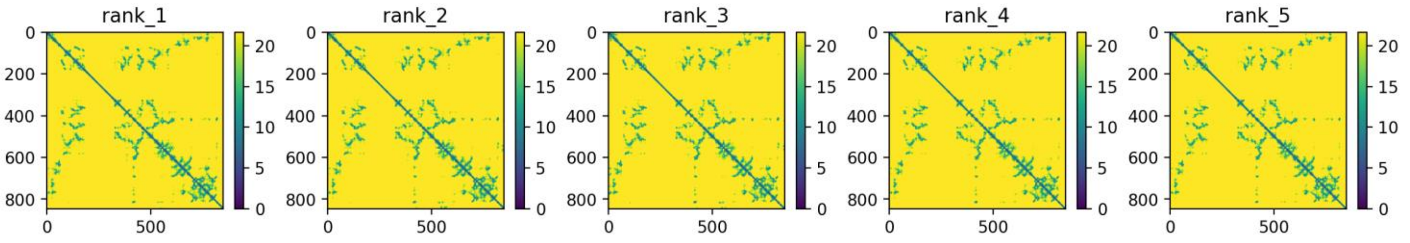


**Legend to Figs. S1-S23.** Images of all models and key quality metrics for the models. The predicted protein folds for all models are displayed in common orientations and color schemes. The color pallet used is: N-terminus, cyan; TP domain, red; T3 motif, violet; Spacer domain, grey; RT1 motif, light blue; RT domain, yellow; RNaseH domain, green; Priming residue, YMDD/YADD/YVDD motifs, D-E-D-D and D-E-E-D motifs, blue spheres; and Aspartate proteinase, magenta. Not all domains/motifs are found in all models. The predicted protein fold is shown the *Predicted Fold* panel. Depth of the multiples sequence alignment used to identify the coordinated variations from which the distance constraints are derived are in the *MSA Coverage* panel. The predicted LDDT value as a function of residue position is in the *Predicted LDDT* panel. Distograms based on the coordinated variations for each of the five models generated for each sequence are found in the *Predicted Distogram* panel.

### Fig. S8. HBV P, Genotype B, Isolate AB554017

#### Predicted fold

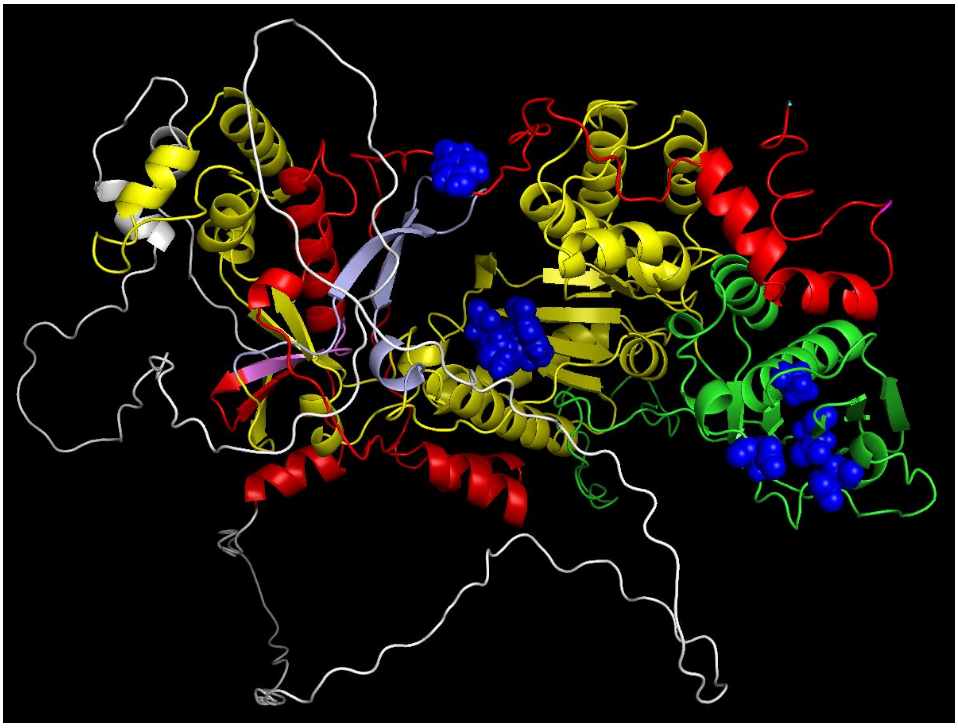

N-terminus, cyan; TP, red; T3 motif, violet; Spacer, grey; RT1 motif, light blue; RT, yellow; RNaseH, green; Y63, YMDD and D-E-D-D, blue

#### MSA coverage

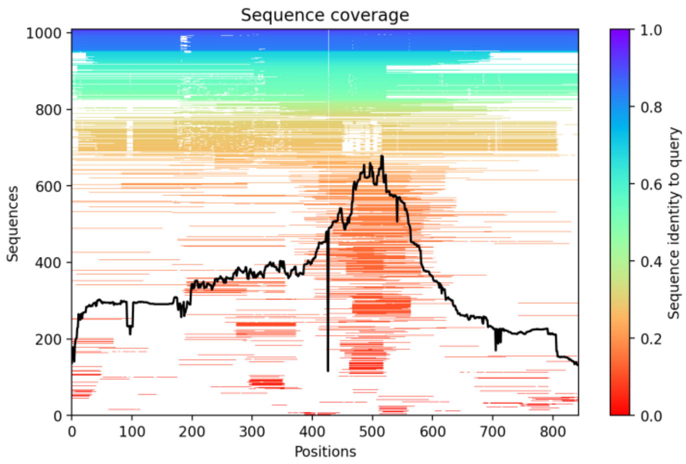


#### Predicted LDDT

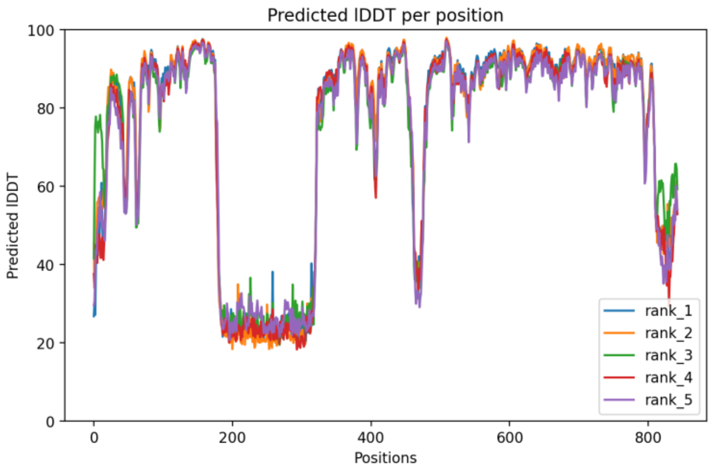


#### Predicted Distogram

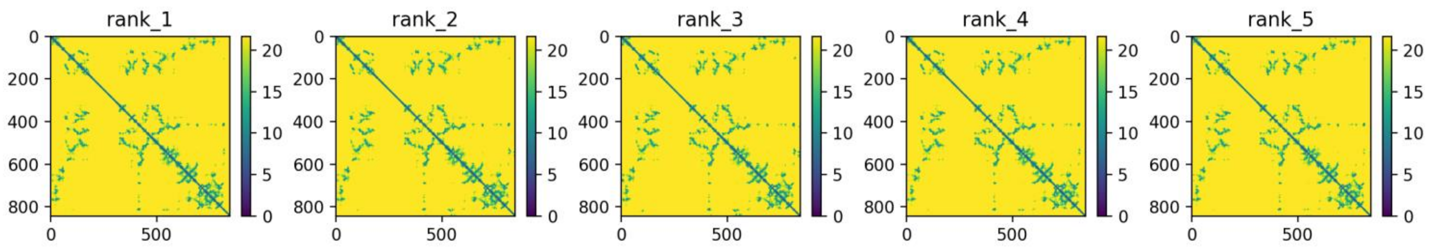


**Legend to Figs. S1-S23.** Images of all models and key quality metrics for the models. The predicted protein folds for all models are displayed in common orientations and color schemes. The color pallet used is: N-terminus, cyan; TP domain, red; T3 motif, violet; Spacer domain, grey; RT1 motif, light blue; RT domain, yellow; RNaseH domain, green; Priming residue, YMDD/YADD/YVDD motifs, D-E-D-D and D-E-E-D motifs, blue spheres; and Aspartate proteinase, magenta. Not all domains/motifs are found in all models. The predicted protein fold is shown the *Predicted Fold* panel. Depth of the multiples sequence alignment used to identify the coordinated variations from which the distance constraints are derived are in the *MSA Coverage* panel. The predicted LDDT value as a function of residue position is in the *Predicted LDDT* panel. Distograms based on the coordinated variations for each of the five models generated for each sequence are found in the *Predicted Distogram* panel.

### Fig. S9. HBV P, Genotype C, Isolate AB560661

#### Predicted fold

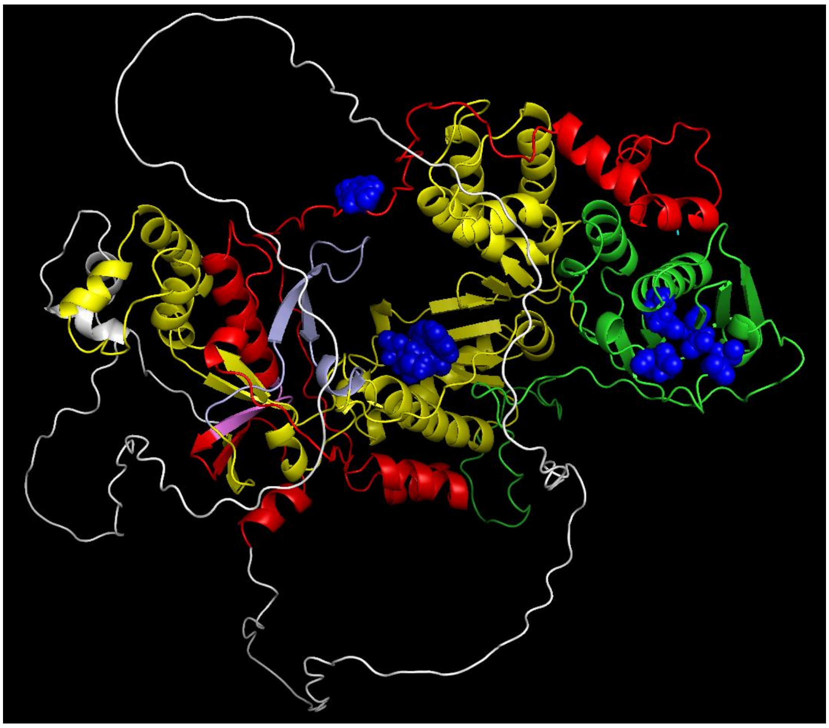

N-terminus, cyan; TP, red; T3 motif, violet; Spacer, grey; RT1 motif, light blue; RT, yellow; RNaseH, green; Y63, YMDD and D-E-D-D, blue

#### MSA coverage

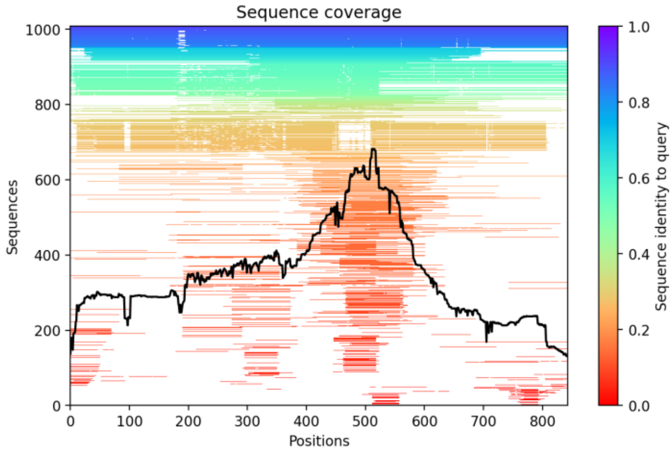


#### Predicted LDDT

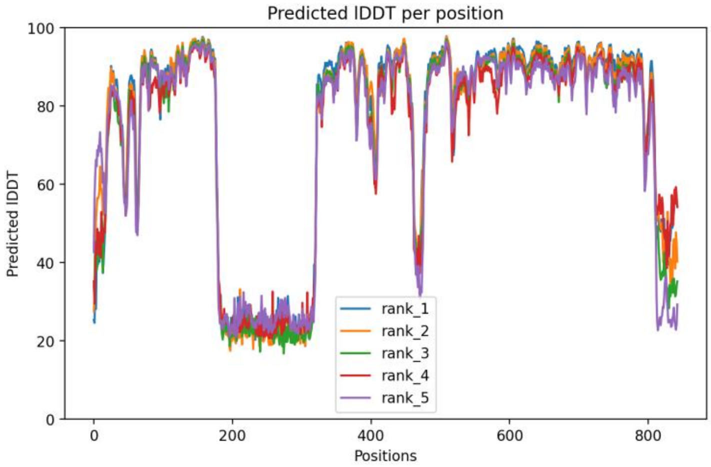


#### Predicted Distogram

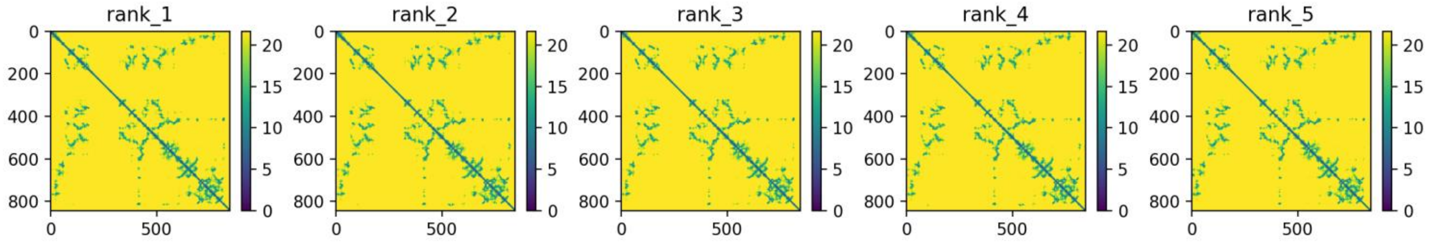


**Legend to Figs. S1-S23.** Images of all models and key quality metrics for the models. The predicted protein folds for all models are displayed in common orientations and color schemes. The color pallet used is: N-terminus, cyan; TP domain, red; T3 motif, violet; Spacer domain, grey; RT1 motif, light blue; RT domain, yellow; RNaseH domain, green; Priming residue, YMDD/YADD/YVDD motifs, D-E-D-D and D-E-E-D motifs, blue spheres; and Aspartate proteinase, magenta. Not all domains/motifs are found in all models. The predicted protein fold is shown the *Predicted Fold* panel. Depth of the multiples sequence alignment used to identify the coordinated variations from which the distance constraints are derived are in the *MSA Coverage* panel. The predicted LDDT value as a function of residue position is in the *Predicted LDDT* panel. Distograms based on the coordinated variations for each of the five models generated for each sequence are found in the *Predicted Distogram* panel.

### Fig. S10. HBV P, Genotype D, Isolate AB554016

#### Predicted fold

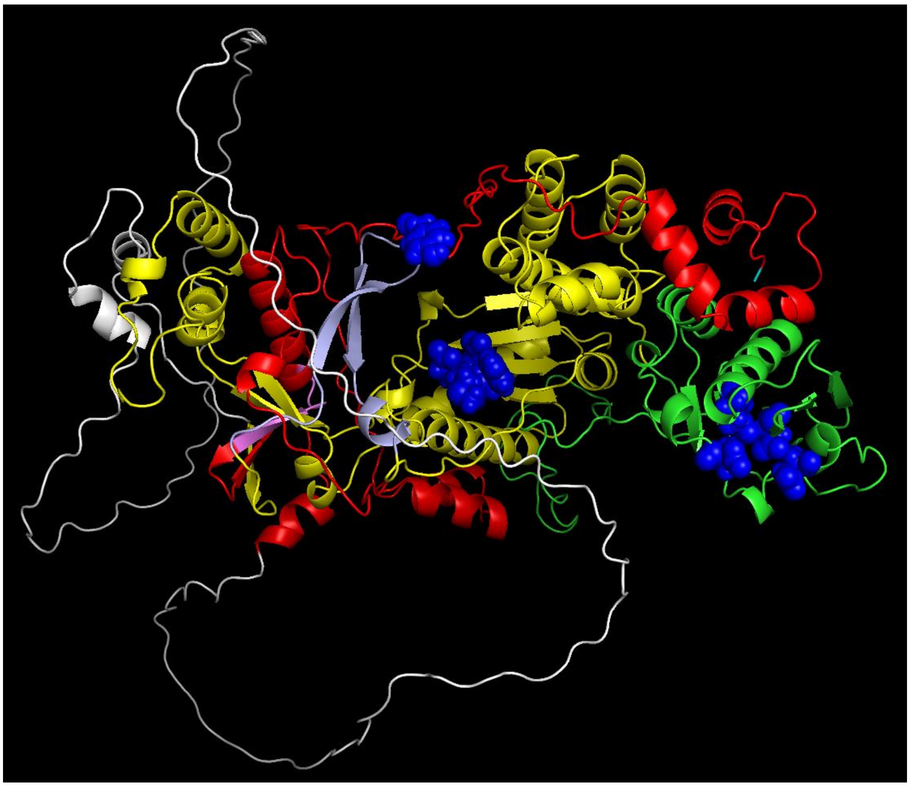

N-terminus, cyan; TP, red; T3 motif, violet; Spacer, grey; RT1 motif, light blue; RT, yellow; RNaseH, green; Y63, YMDD and D-E-D-D, blue

#### MSA coverage

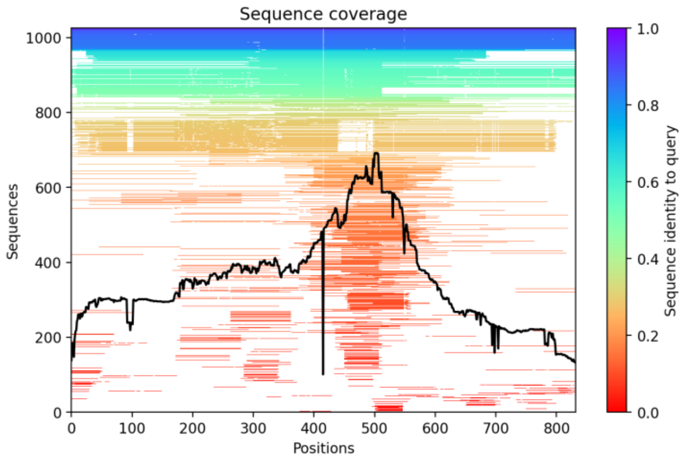


#### Predicted LDDT

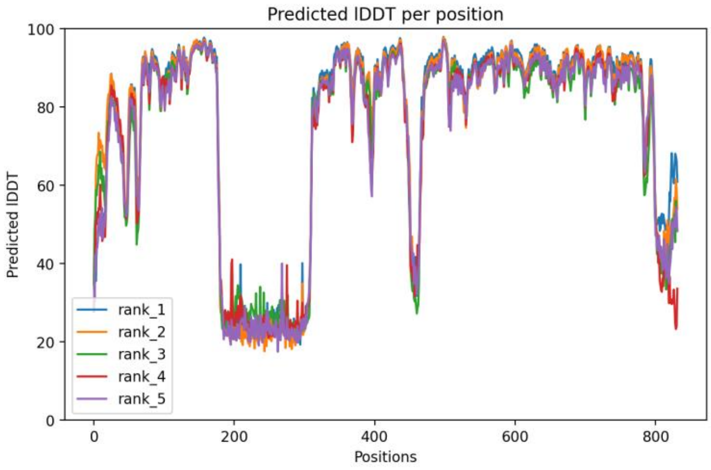


#### Predicted Distogram

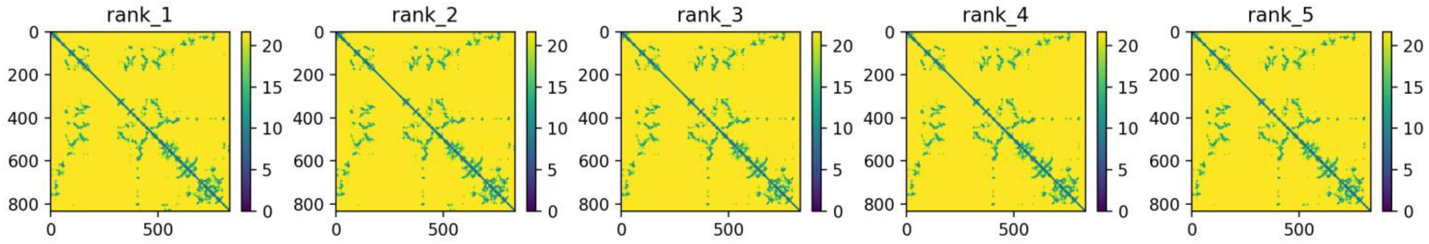


**Legend to Figs. S1-S23.** Images of all models and key quality metrics for the models. The predicted protein folds for all models are displayed in common orientations and color schemes. The color pallet used is: N-terminus, cyan; TP domain, red; T3 motif, violet; Spacer domain, grey; RT1 motif, light blue; RT domain, yellow; RNaseH domain, green; Priming residue, YMDD/YADD/YVDD motifs, D-E-D-D and D-E-E-D motifs, blue spheres; and Aspartate proteinase, magenta. Not all domains/motifs are found in all models. The predicted protein fold is shown the *Predicted Fold* panel. Depth of the multiples sequence alignment used to identify the coordinated variations from which the distance constraints are derived are in the *MSA Coverage* panel. The predicted LDDT value as a function of residue position is in the *Predicted LDDT* panel. Distograms based on the coordinated variations for each of the five models generated for each sequence are found in the *Predicted Distogram* panel.

### Fig. S11. HBV P, Genotype E, Isolate AB091255

#### Predicted fold

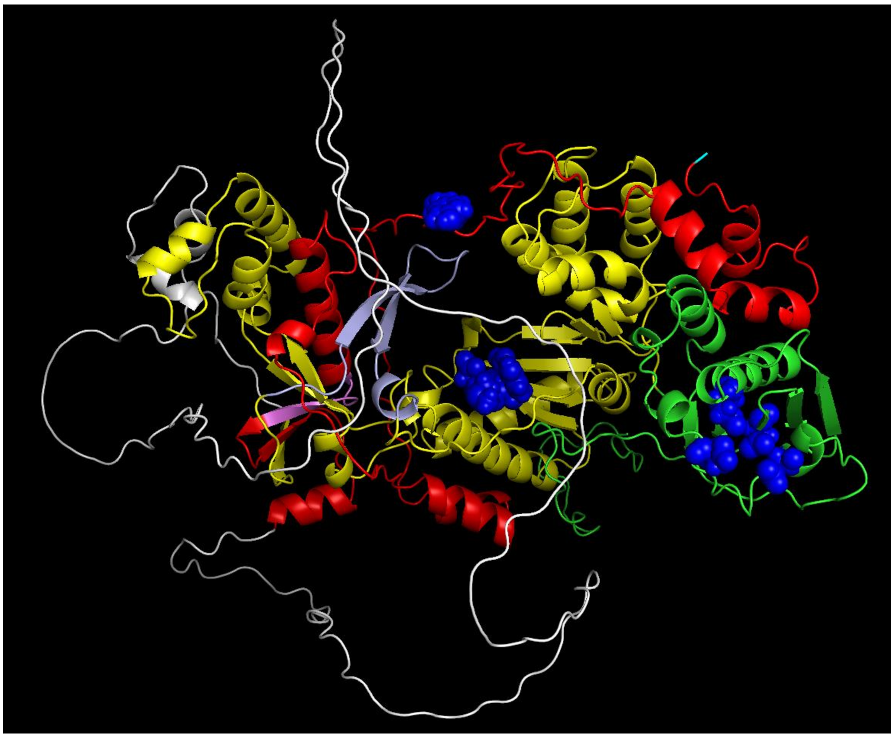

N-terminus, cyan; TP, red; T3 motif, violet; Spacer, grey; RT1 motif, light blue; RT, yellow; RNaseH, green; Y63, YMDD and D-E-D-D, blue

#### MSA coverage

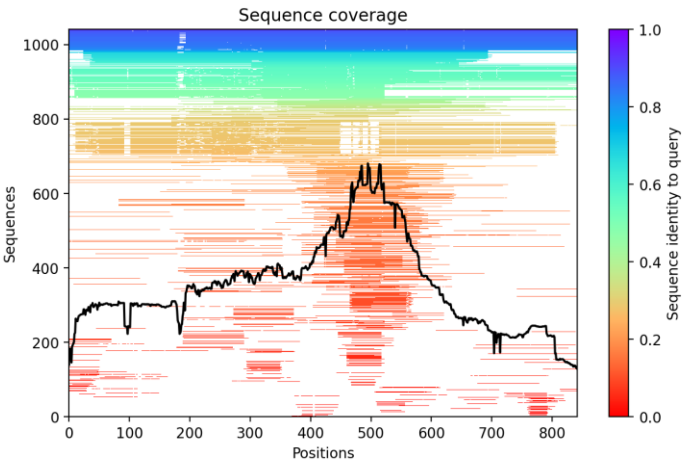


#### Predicted LDDT

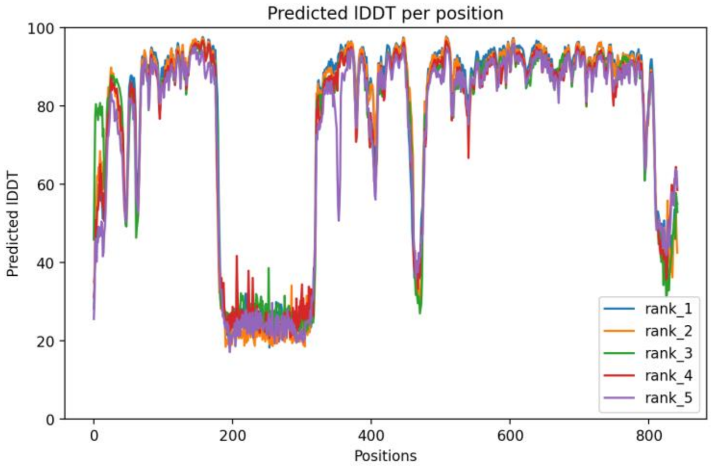


#### Predicted Distogram

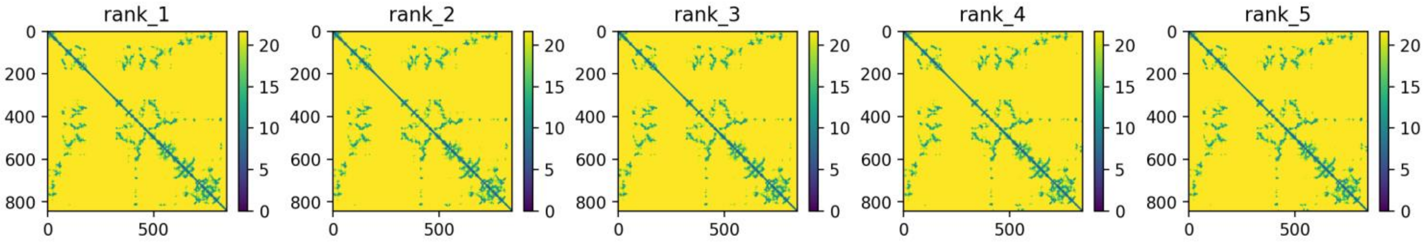


**Legend to Figs. S1-S23.** Images of all models and key quality metrics for the models. The predicted protein folds for all models are displayed in common orientations and color schemes. The color pallet used is: N-terminus, cyan; TP domain, red; T3 motif, violet; Spacer domain, grey; RT1 motif, light blue; RT domain, yellow; RNaseH domain, green; Priming residue, YMDD/YADD/YVDD motifs, D-E-D-D and D-E-E-D motifs, blue spheres; and Aspartate proteinase, magenta. Not all domains/motifs are found in all models. The predicted protein fold is shown the *Predicted Fold* panel. Depth of the multiples sequence alignment used to identify the coordinated variations from which the distance constraints are derived are in the *MSA Coverage* panel. The predicted LDDT value as a function of residue position is in the *Predicted LDDT* panel. Distograms based on the coordinated variations for each of the five models generated for each sequence are found in the *Predicted Distogram* panel.

### Fig. S12. HBV P, Genotype F, Isolate AB064316

#### Predicted fold

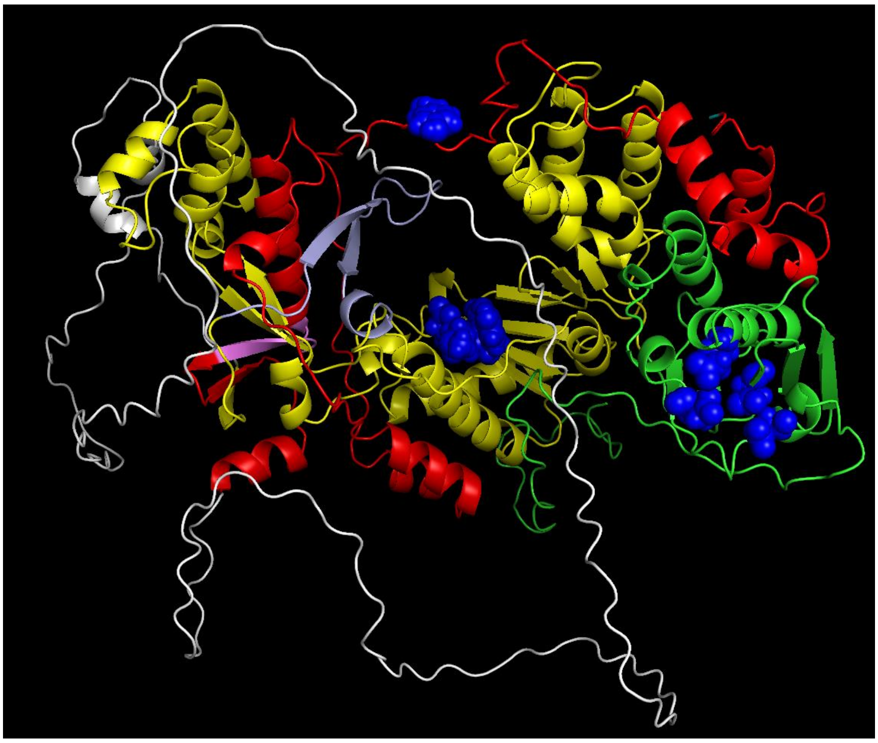

N-terminus, cyan; TP, red; T3 motif, violet; Spacer, grey; RT1 motif, light blue; RT, yellow; RNaseH, green; Y63, YMDD and D-E-D-D, blue

#### MSA coverage

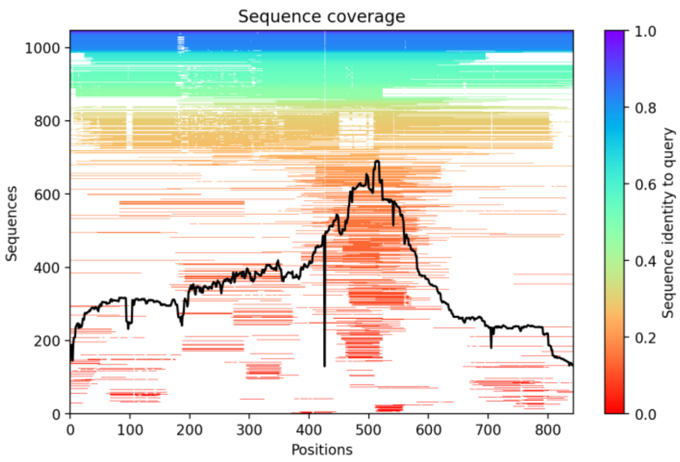


#### Predicted LDDT

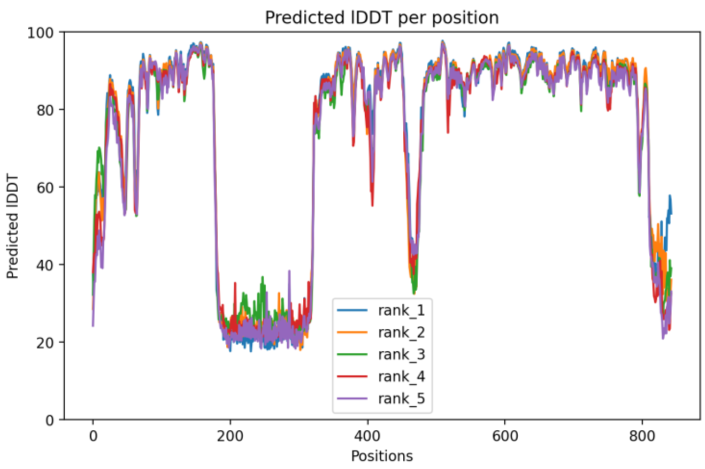


#### Predicted Distogram

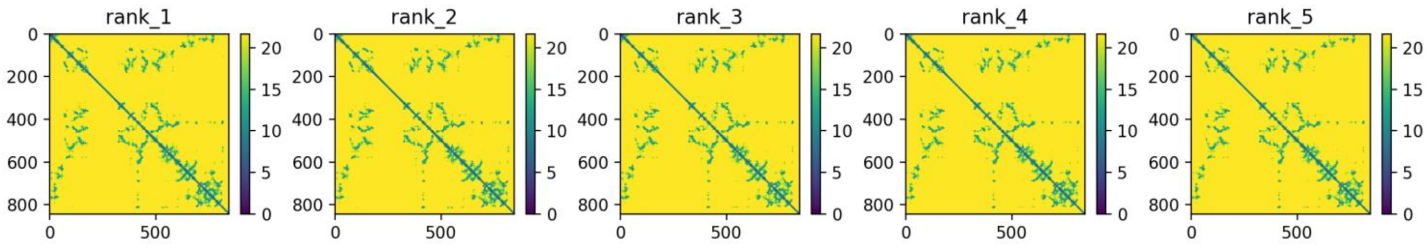


**Legend to Figs. S1-S23.** Images of all models and key quality metrics for the models. The predicted protein folds for all models are displayed in common orientations and color schemes. The color pallet used is: N-terminus, cyan; TP domain, red; T3 motif, violet; Spacer domain, grey; RT1 motif, light blue; RT domain, yellow; RNaseH domain, green; Priming residue, YMDD/YADD/YVDD motifs, D-E-D-D and D-E-E-D motifs, blue spheres; and Aspartate proteinase, magenta. Not all domains/motifs are found in all models. The predicted protein fold is shown the *Predicted Fold* panel. Depth of the multiples sequence alignment used to identify the coordinated variations from which the distance constraints are derived are in the *MSA Coverage* panel. The predicted LDDT value as a function of residue position is in the *Predicted LDDT* panel. Distograms based on the coordinated variations for each of the five models generated for each sequence are found in the *Predicted Distogram* panel.

### Fig. S13. HBV P, Genotype G, Isolate AB064310

#### Predicted fold

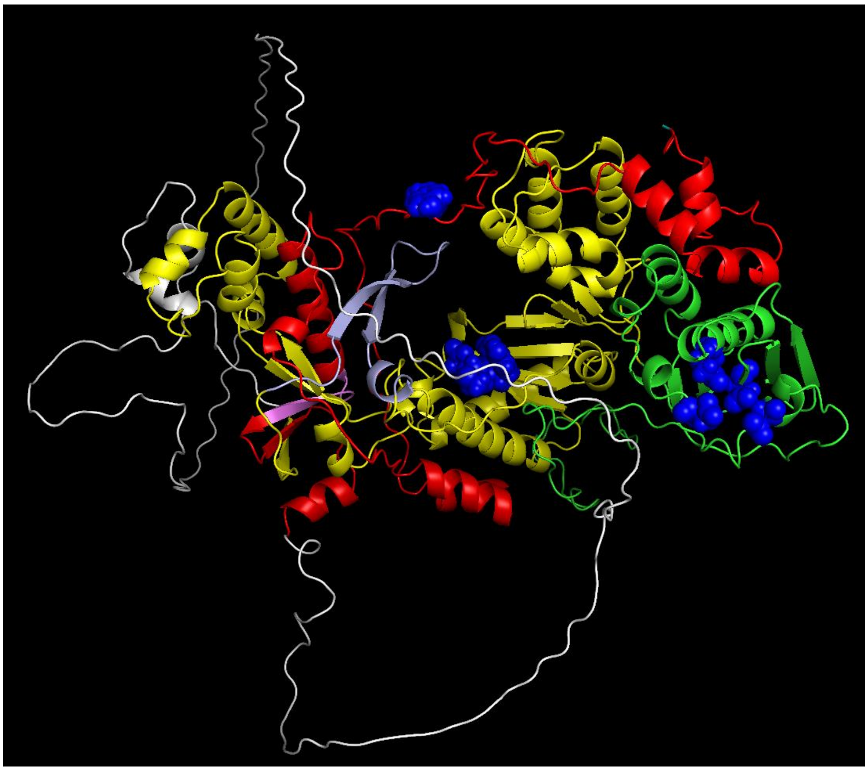

N-terminus, cyan; TP, red; T3 motif, violet; Spacer, grey; RT1 motif, light blue; RT, yellow; RNaseH, green; Y63, YMDD and D-E-D-D, blue

#### MSA coverage

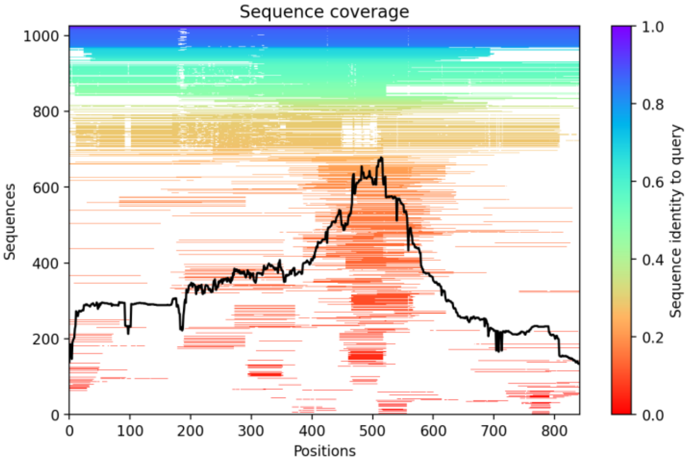


#### Predicted LDDT

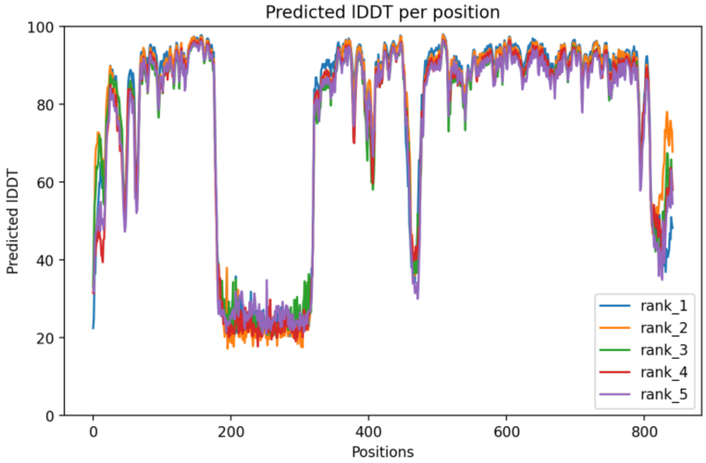


#### Predicted Distogram

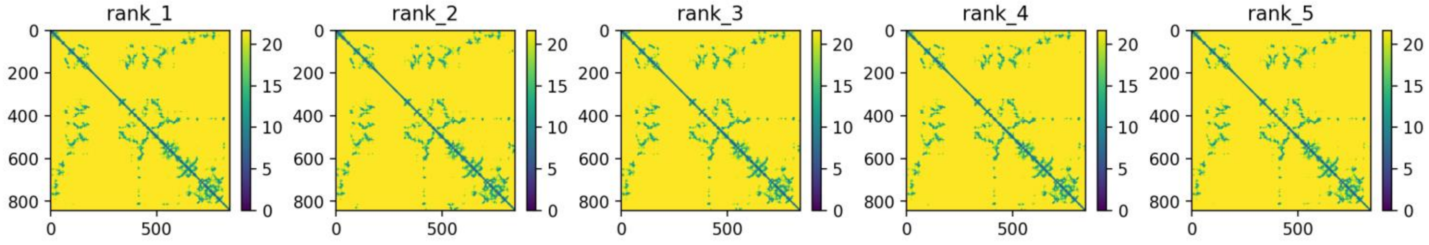


**Legend to Figs. S1-S23.** Images of all models and key quality metrics for the models. The predicted protein folds for all models are displayed in common orientations and color schemes. The color pallet used is: N-terminus, cyan; TP domain, red; T3 motif, violet; Spacer domain, grey; RT1 motif, light blue; RT domain, yellow; RNaseH domain, green; Priming residue, YMDD/YADD/YVDD motifs, D-E-D-D and D-E-E-D motifs, blue spheres; and Aspartate proteinase, magenta. Not all domains/motifs are found in all models. The predicted protein fold is shown the *Predicted Fold* panel. Depth of the multiples sequence alignment used to identify the coordinated variations from which the distance constraints are derived are in the *MSA Coverage* panel. The predicted LDDT value as a function of residue position is in the *Predicted LDDT* panel. Distograms based on the coordinated variations for each of the five models generated for each sequence are found in the *Predicted Distogram* panel.

### Fig. S14. HBV P, Genotype H, Isolate AB298362

#### Predicted fold

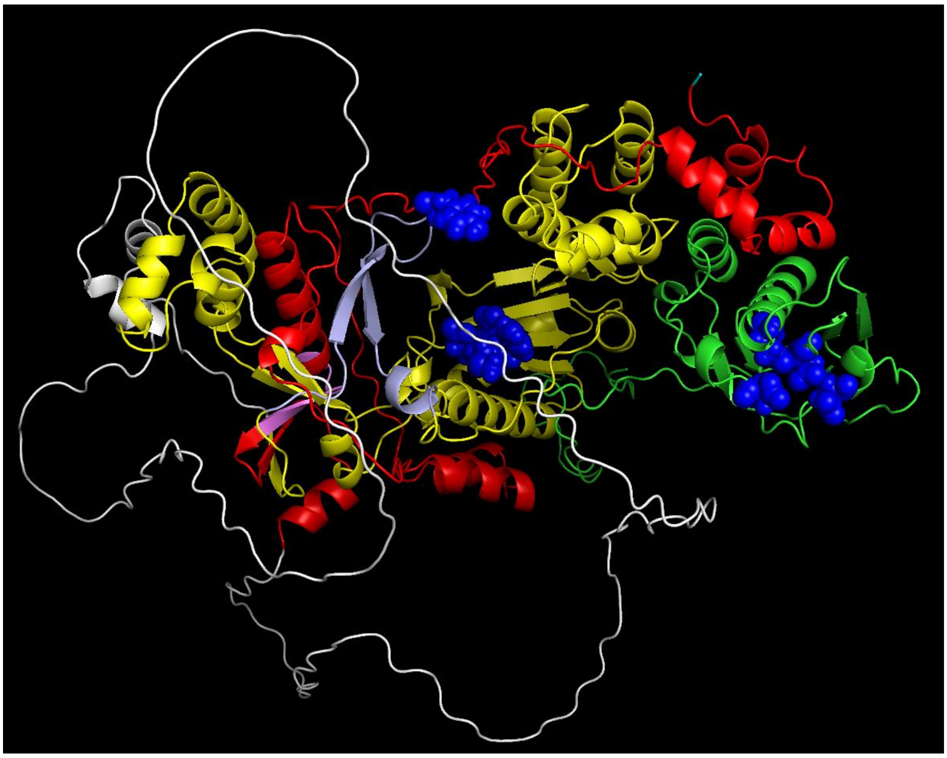

N-terminus, cyan; TP, red; T3 motif, violet; Spacer, grey; RT1 motif, light blue; RT, yellow; RNaseH, green; Y63, YMDD and D-E-D-D, blue

#### MSA coverage

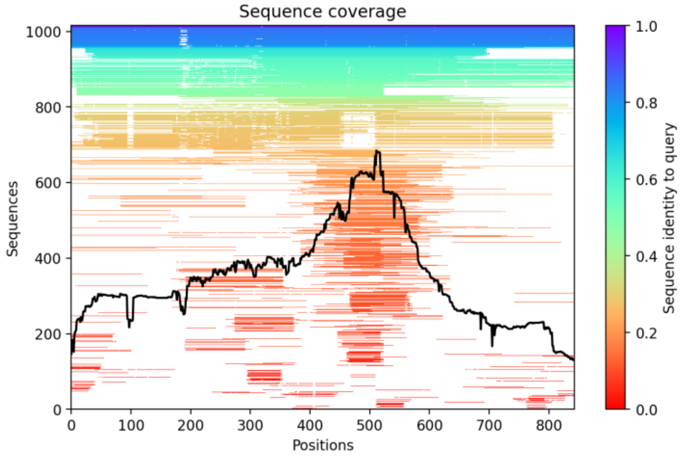


#### Predicted LDDT

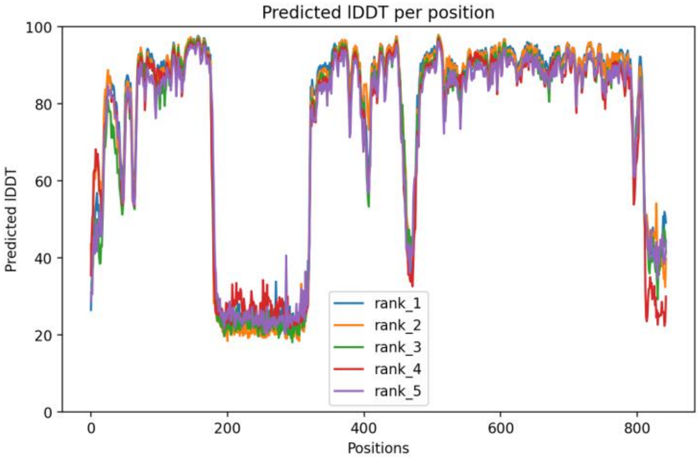


#### Predicted Distogram

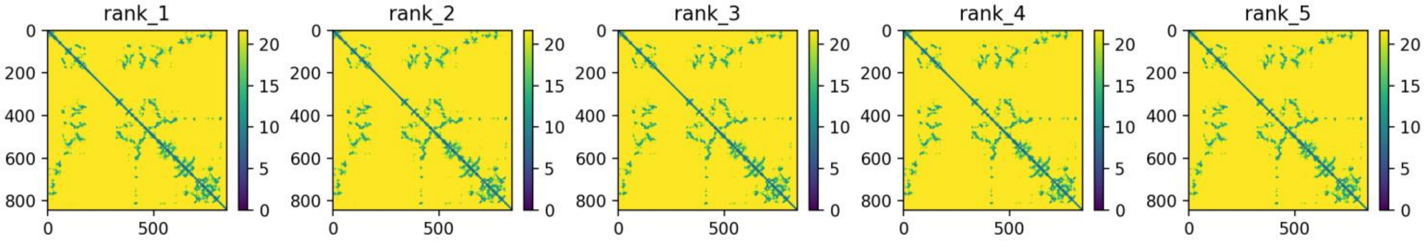


**Legend to Figs. S1-S23.** Images of all models and key quality metrics for the models. The predicted protein folds for all models are displayed in common orientations and color schemes. The color pallet used is: N-terminus, cyan; TP domain, red; T3 motif, violet; Spacer domain, grey; RT1 motif, light blue; RT domain, yellow; RNaseH domain, green; Priming residue, YMDD/YADD/YVDD motifs, D-E-D-D and D-E-E-D motifs, blue spheres; and Aspartate proteinase, magenta. Not all domains/motifs are found in all models. The predicted protein fold is shown the *Predicted Fold* panel. Depth of the multiples sequence alignment used to identify the coordinated variations from which the distance constraints are derived are in the *MSA Coverage* panel. The predicted LDDT value as a function of residue position is in the *Predicted LDDT* panel. Distograms based on the coordinated variations for each of the five models generated for each sequence are found in the *Predicted Distogram* panel.

### Fig. S15. HBV P, Genotype I, Isolate MH368022

#### Predicted fold

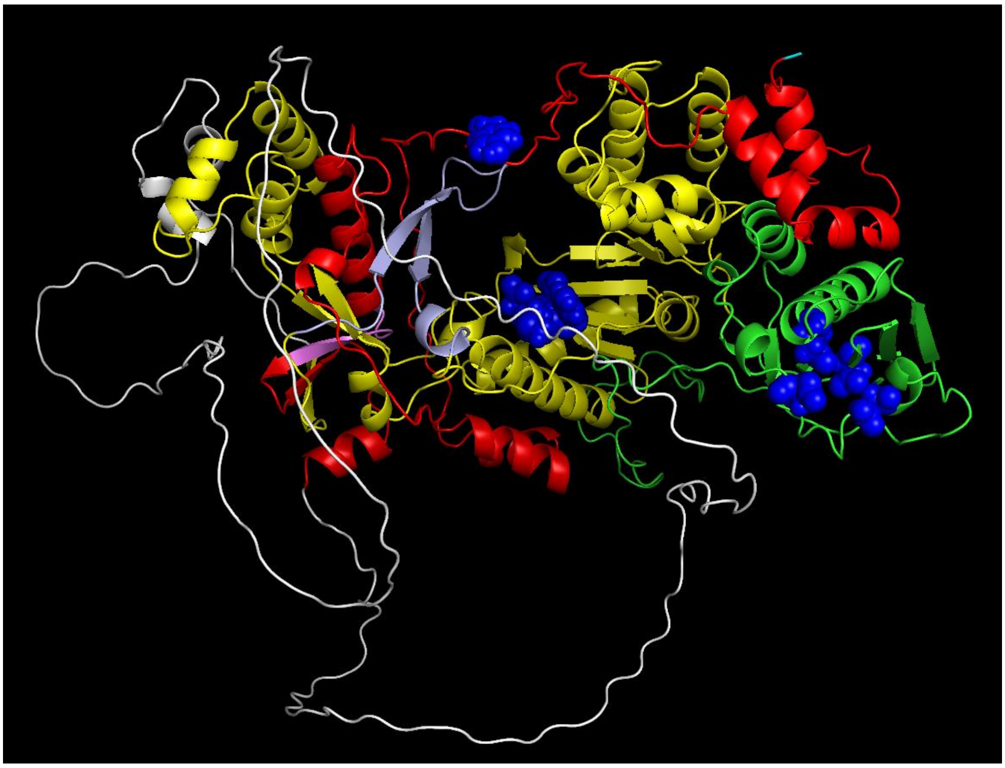

N-terminus, cyan; TP, red; T3 motif, violet; Spacer, grey; RT1 motif, light blue; RT, yellow; RNaseH, green; Y63, YMDD and D-E-D-D, blue

#### MSA coverage

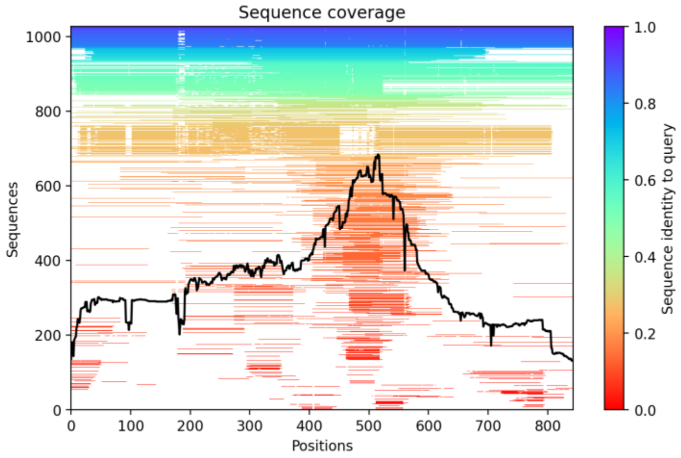


#### Predicted LDDT

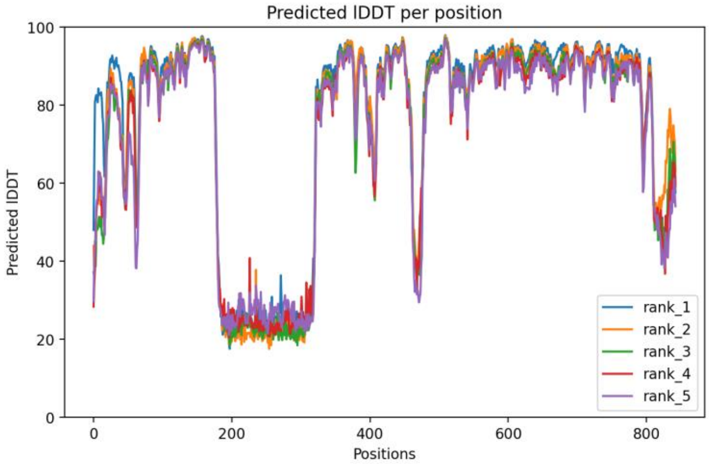


#### Predicted Distogram

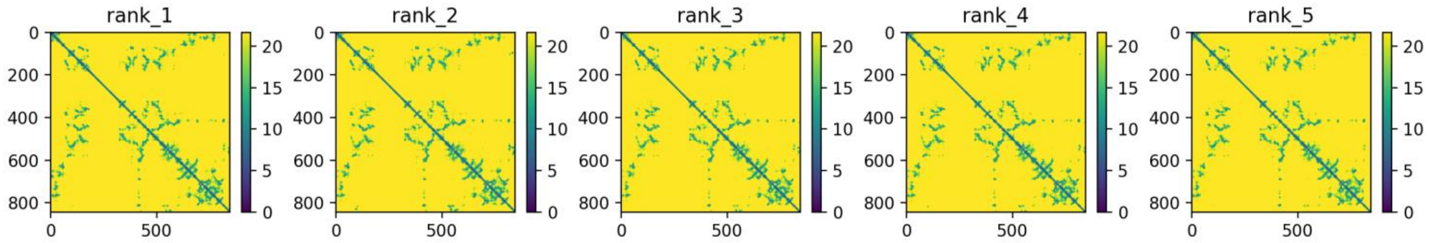


**Legend to Figs. S1-S23.** Images of all models and key quality metrics for the models. The predicted protein folds for all models are displayed in common orientations and color schemes. The color pallet used is: N-terminus, cyan; TP domain, red; T3 motif, violet; Spacer domain, grey; RT1 motif, light blue; RT domain, yellow; RNaseH domain, green; Priming residue, YMDD/YADD/YVDD motifs, D-E-D-D and D-E-E-D motifs, blue spheres; and Aspartate proteinase, magenta. Not all domains/motifs are found in all models. The predicted protein fold is shown the *Predicted Fold* panel. Depth of the multiples sequence alignment used to identify the coordinated variations from which the distance constraints are derived are in the *MSA Coverage* panel. The predicted LDDT value as a function of residue position is in the *Predicted LDDT* panel. Distograms based on the coordinated variations for each of the five models generated for each sequence are found in the *Predicted Distogram* panel.

### Fig. S16. WHV P, Isolate NC_004107

#### Predicted fold

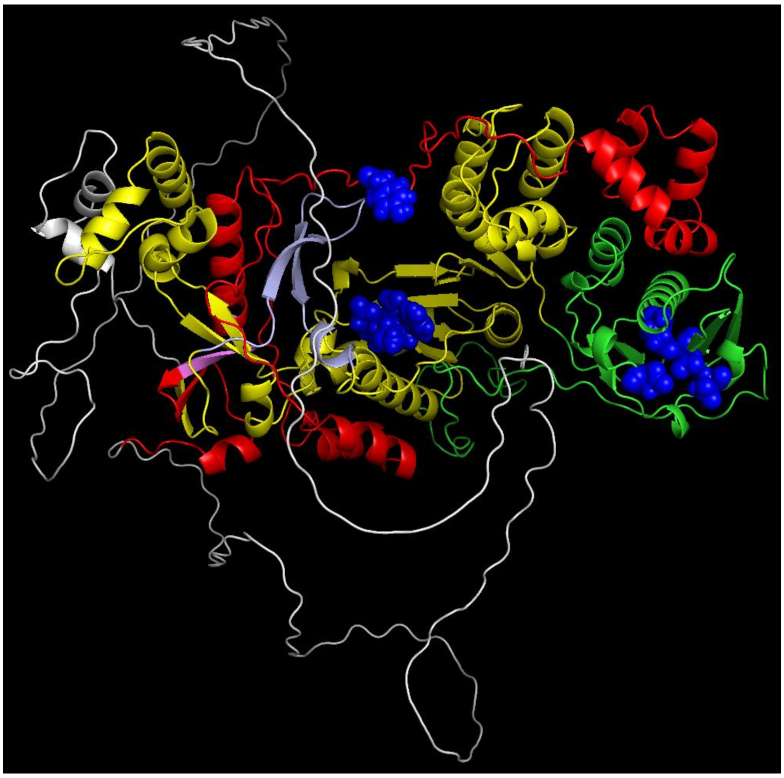

N-terminus, cyan; TP, red; T3 motif, violet; Spacer, grey; RT1 motif, light blue; RT, yellow; RNaseH, green; Y63, YMDD and D-E-D-D, blue

#### MSA coverage

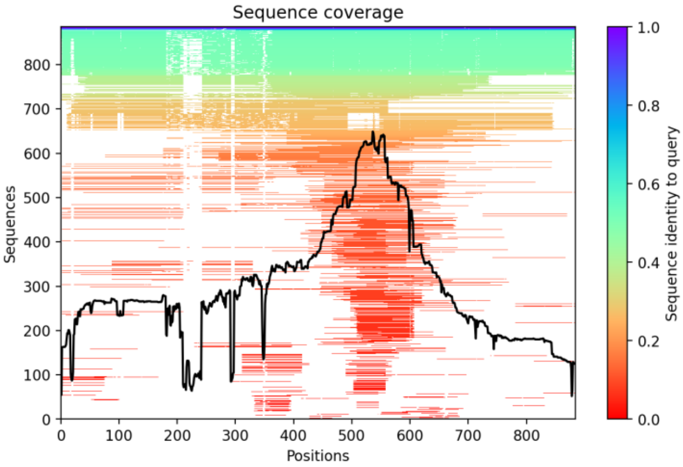


#### Predicted LDDT

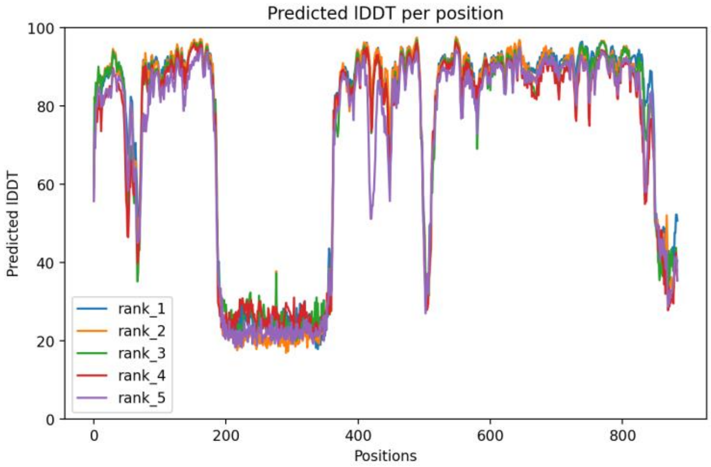


#### Predicted Distogram

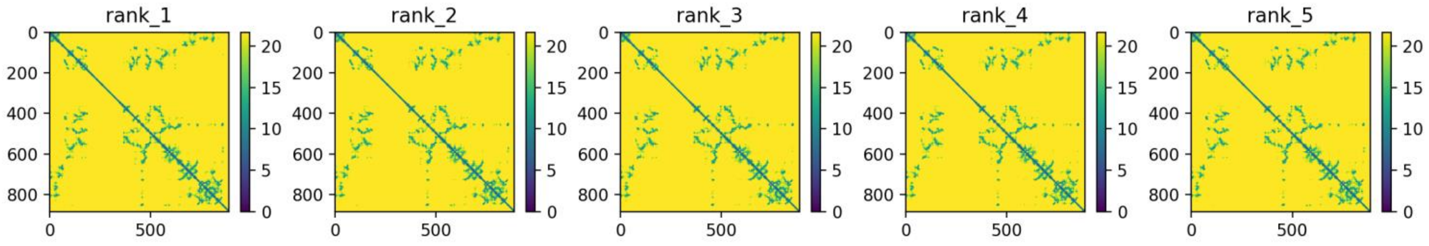


**Legend to Figs. S1-S23.** Images of all models and key quality metrics for the models. The predicted protein folds for all models are displayed in common orientations and color schemes. The color pallet used is: N-terminus, cyan; TP domain, red; T3 motif, violet; Spacer domain, grey; RT1 motif, light blue; RT domain, yellow; RNaseH domain, green; Priming residue, YMDD/YADD/YVDD motifs, D-E-D-D and D-E-E-D motifs, blue spheres; and Aspartate proteinase, magenta. Not all domains/motifs are found in all models. The predicted protein fold is shown the *Predicted Fold* panel. Depth of the multiples sequence alignment used to identify the coordinated variations from which the distance constraints are derived are in the *MSA Coverage* panel. The predicted LDDT value as a function of residue position is in the *Predicted LDDT* panel. Distograms based on the coordinated variations for each of the five models generated for each sequence are found in the *Predicted Distogram* panel.

### Fig. S17. DHBV P, Strain 3, Isolate DQ195079

#### Predicted fold

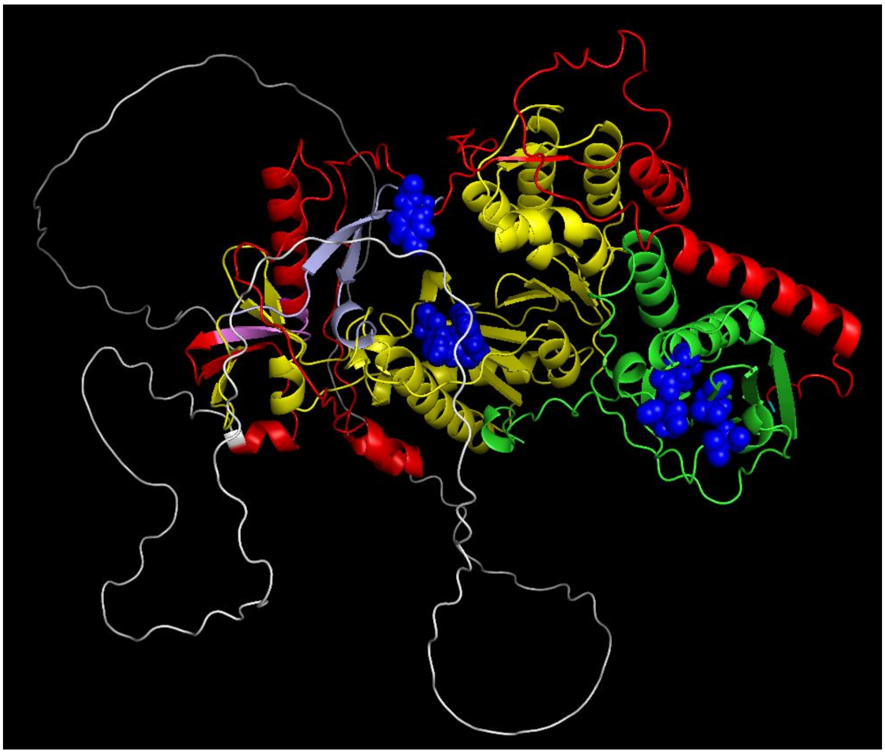

N-terminus, cyan; TP, red; T3 motif, violet; Spacer, grey; RT1 motif, light blue; RT, yellow; RNaseH, green; Y63, YMDD and D-E-D-D, blue

#### MSA coverage

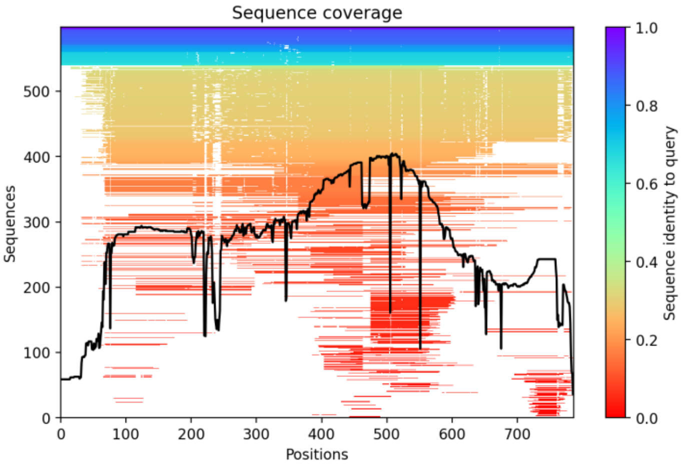


#### Predicted LDDT

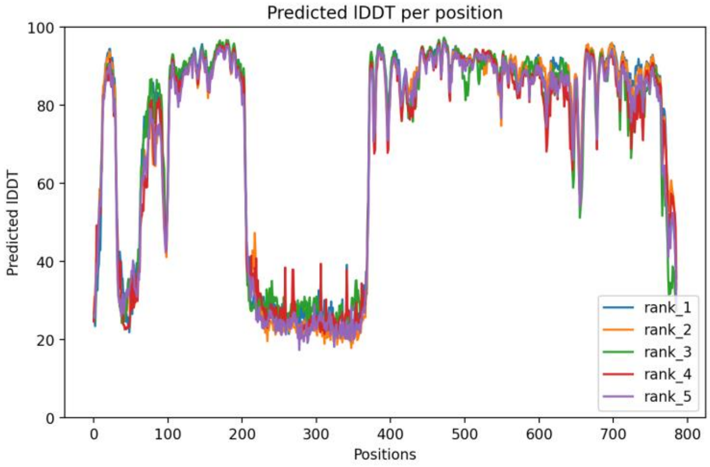


#### Predicted Distogram

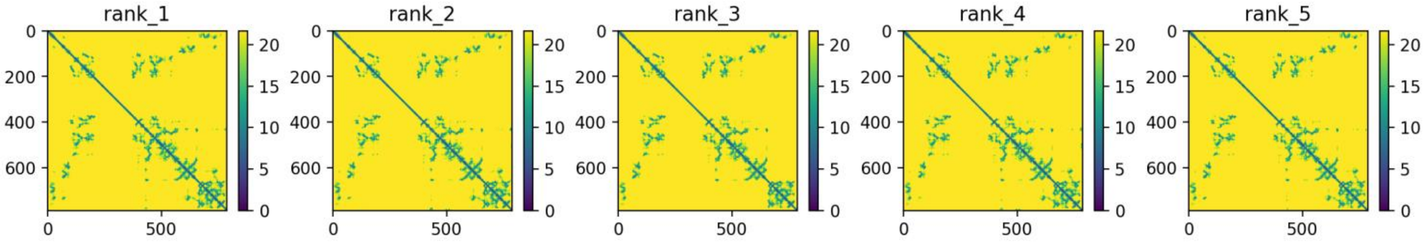


**Legend to Figs. S1-S23.** Images of all models and key quality metrics for the models. The predicted protein folds for all models are displayed in common orientations and color schemes. The color pallet used is: N-terminus, cyan; TP domain, red; T3 motif, violet; Spacer domain, grey; RT1 motif, light blue; RT domain, yellow; RNaseH domain, green; Priming residue, YMDD/YADD/YVDD motifs, D-E-D-D and D-E-E-D motifs, blue spheres; and Aspartate proteinase, magenta. Not all domains/motifs are found in all models. The predicted protein fold is shown the *Predicted Fold* panel. Depth of the multiples sequence alignment used to identify the coordinated variations from which the distance constraints are derived are in the *MSA Coverage* panel. The predicted LDDT value as a function of residue position is in the *Predicted LDDT* panel. Distograms based on the coordinated variations for each of the five models generated for each sequence are found in the *Predicted Distogram* panel.

**Fig. S18. SkHBV P** (Lauber et al. Cell Host & Microbe, 2017, 22:387)

Predicted fold

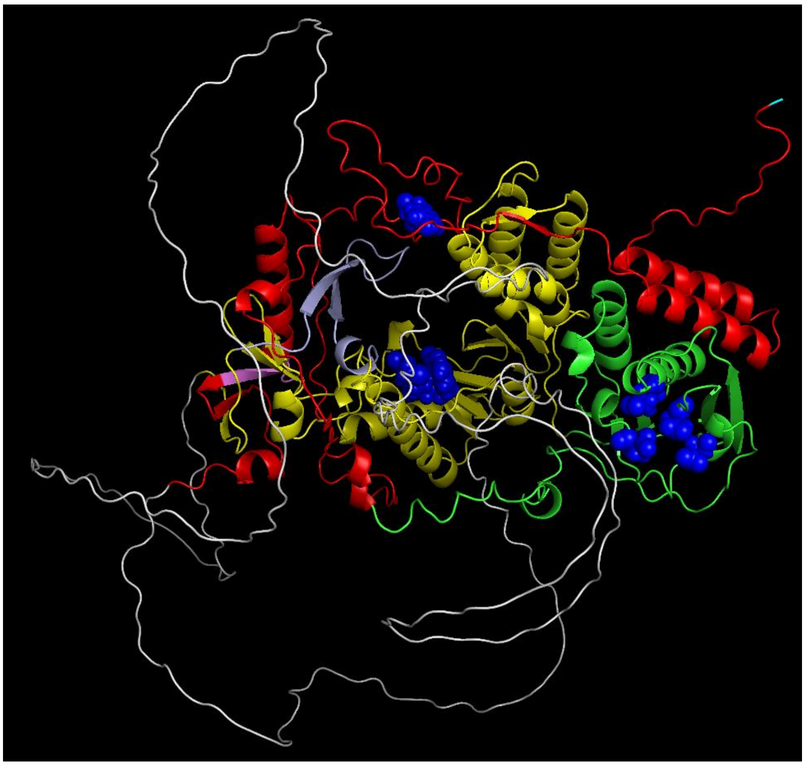

N-terminus, cyan; TP, red; T3 motif, violet; Spacer, grey; RT1 motif, light blue; RT, yellow; RNaseH, green; Y63, YMDD and D-E-D-D, blue

MSA coverage

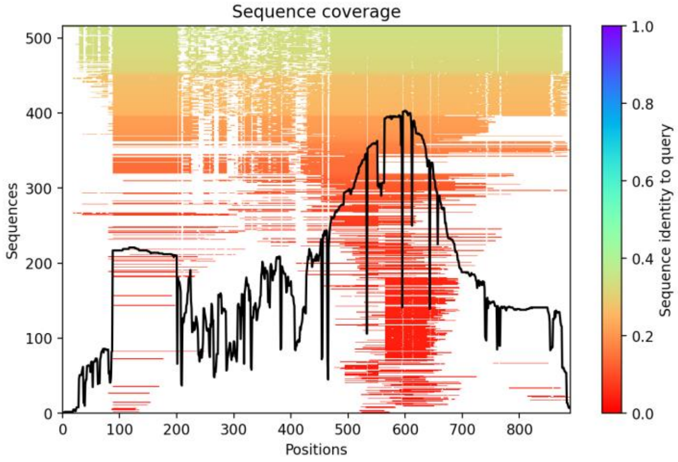


Predicted LDDT

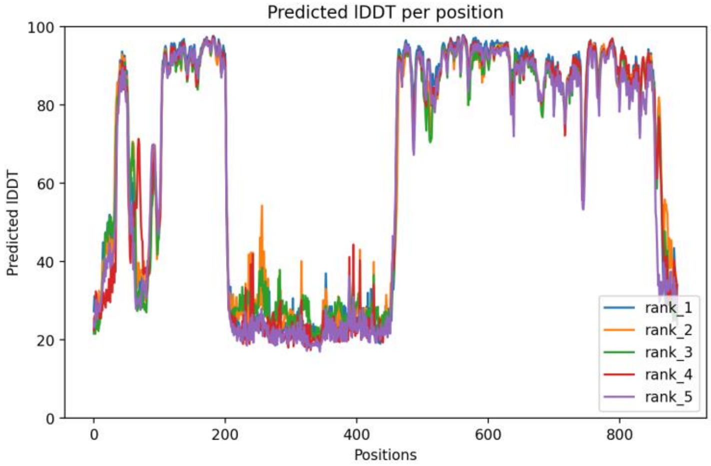


Predicted Distogram

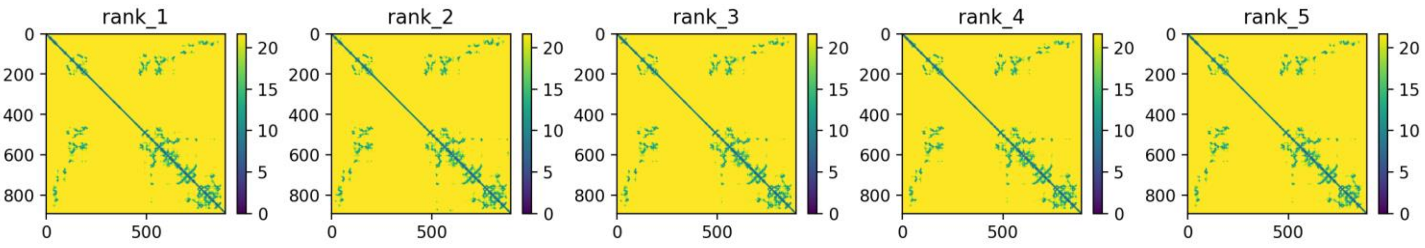


**Legend to Figs. S1-S23.** Images of all models and key quality metrics for the models. The predicted protein folds for all models are displayed in common orientations and color schemes. The color pallet used is: N-terminus, cyan; TP domain, red; T3 motif, violet; Spacer domain, grey; RT1 motif, light blue; RT domain, yellow; RNaseH domain, green; Priming residue, YMDD/YADD/YVDD motifs, D-E-D-D and D-E-E-D motifs, blue spheres; and Aspartate proteinase, magenta. Not all domains/motifs are found in all models. The predicted protein fold is shown the *Predicted Fold* panel. Depth of the multiples sequence alignment used to identify the coordinated variations from which the distance constraints are derived are in the *MSA Coverage* panel. The predicted LDDT value as a function of residue position is in the *Predicted LDDT* panel. Distograms based on the coordinated variations for each of the five models generated for each sequence are found in the *Predicted Distogram* panel.

#### **Fig. S19. TFHBV P** (Lauber et al. Cell Host & Microbe, 2017, 22:387)

Predicted fold

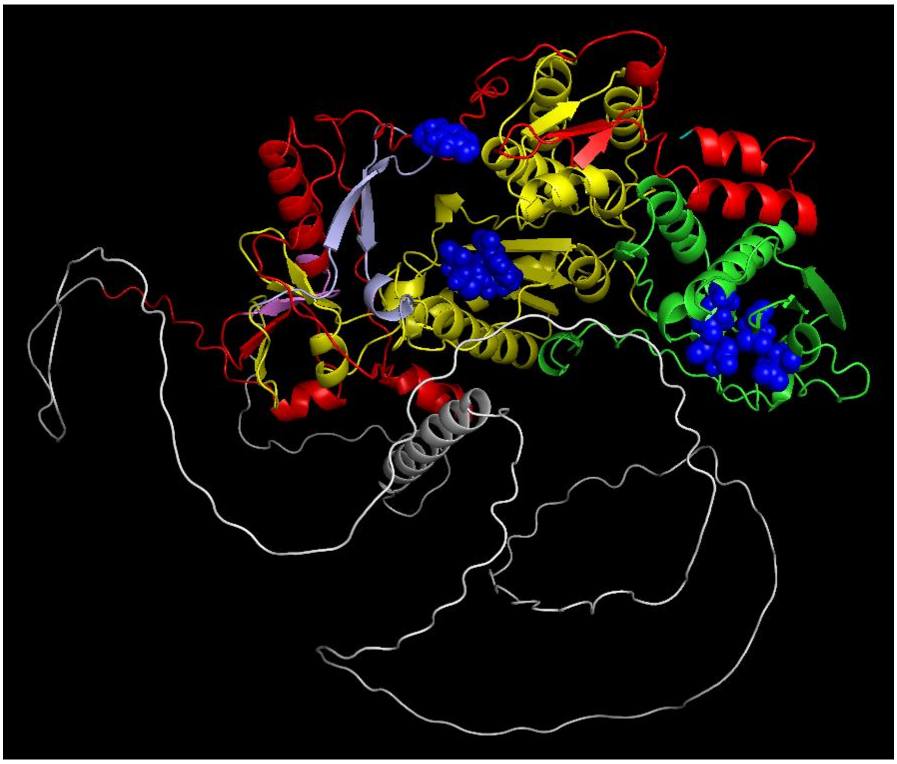

N-terminus, cyan; TP, red; T3 motif, violet; Spacer, grey; RT1 motif, light blue; RT, yellow; RNaseH, green; Y63, YMDD and D-E-D-D, blue

MSA coverage

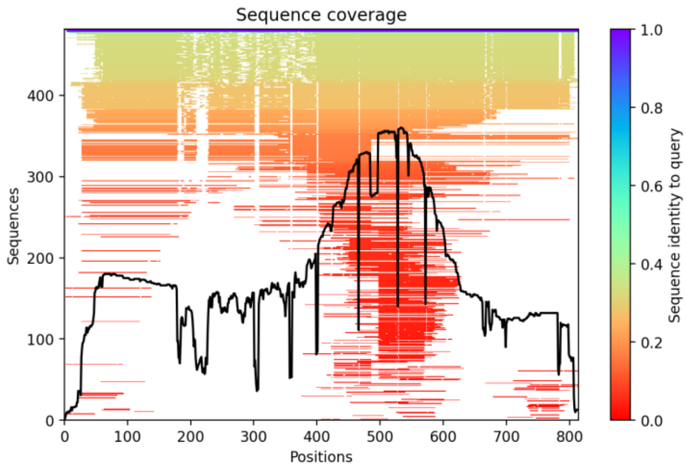


Predicted LDDT

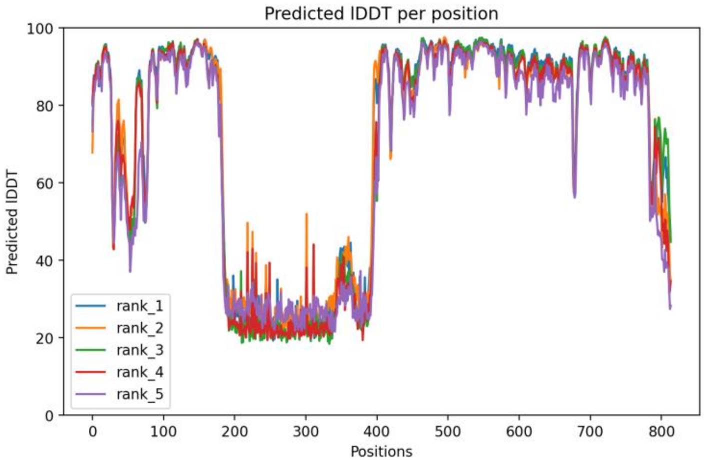


Predicted Distogram

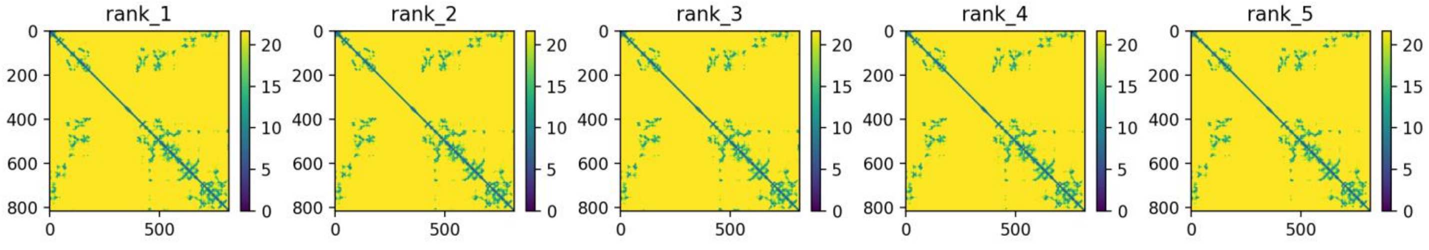


**Legend to Figs. S1-S23.** Images of all models and key quality metrics for the models. The predicted protein folds for all models are displayed in common orientations and color schemes. The color pallet used is: N-terminus, cyan; TP domain, red; T3 motif, violet; Spacer domain, grey; RT1 motif, light blue; RT domain, yellow; RNaseH domain, green; Priming residue, YMDD/YADD/YVDD motifs, D-E-D-D and D-E-E-D motifs, blue spheres; and Aspartate proteinase, magenta. Not all domains/motifs are found in all models. The predicted protein fold is shown the *Predicted Fold* panel. Depth of the multiples sequence alignment used to identify the coordinated variations from which the distance constraints are derived are in the *MSA Coverage* panel. The predicted LDDT value as a function of residue position is in the *Predicted LDDT* panel. Distograms based on the coordinated variations for each of the five models generated for each sequence are found in the *Predicted Distogram* panel.

#### **Fig. S20. TMDV P** (Lauber et al. Cell Host & Microbe, 2017, 22:387)

Predicted fold

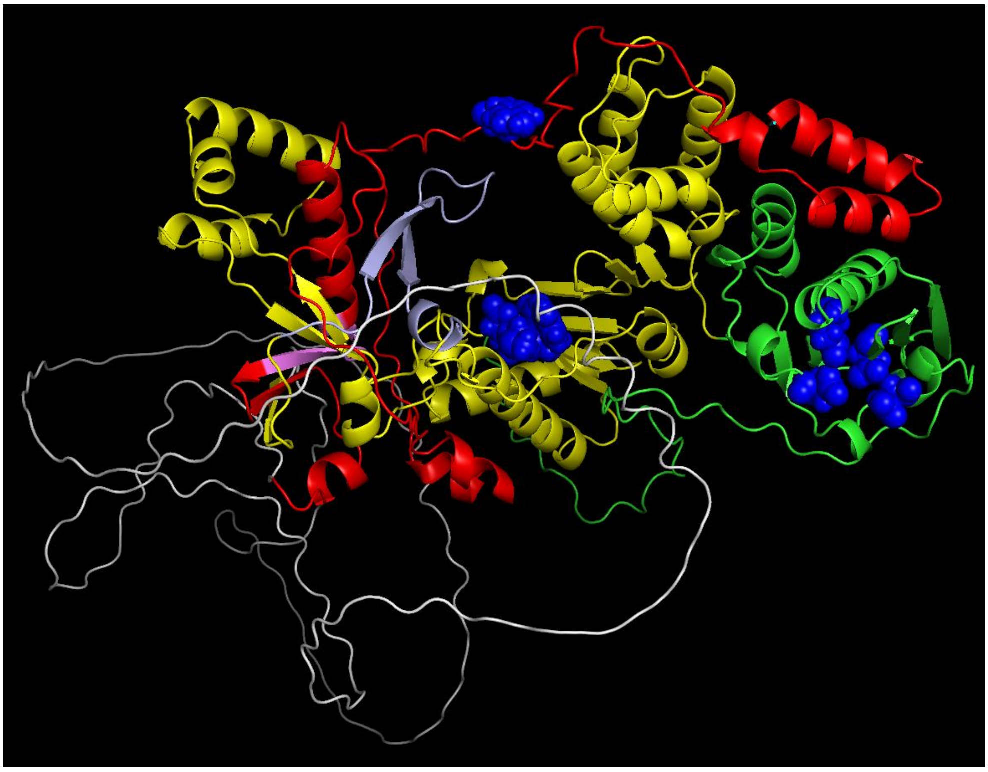

N-terminus, cyan; TP, red; T3 motif, violet; Spacer, grey; RT1 motif, light blue; RT, yellow; RNaseH, green; Y63, YMDD and D-E-D-D, blue

MSA coverage

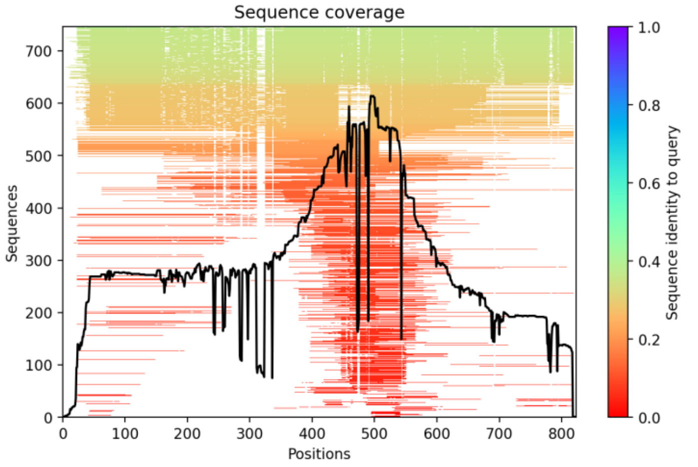


Predicted LDDT

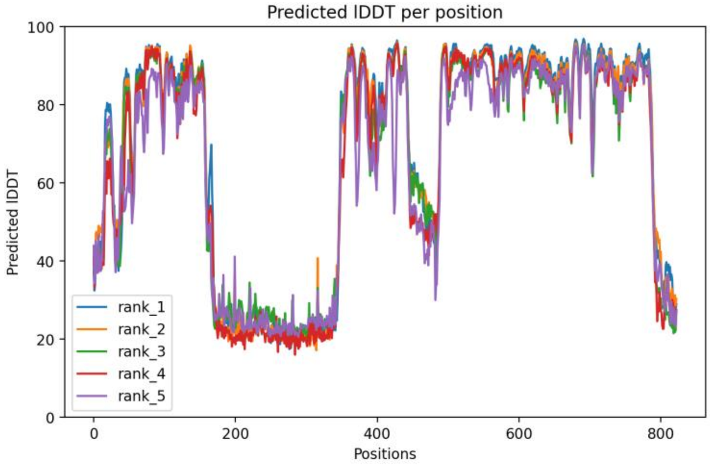


Predicted Distogram

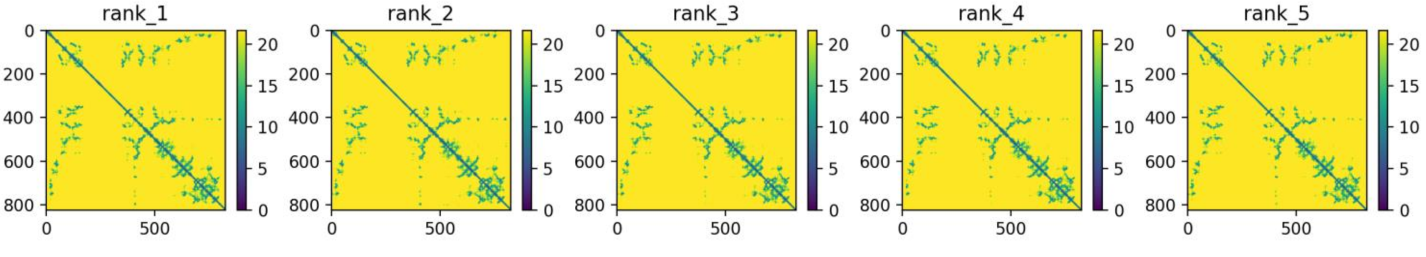


**Legend to Figs. S1-S23.** Images of all models and key quality metrics for the models. The predicted protein folds for all models are displayed in common orientations and color schemes. The color pallet used is: N-terminus, cyan; TP domain, red; T3 motif, violet; Spacer domain, grey; RT1 motif, light blue; RT domain, yellow; RNaseH domain, green; Priming residue, YMDD/YADD/YVDD motifs, D-E-D-D and D-E-E-D motifs, blue spheres; and Aspartate proteinase, magenta. Not all domains/motifs are found in all models. The predicted protein fold is shown the *Predicted Fold* panel. Depth of the multiples sequence alignment used to identify the coordinated variations from which the distance constraints are derived are in the *MSA Coverage* panel. The predicted LDDT value as a function of residue position is in the *Predicted LDDT* panel. Distograms based on the coordinated variations for each of the five models generated for each sequence are found in the *Predicted Distogram* panel.

### Fig. S21. RNDV P (Lauber et al. Cell Host & Microbe, 2017, 22:387)

#### Predicted fold

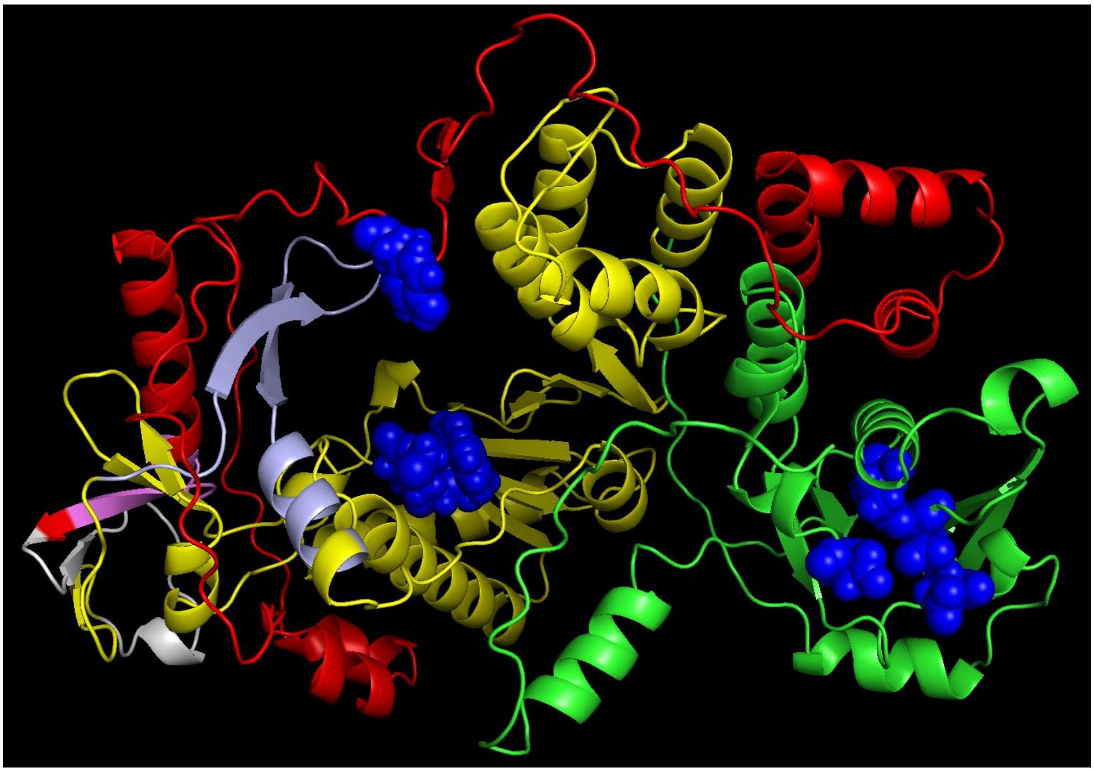

N-terminus, cyan; TP, red; T3 motif, violet; Spacer, grey; RT1 motif, light blue; RT, yellow; RNaseH, green; Y63, YMDD and D-E-D-D, blue

#### MSA coverage

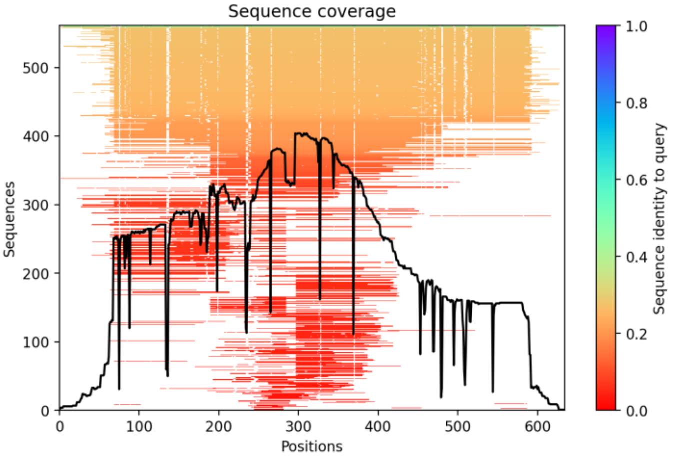


#### Predicted LDDT

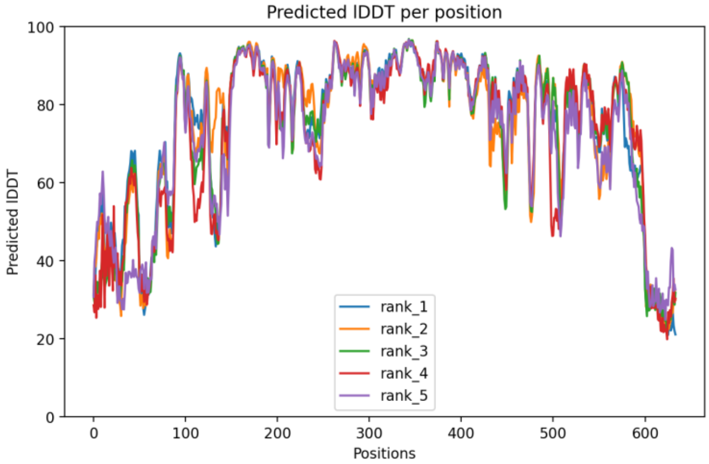


#### Predicted Distogram

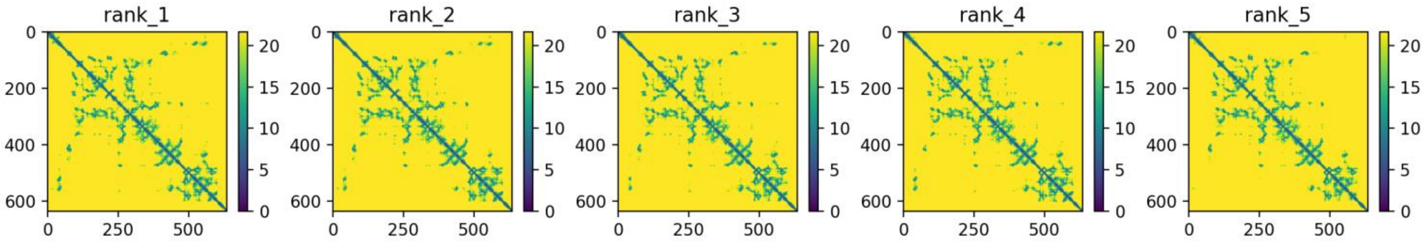


**Legend to Figs. S1-S23.** Images of all models and key quality metrics for the models. The predicted protein folds for all models are displayed in common orientations and color schemes. The color pallet used is: N-terminus, cyan; TP domain, red; T3 motif, violet; Spacer domain, grey; RT1 motif, light blue; RT domain, yellow; RNaseH domain, green; Priming residue, YMDD/YADD/YVDD motifs, D-E-D-D and D-E-E-D motifs, blue spheres; and Aspartate proteinase, magenta. Not all domains/motifs are found in all models. The predicted protein fold is shown the *Predicted Fold* panel. Depth of the multiples sequence alignment used to identify the coordinated variations from which the distance constraints are derived are in the *MSA Coverage* panel. The predicted LDDT value as a function of residue position is in the *Predicted LDDT* panel. Distograms based on the coordinated variations for each of the five models generated for each sequence are found in the *Predicted Distogram* panel.

### Fig. S22. pFOXC3 RT, Isolate AAD38504.1

#### Predicted fold

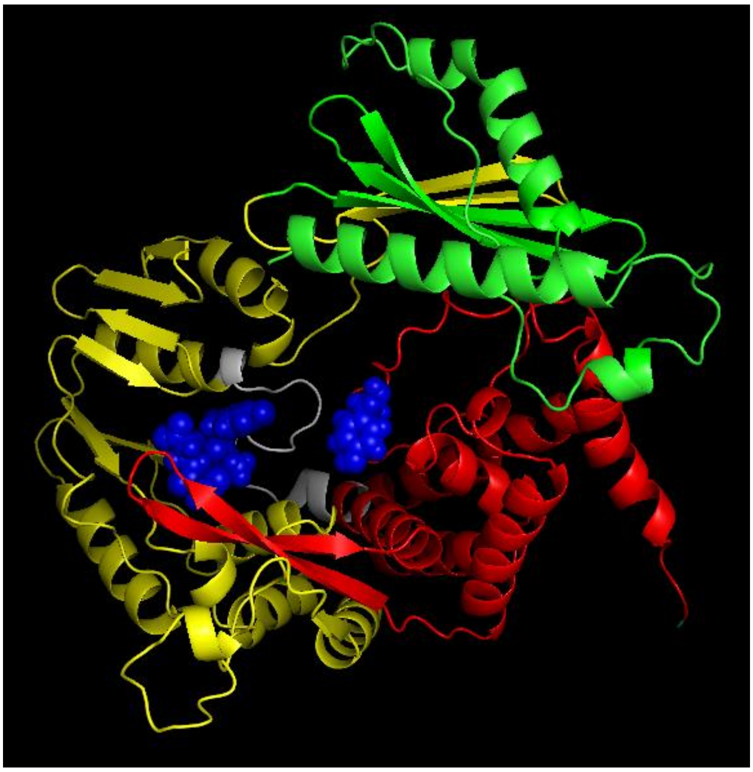

*Domain identification is arbitrarily based on alignment with RNDV P. It is unlikely that the sequences labeled as the TP function the same way as in HBV. The candidate priming residue (Y35) is unconfirmed experimentally. No candidate D-E-D-D motif could be identified.*

N-terminus, cyan; TP, red; Spacer, grey; RT, yellow; RNaseH, green; Y35 priming residue candidate and YADD, blue

#### MSA coverage

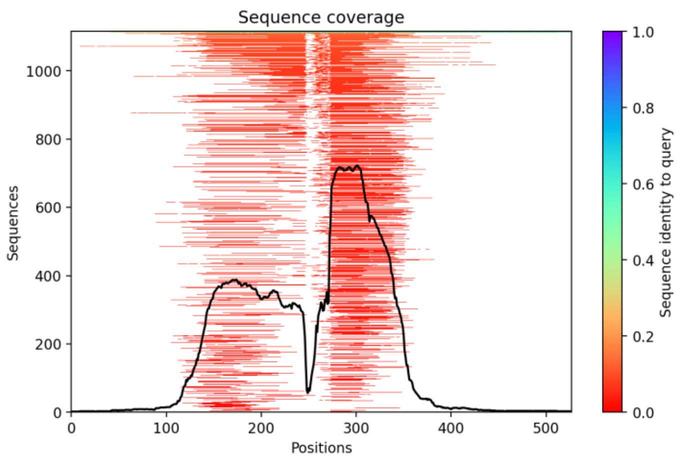


#### Predicted LDDT

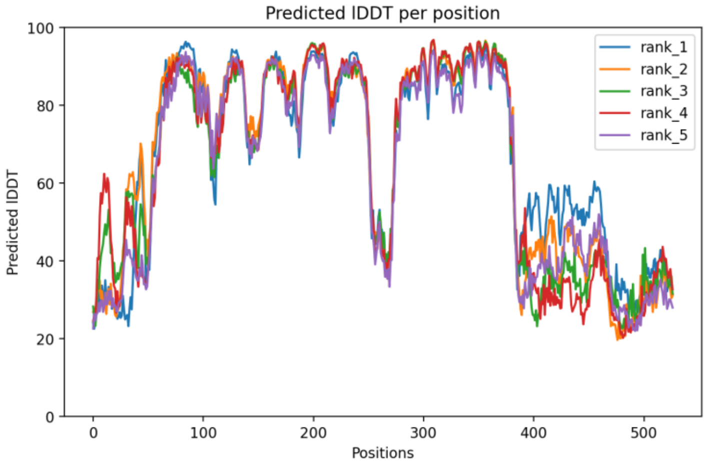


#### Predicted Distogram

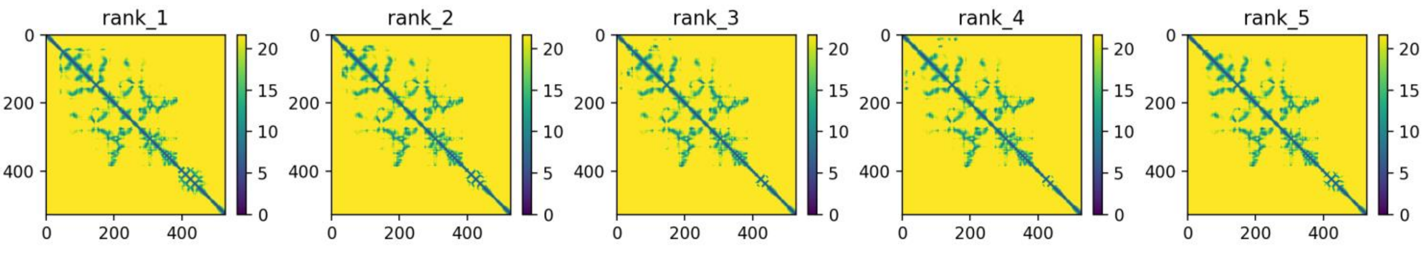


**Legend to Figs. S1-S23.** Images of all models and key quality metrics for the models. The predicted protein folds for all models are displayed in common orientations and color schemes. The color pallet used is: N-terminus, cyan; TP domain, red; T3 motif, violet; Spacer domain, grey; RT1 motif, light blue; RT domain, yellow; RNaseH domain, green; Priming residue, YMDD/YADD/YVDD motifs, D-E-D-D and D-E-E-D motifs, blue spheres; and Aspartate proteinase, magenta. Not all domains/motifs are found in all models. The predicted protein fold is shown the *Predicted Fold* panel. Depth of the multiples sequence alignment used to identify the coordinated variations from which the distance constraints are derived are in the *MSA Coverage* panel. The predicted LDDT value as a function of residue position is in the *Predicted LDDT* panel. Distograms based on the coordinated variations for each of the five models generated for each sequence are found in the *Predicted Distogram* panel.

### Fig. S23. CaMV RT, Isolate M90542.1

#### Predicted fold

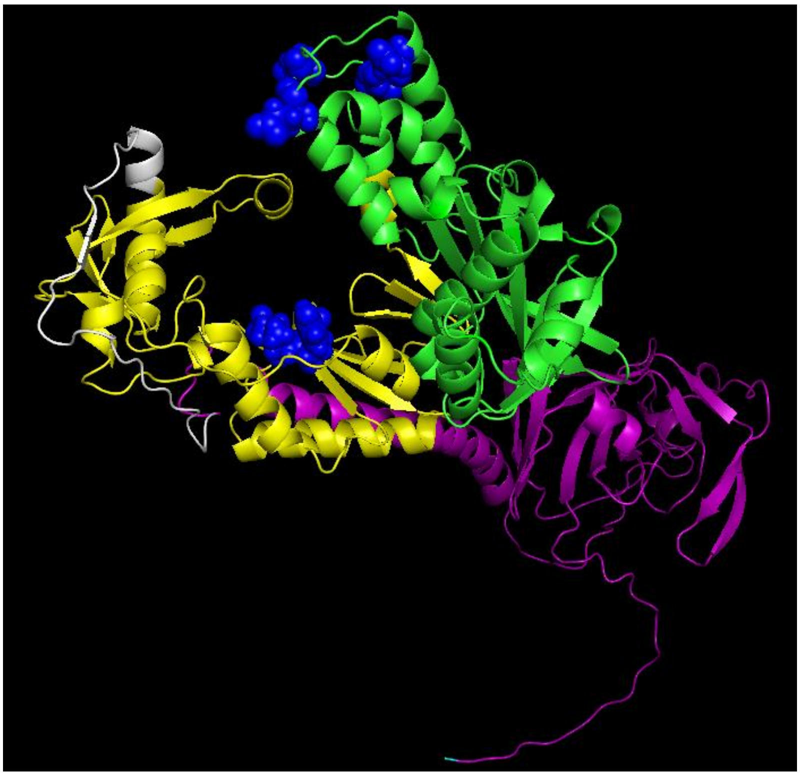

*Domain boundaries are arbitrary and based on alignment with RNDV P. DEDD candidate residues are unconfirmed experimentally.*

N-terminus, cyan; Aspartate proteinase, magenta; Spacer, grey; RT, yellow; RNaseH, green; YVDD and D-E-E-D motif, blue

#### MSA coverage

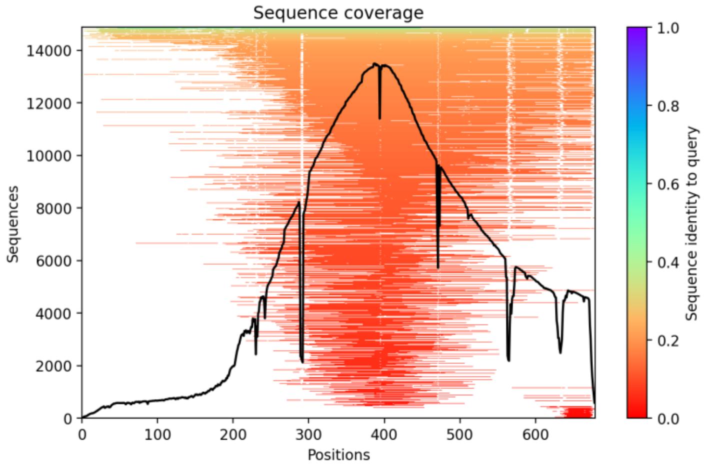


#### Predicted LDDT

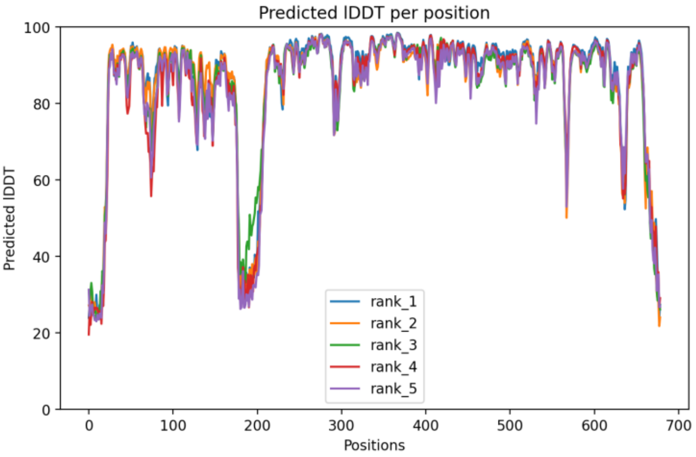


#### Predicted Distogram

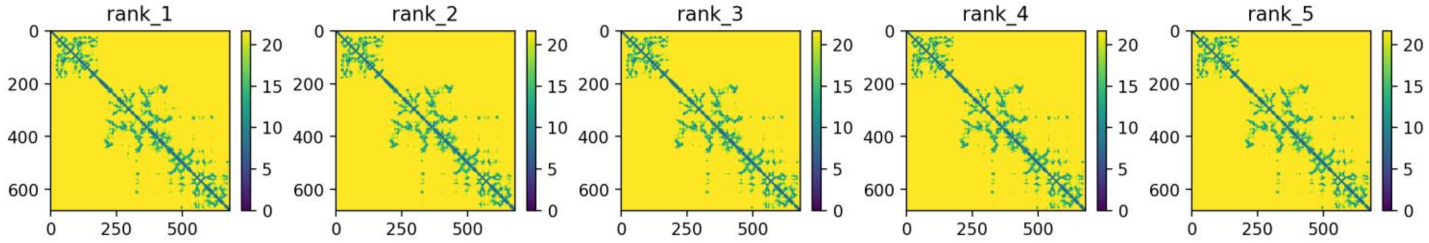


**Legend to Figs. S1-S23.** Images of all models and key quality metrics for the models. The predicted protein folds for all models are displayed in common orientations and color schemes. The color pallet used is: N-terminus, cyan; TP domain, red; T3 motif, violet; Spacer domain, grey; RT1 motif, light blue; RT domain, yellow; RNaseH domain, green; Priming residue, YMDD/YADD/YVDD motifs, D-E-D-D and D-E-E-D motifs, blue spheres; and Aspartate proteinase, magenta. Not all domains/motifs are found in all models. The predicted protein fold is shown the *Predicted Fold* panel. Depth of the multiples sequence alignment used to identify the coordinated variations from which the distance constraints are derived are in the *MSA Coverage* panel. The predicted LDDT value as a function of residue position is in the *Predicted LDDT* panel. Distograms based on the coordinated variations for each of the five models generated for each sequence are found in the *Predicted Distogram* panel.

### Fig. 24. HBV P catalytic core (RT-RNase H) alignments to HBV domain models and non-HBV enzymes

A

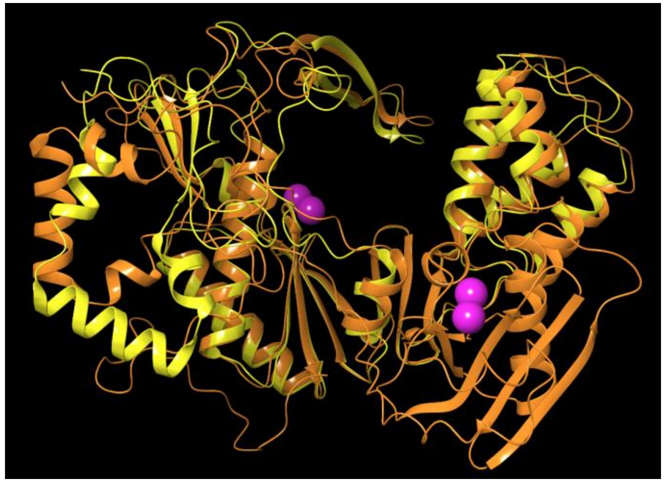

gtB RT-RNaseH (orange) to gtB RT (yellow), RMSD=2.88 Å

B

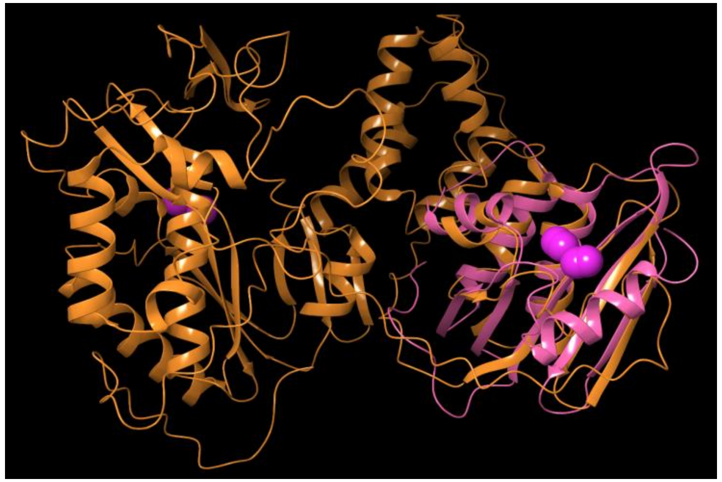

gtB RT-RNaseH to HIV RH (pink), RMSD=2.84 Å

C

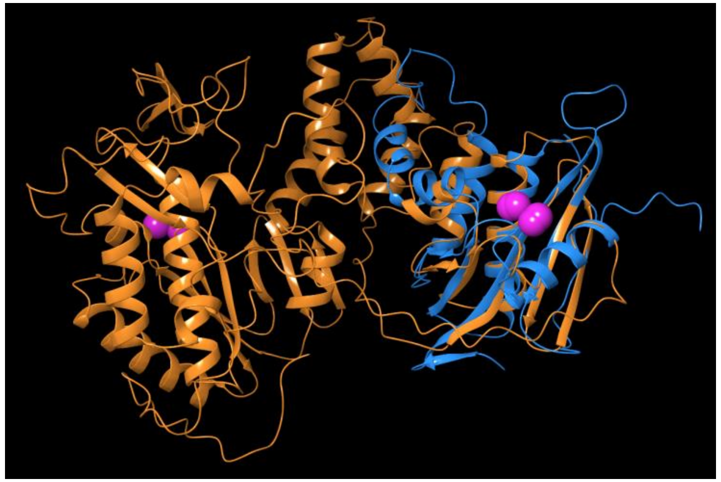

gtB RT-RNaseH to human RNaseH1 (azure), RMSD=2.66 Å

D

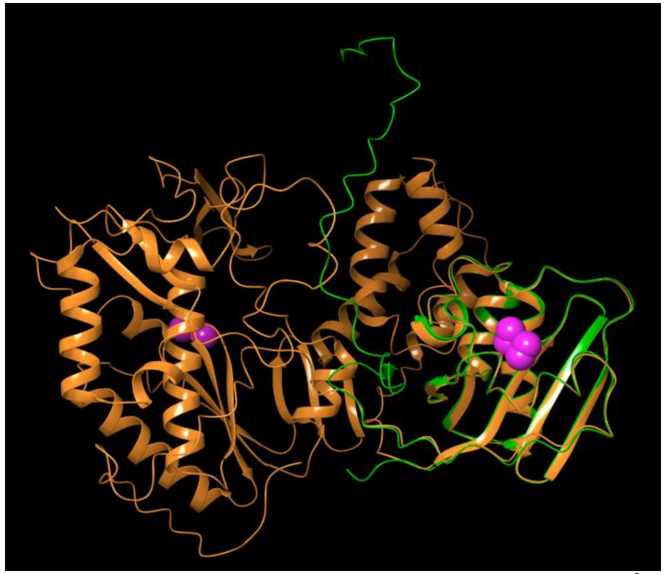

gtB RT-RNaseH to HBV RH (green), RMSD=1.38 Å

E

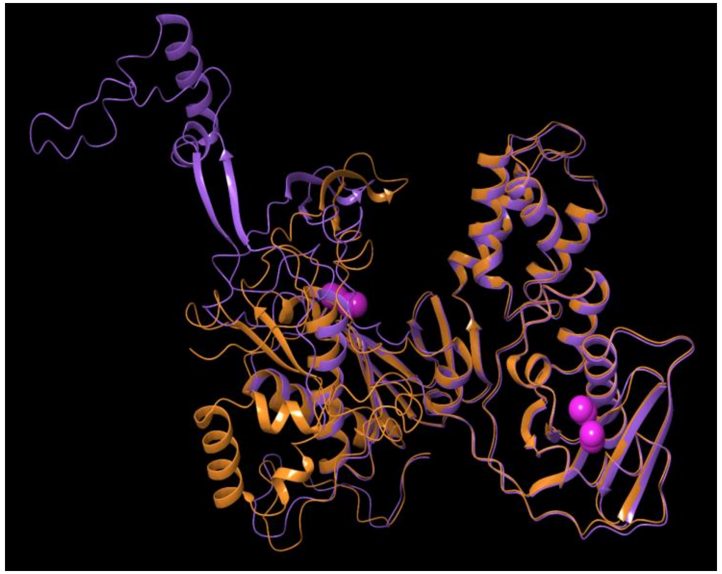

gtB RT-RNaseH to gtD RT-RNaseH (faded violet), RMSD=2.15 Å

F

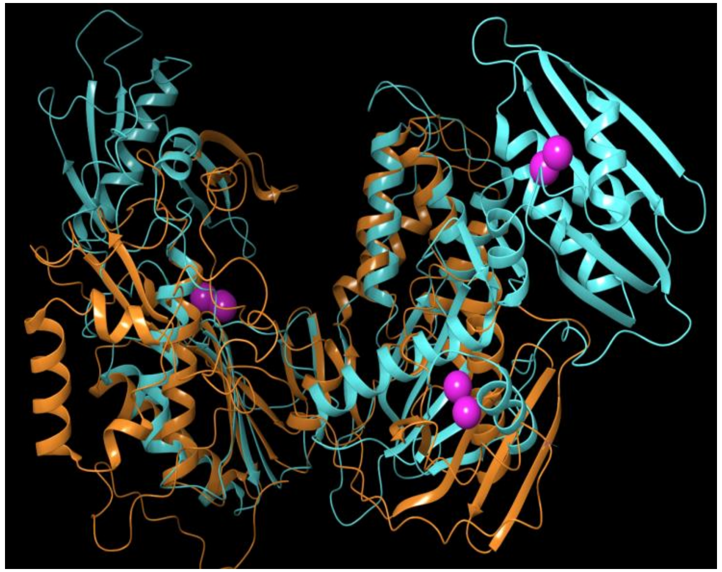

gtB RT-RNaseH truncated to HIV RT-RNase H (cyan), RMSD=3.75 Å

**Legend to Fig. S24.** Superpositions of the HBV P catalytic core (RT-RNase H) to HBV domain models and non-HBV enzymes. Models were superposed and RMSD values were calculated. Models compared and the colors used to represent them are indicted below the images. RMSD values are below the images. gt, genotype.

#### Fig. S24 (*Continued*). HBV P catalytic core (RT-RNase H) alignments to HBV domain models and non-HBV enzymes

G

H

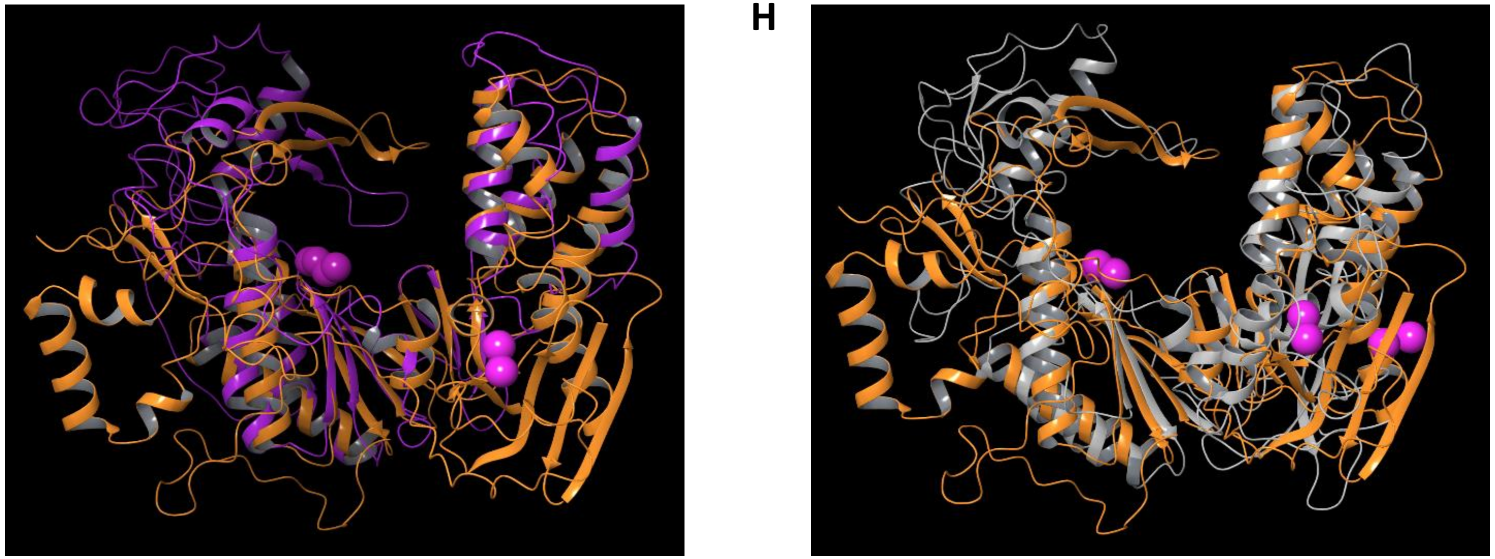

gtB RT-RNaseH to Das RT (violet), RMSD=3.38 Å

gtB RT-RNaseH to Ty3 RT (faded gray), RMSD=3.6 Å

**Legend to Fig. S24.** Superpositions of the HBV P catalytic core (RT-RNase H) to HBV domain models and non-HBV enzymes. Models were superposed and RMSD values were calculated. Models compared and the colors used to represent them are indicted below the images. RMSD values are below the images. gt, genotype.

#### Fig. S25. HBV genotype B P alignments to other HBV P genotypes

A

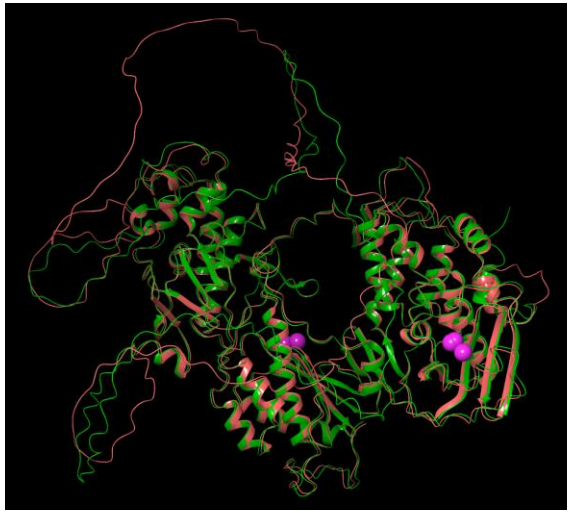

HBV P gtB (green) to gtA (red-orange), RMSD=1.79 Å

B

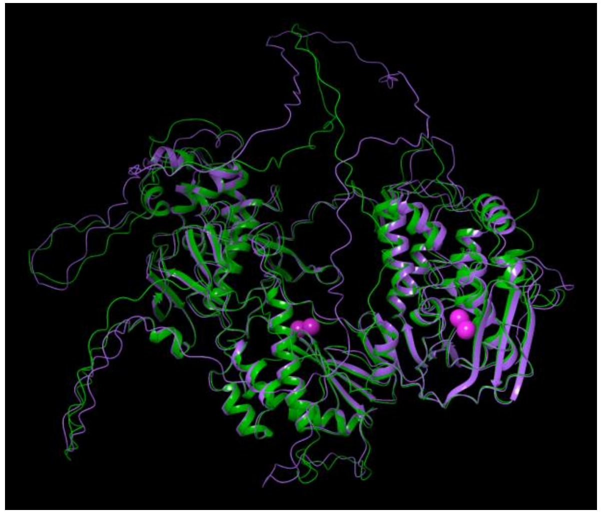

HBV P gtB to gtC (faded violet), RMSD=2.07 Å

C

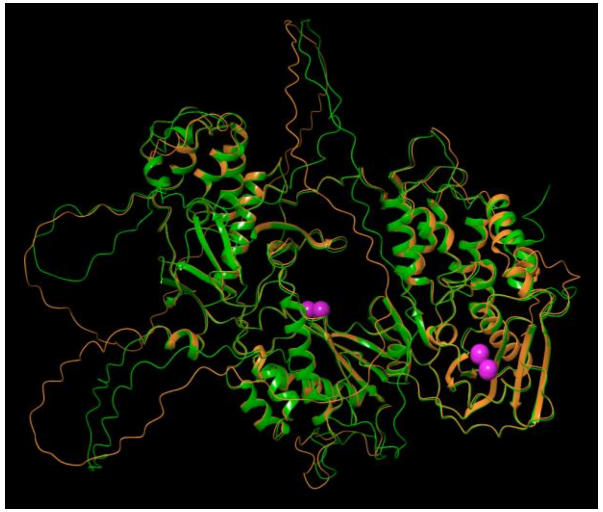

HBV P gtB to gtD (orange), RMSD=1.96 Å

D

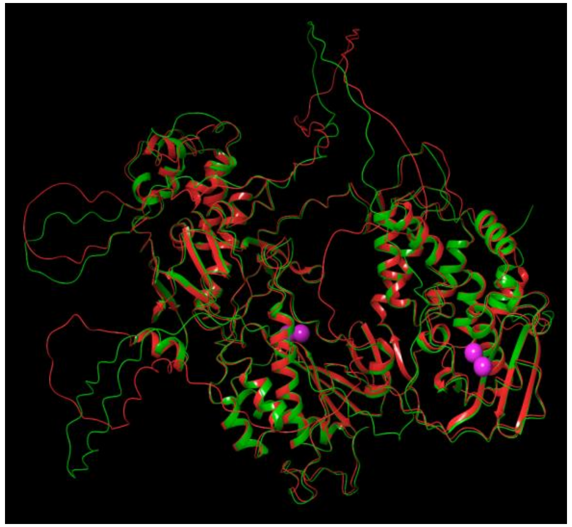

HBV P gtB to gtE (red), RMSD=2.16 Å

E

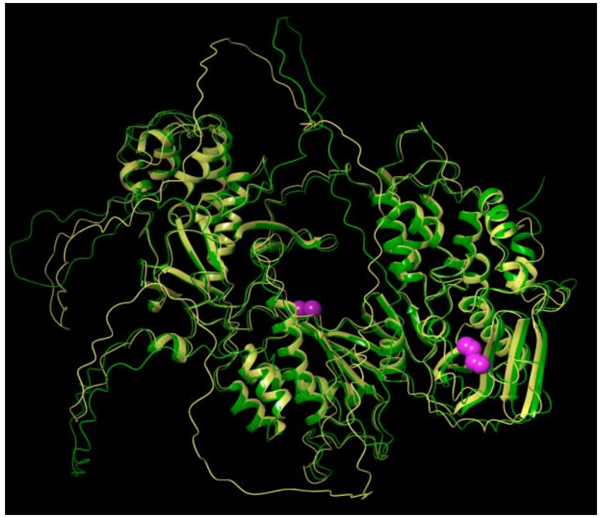

HBV P gtB to gtF (yellow), RMSD=2.16 Å

F

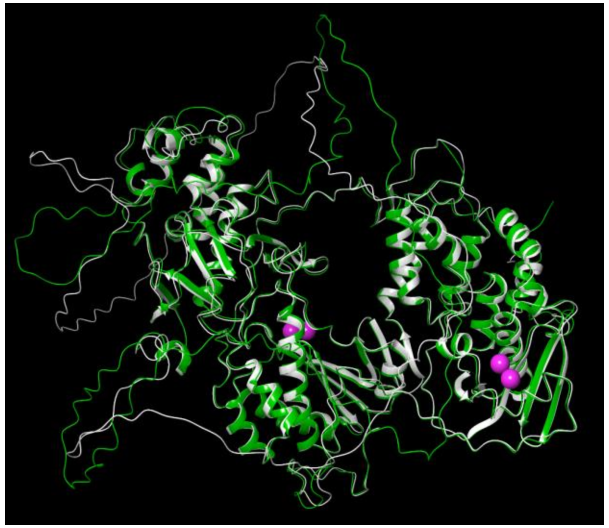

HBV P gtB to gtG (platinum), RMSD=1.94 Å

**Legend to Fig. S25.** Superpositions of HBV genotype B P models to other HBV P genotypes. Models were superposed and RMSD values were calculated. Models compared and the colors used to represent them are indicted below the images. RMSD values are below the images. gt, genotype.

### Fig. S25 (*Continued*). HBV genotype B P alignments to other HBV P genotypes

**G**

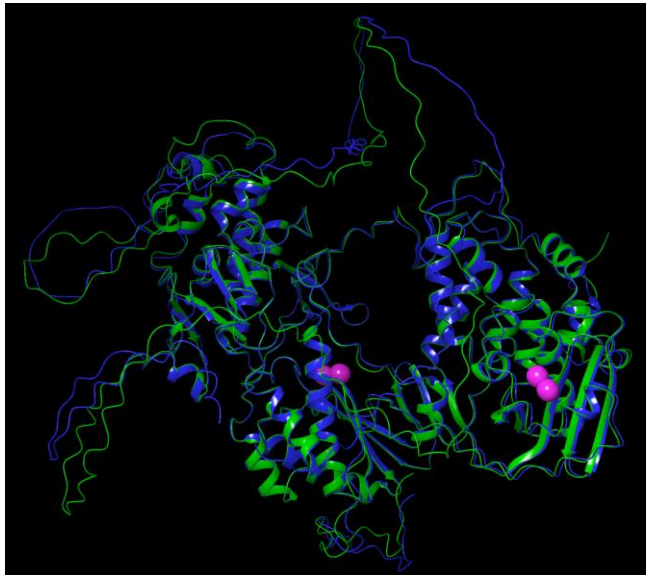

HBV P gtB to gtH (blue), RMSD=2.12 Å

**H**

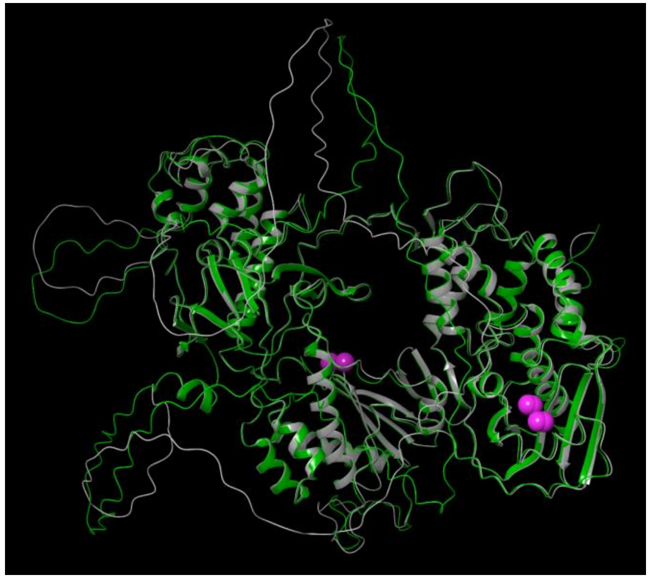

HBV P gtB to gtI (gray), RMSD=1.67 Å

**Legend to Fig. S25.** Superpositions of HBV genotype B P models to other HBV P genotypes. Models were superposed and RMSD values were calculated. Models compared and the colors used to represent them are indicted below the images. RMSD values are below the images. gt, genotype.

#### Fig. S26. HBV genotype B P alignments to isolated HBV P domain models

A

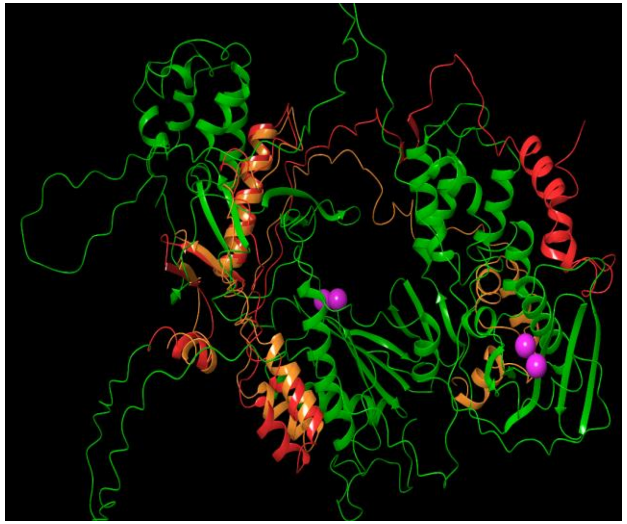

HBV P gtB TP (red) to isolated TP (orange) RMSD=3.07 Å

B

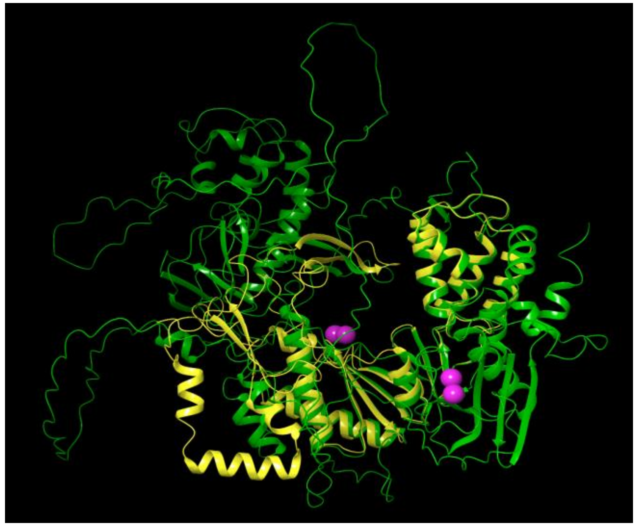

HBV P gtB to RT (yellow) RMSD=2.4 Å

C

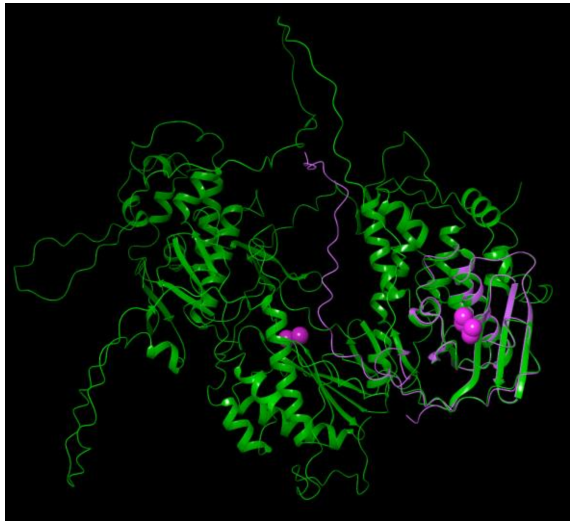

HBV P gtB to RNaseH (faded violet), RMSD=1.21 Å

D

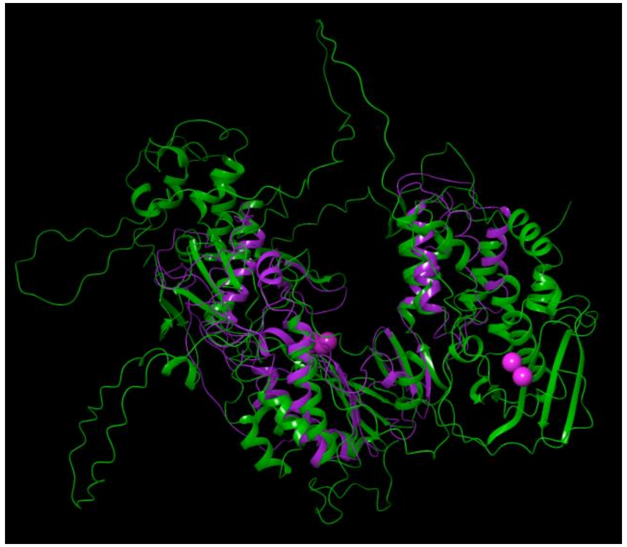

HBV P gtB to Das et al. RT (violet), RMSD=3.24 Å

E

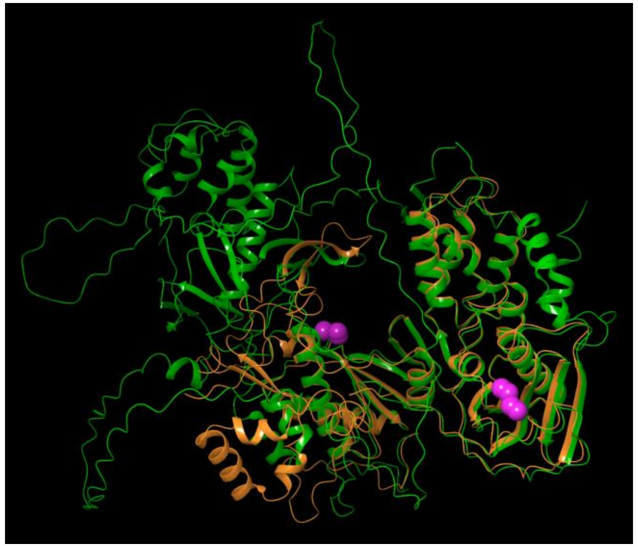

HBV P gtB to gtB RT-RNaseH (orange), RMSD=2.37 Å

**Legend to Fig. S26.** HBV genotype B P superpositions with isolated HBV P domain models. Models were superposed and RMSD values were calculated. Models compared and the colors used to represent them are indicted below the images. RMSD values are below the images. gt, genotype.

#### Fig. S27. HBV P and domain alignments to non-hepadnaviral enzymes

A

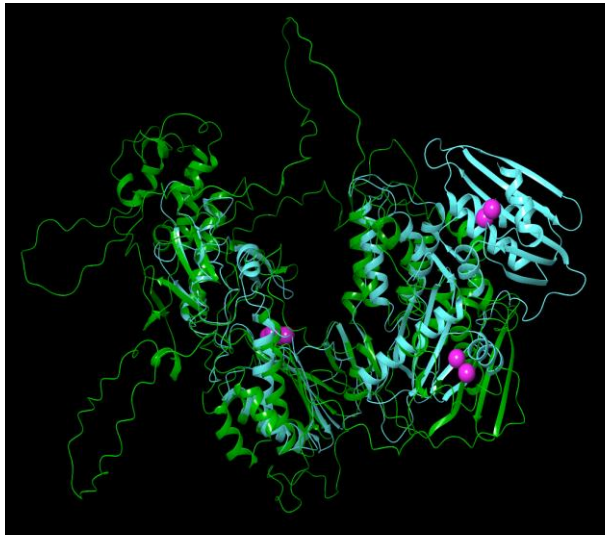

HBV P gtB (green) to HIV RT-RNaseH (cyan), RMSD=3.42 Å

B

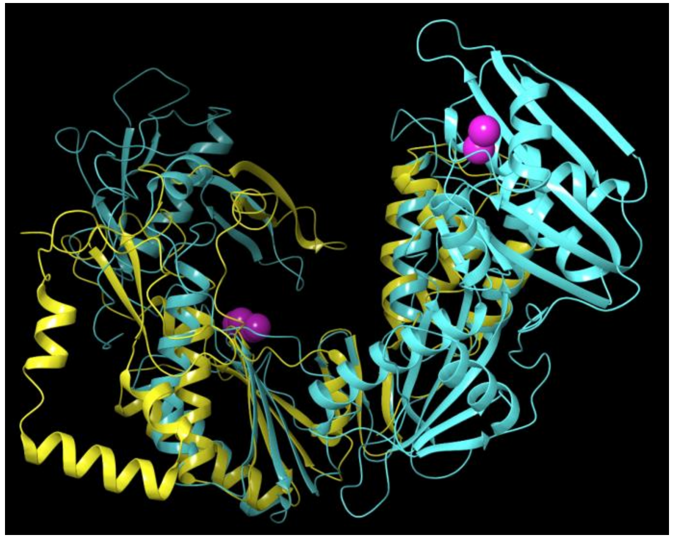

HBV RT gtB (yellow) to HIV RT-RNaseH (cyan), RMSD=3.42 Å

C

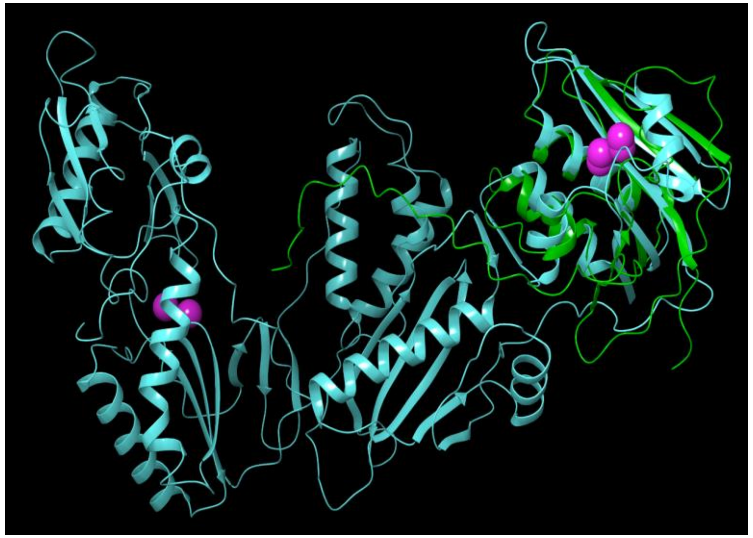

HBV RH gtB (green) to HIV RT-RNaseH (cyan), RMSD=2.36 Å

D

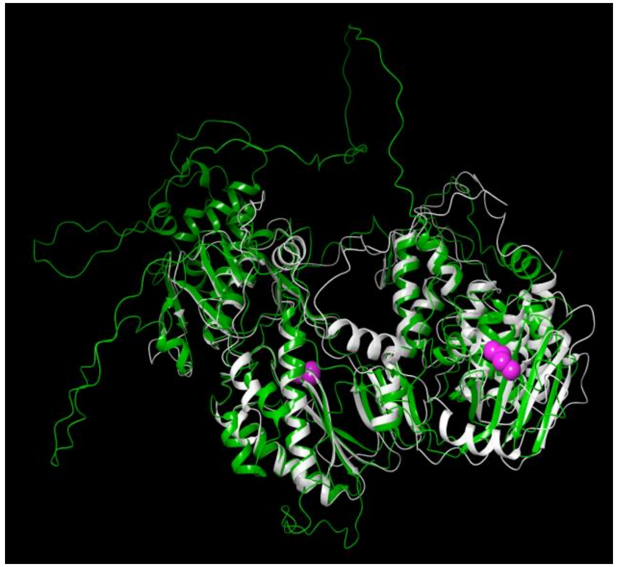

HBV P gtB to RNDV P (white), RMSD=2.76 Å

E

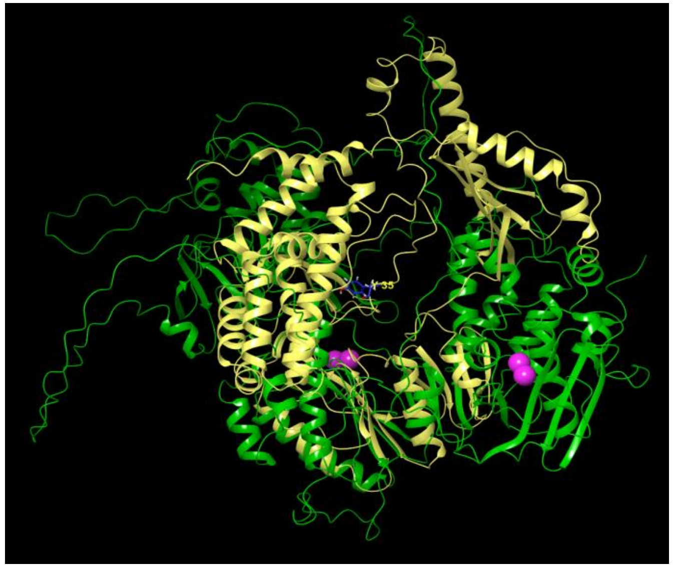

HBV P gtB to pFOXC3 RT (faded yellow), RMSD=4.52 Å

F

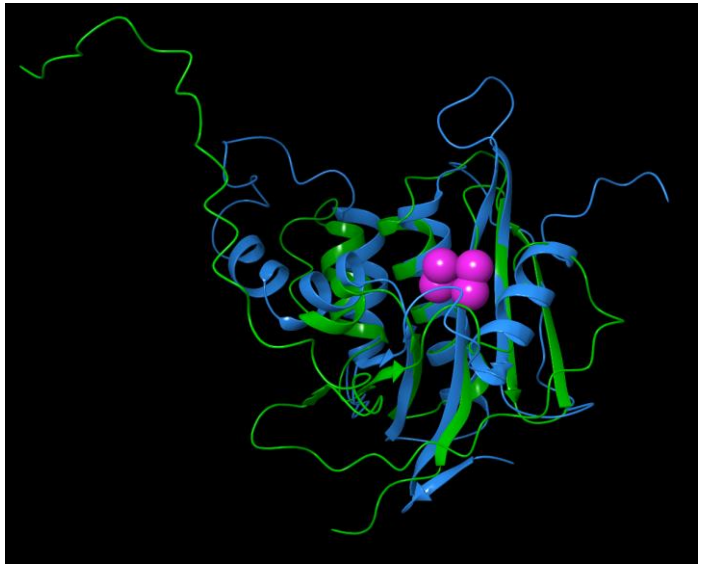

HBV RNaseH gtB (green) to human RNase H1 (azure), RMSD=2.76 Å

**Legend to Fig. S27.** HBV P and domain models superpositions with non-hepadnaviral enzymes. Models were superposed and RMSD values were calculated. Models compared and the colors used to represent them are indicted below the images. RMSD values are below the images. gt, genotype.

**Fig. S27 (*Continued*). HBV P and domain alignments to non-hepadnaviral enzymes**

G

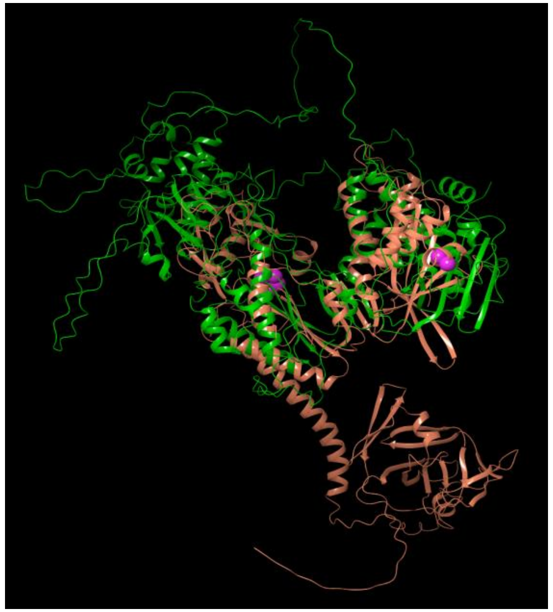

HBV P gtB to CaMV RT (faded orange), RMSD=3.96 Å

H

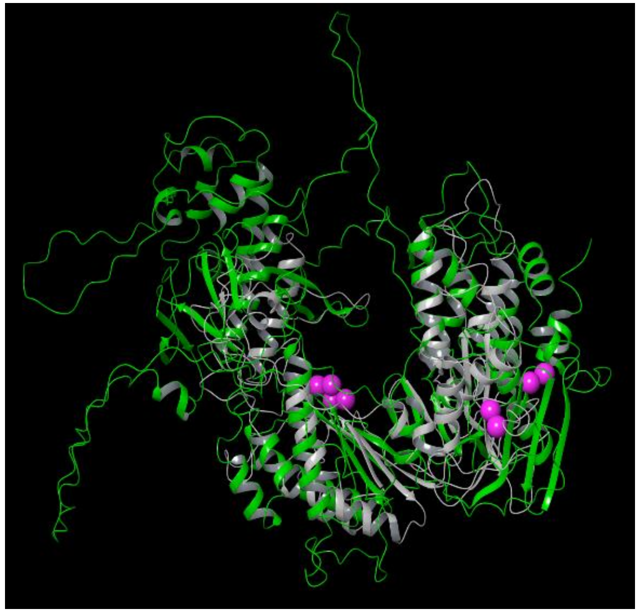

HBV P gtB to Ty3 RT (gray), RMSD=3.34 Å

**Legend to Fig. S27.** HBV P and domain models superpositions with non-hepadnaviral enzymes. Models were superposed and RMSD values were calculated. Models compared and the colors used to represent them are indicted below the images. RMSD values are below the images. gt, genotype.

Fig. S28. Docking of RNA:DNA heteroduplexes into models for HBV P and the catalytic core of P

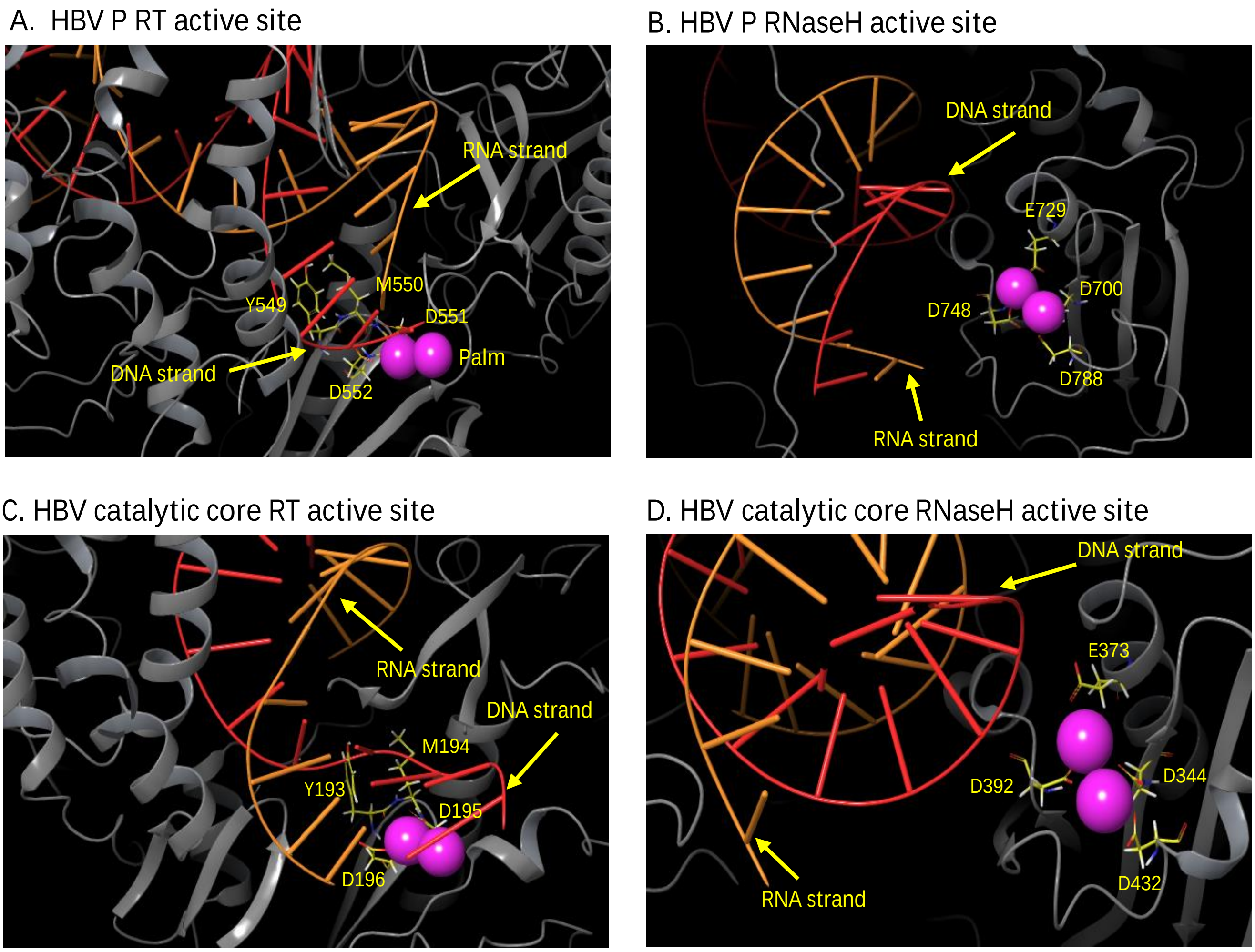


Fig. S28. Docking of RNA:DNA heteroduplexes into models for HBV P and the catalytic core of P. A. RT active site of the HBV P genotype B model. B. RNaseH active site of the HBV P genotype B model. C. RT active site of the genotype B HBV RT-RNaseH catalytic core model. D. RNaseH active site of the genotype B HBV RT-RNaseH catalytic core model. E. Heteroduplex docked to the HBV genotype D RT-RNaseH catalytic core model. F. RT active site of the genotype D HBV RT-RNaseH catalytic core model. G. RNaseH active site of the genotype D HBV RT-RNaseH catalytic core model. The YMDD and D-E-D-D residues are shown as sticks and are labeled. Red, DNA strand; Orange, RNA strand; Magenta spheres, $Mg^{++}$ ions.

Fig. S28 (*continued*). Docking of RNA:DNA heteroduplexes into models for HBV P and the catalytic core of P

E. HBV genotype D catalytic core

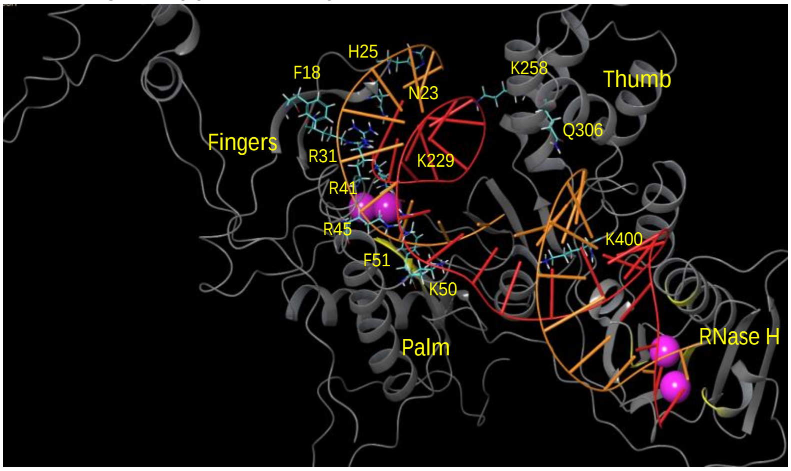


F. Genotype B RT active site

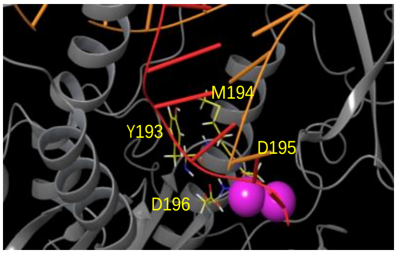


G. Genotype D RNaseH active site

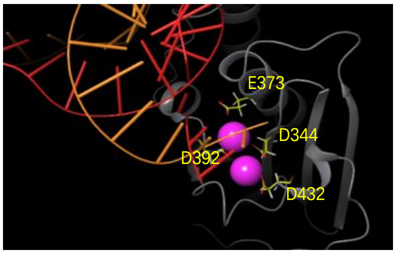


Fig. S28. Docking of RNA:DNA heteroduplexes into models for HBV P and the catalytic core of P . A. RT active site of the HBV P genotype B model. B. RNaseH active site of the HBV P genotype B model. C. RT active site of the genotype B HBV RT-RNaseH catalytic core model. D. RNaseH active site of the genotype B HBV RT-RNaseH catalytic core model. E. Heteroduplex docked to the HBV genotype D RT-RNaseH catalytic core model. F. RT active site of the genotype D HBV RT-RNaseH catalytic core model. G. RNaseH active site of the genotype D HBV RT-RNaseH catalytic core model. The YMDD and D-E-D-D residues are shown as sticks and are labeled. Red, DNA strand; Orange, RNA strand; Magenta spheres, $Mg^{++}$ ions.

Fig. S29. Docking poses for the **ε** RNA to the HBV P genotype B model

A. **ε** docking pose 1

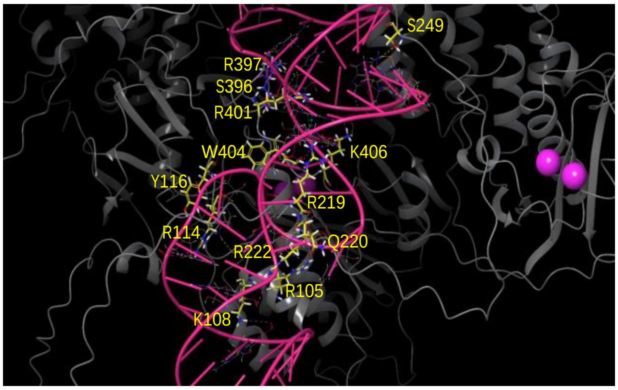


B. **ε** docking pose 2

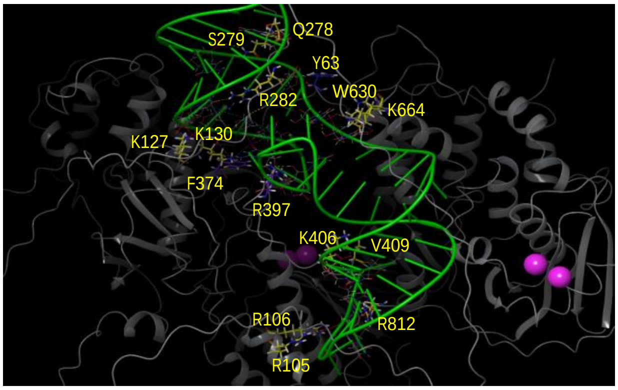


C. **ε** docking pose 3

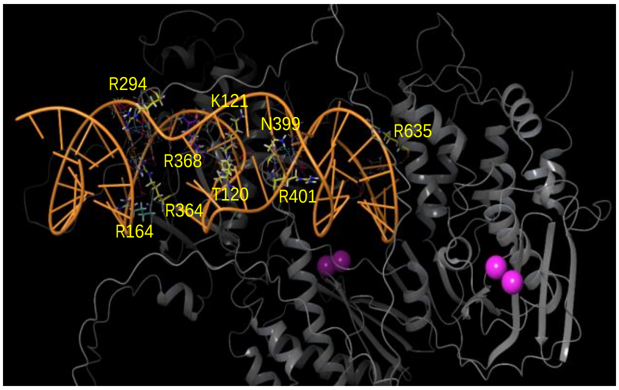


Fig. S29. Docking poses for the **ε** RNA to the HBV P genotype B model. A.-C. show representatives for the the three classes of docking poses for ε in the HBV P model. Residues interacting with ε are indicated in stick representation and are labeled. Cyan, residues in the T3 motif; Violet, residues in the RT1 motif; Magenta spheres, $Mg^{++}$ ions.

**Fig. S30. Comparisons of interaction networks for the wild-type and mutant residues used to validate the HBV P model**

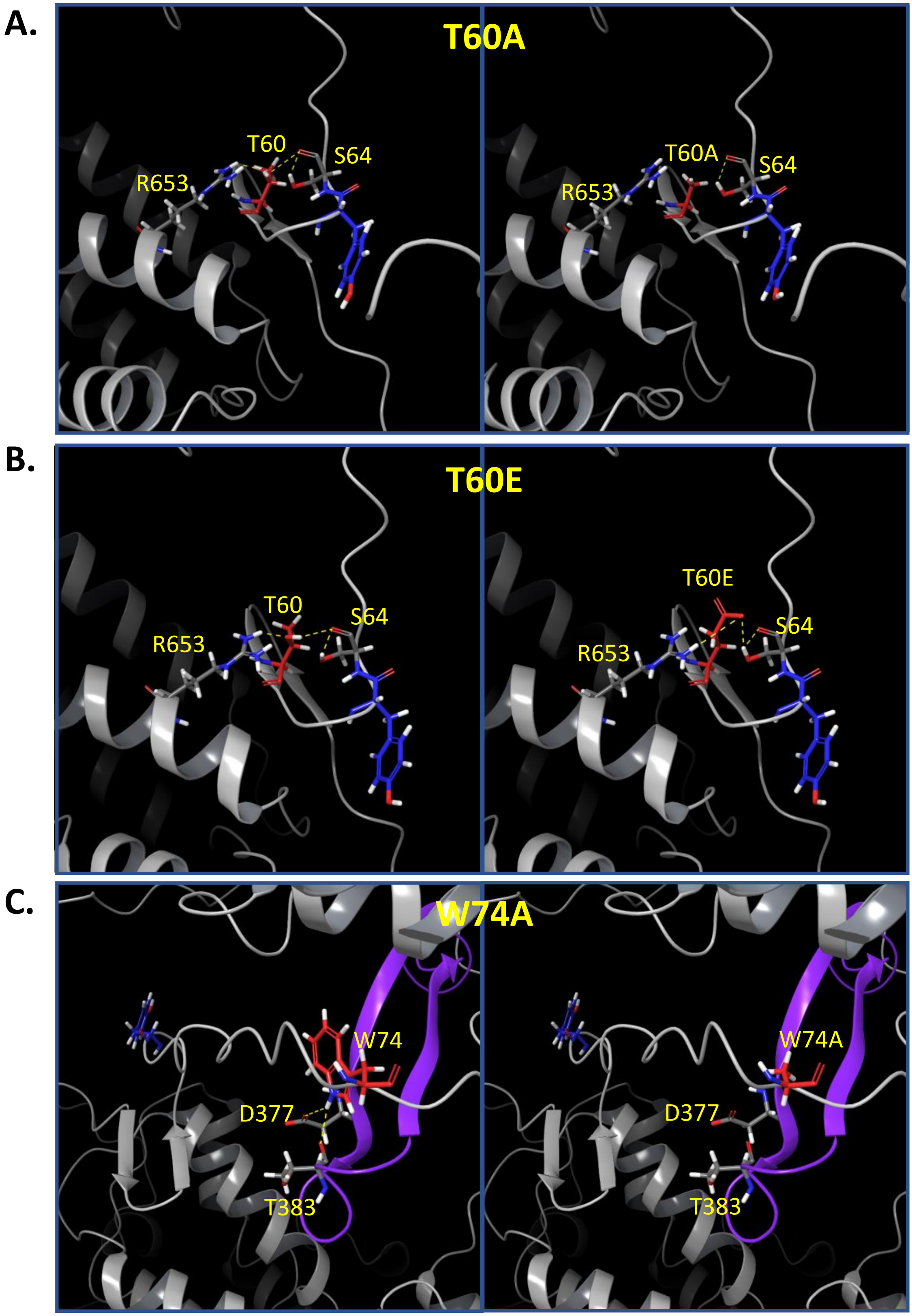

**Fig. S30. Comparisons of interaction networks for the wild-type and mutant residues used to validate the HBV P model**

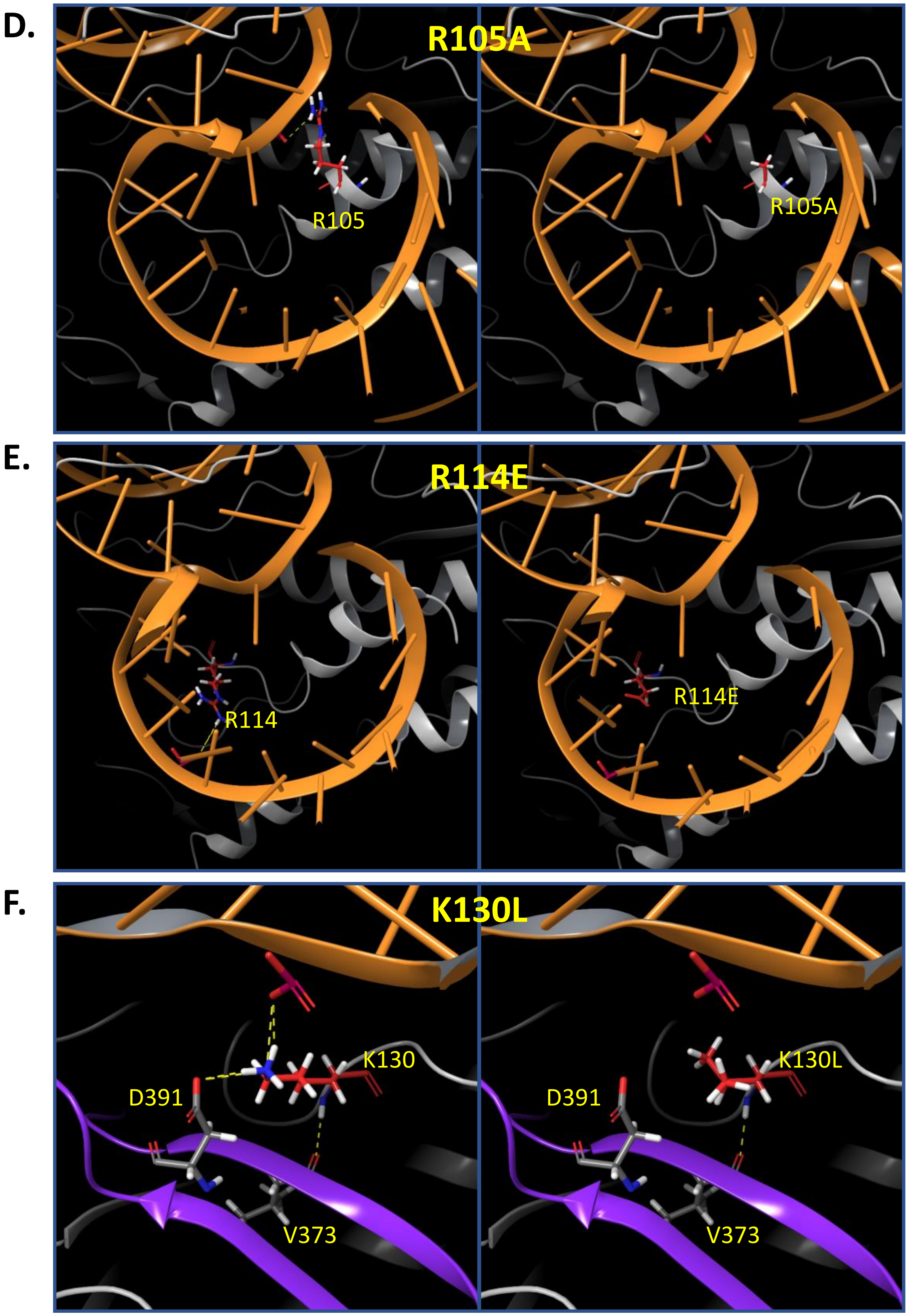

**Fig. S30. Comparisons of interaction networks for the wild-type and mutant residues used to validate the HBV P model**

G.

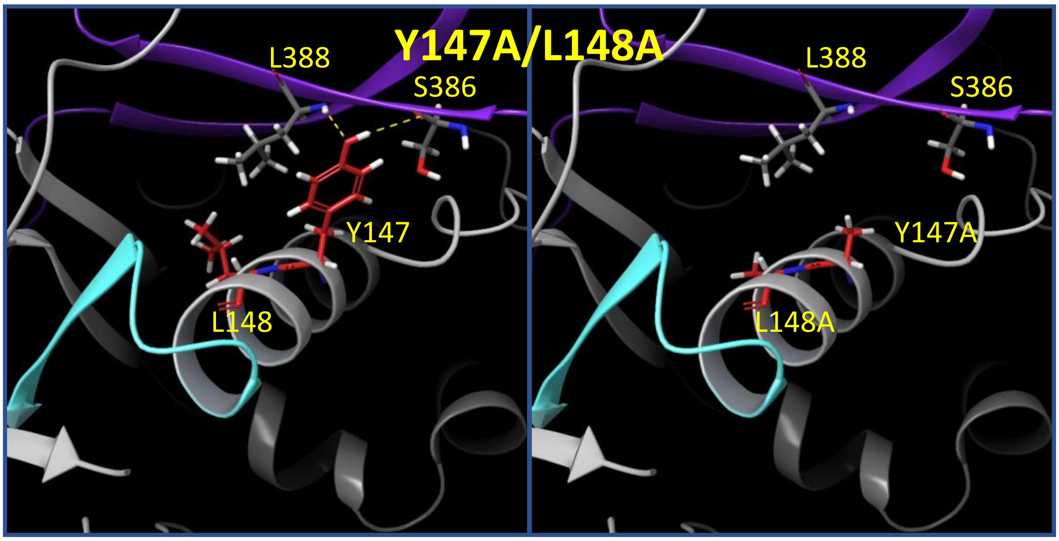


H.

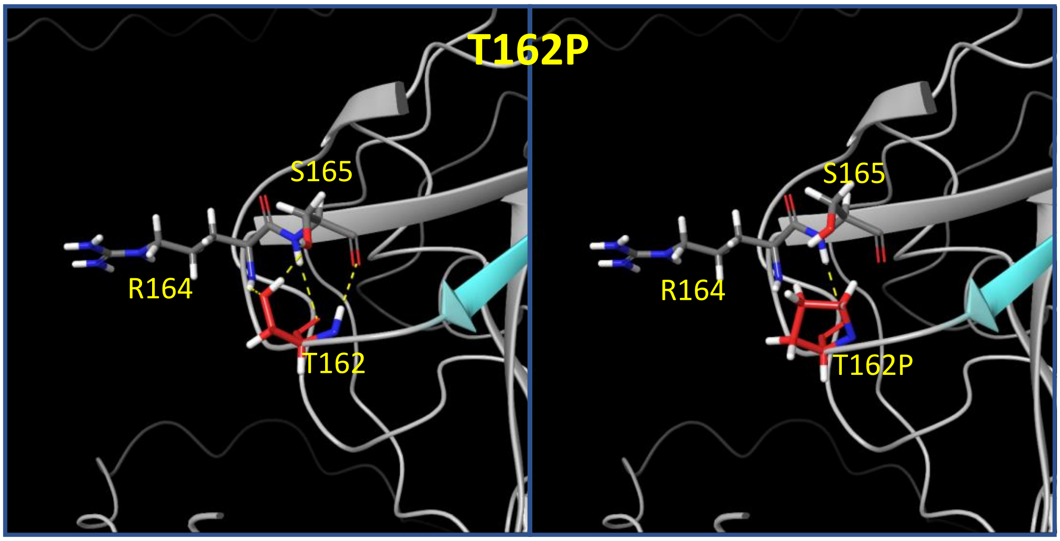


I.

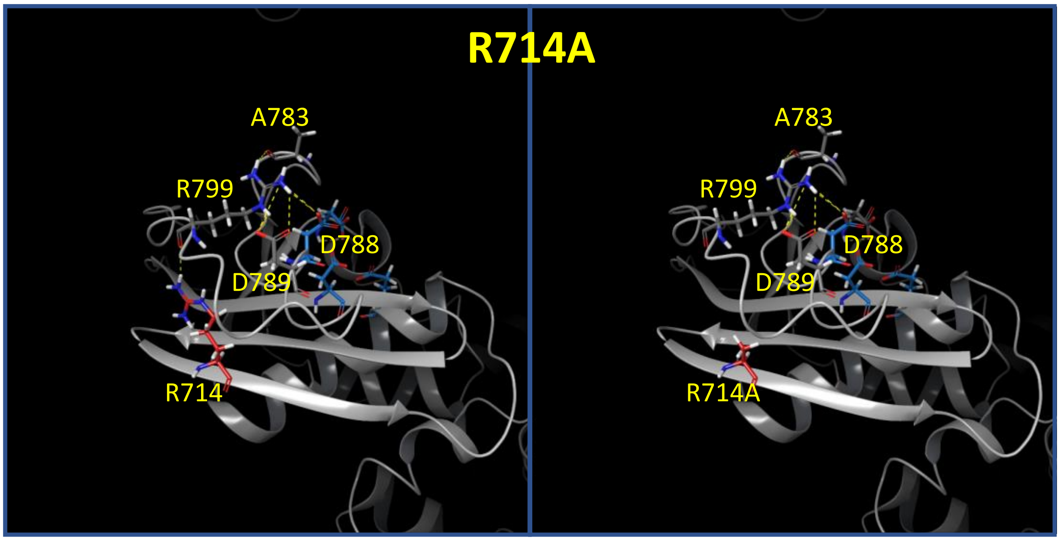

**Fig. S30. Comparisons of interaction networks for the wild-type and mutant residues used to validate the HBV P model. A. – I.** Residues and their key interactions used in validating the HBV genotype B model for the full-length P. *Left Panels*: WT residues; *Right Panels*: Mutant residues. Red, residues mutated; , Orange, ε RNA; Purple, RT1 motif; Cyan, T3 motif; Blue, Y63; Magenta, YMDD motif; Light Blue, D-E-D-D motif; Dashed lines, Polar contacts.
